# stCNASim: Allele-aware spatial RNA-seq simulator enables systematic benchmarking of copy number inference

**DOI:** 10.64898/2026.08.06.743179

**Authors:** Xianjie Huang, Rongting Huang, Jiamu Qiao, Yuanhua Huang

## Abstract

Spatial transcriptomics (ST) is revolutionizing the study of tumor evolution by enabling spatially resolved copy-number alteration (CNA) analysis. However, evaluating the accuracy and robustness of current single-cell (SC) and ST-specific CNA inference tools remains challenging due to the absence of ground-truth datasets. Here, we present stCNASim, an allele-aware spatial RNA-seq simulator that generates raw reads within realistic spatial contexts. We synthesized 46 benchmarking datasets across varying technical settings and spatial architectures to evaluate five widely used computational methods. Our analysis reveals that while SC-based methods adapt well to ST data, ST-specific methods successfully benefit from considering spatial autocorrelation but struggle under high spatial intermixing. The allele-aware methods CalicoST, Numbat, and XClone achieved top-tier performance with unique advantages in extreme scenarios, yet showed distinct sensitivities to low purity, mirrored alleles, and low coverage, respectively. By providing a scalable simulator and a rigorous benchmark, this work establishes a much-needed framework to guide and accelerate future tool development in spatial CNA analysis.

## 1 Introduction

Intra-tumor heterogeneity driven by subclonal evolution poses a formidable challenge to understanding cancer progression and therapeutic resistance. Somatic copy number alterations (CNAs, interchangeably known as copy number variations, CNVs)—encompassing copy gains, losses, and loss of heterozygosity (LOH)—are fundamental drivers of this heterogeneity across diverse cancer types. By profoundly altering gene expression and driving phenotypic plasticity, CNAs dictate tumor evolution and shape the genomic landscape of cancer.

Initially, the detection of CNAs has relied on bulk whole-genome or exome sequencing, which has illuminated broad CNA landscapes across pan-cancer cohorts [1, 2]. Subsequently, single-cell sequencing revolutionized the field by enabling the resolution of CNA-driven clonal architectures at single-cell resolution with RNA alone or paired DNA [3, 4]. More recently, spatial transcriptomics (ST) has introduced a crucial new dimension: preserving the native histological context. ST enables not only the mapping of subclonal spatial distributions but also the investigation of their colocalization and interactions with the tumor microenvironment (TME), including immune and stromal compartments. Prominent examples of spatial CNA subclones include prostate cancer [5], ovarian cancer [6], metastatic pancreatic cancer [7], and across multiple cancer types in the HTAN studies [8].

However, exploiting ST for mutation detection is hindered by biased genomic coverage and high transcript dropout rates. Consequently, large-scale CNAs are currently the major class of somatic mutations that can be reliably inferred from ST data. Furthermore, the capacity to detect specific CNA modalities depends heavily on the underlying ST chemistry (Supplementary Data 1). Poly-A-capture-based spatial RNA-seq (e.g., 10x Visium v1, Stereo-seq) captures transcribed single-nucleotide polymorphisms (SNPs), offering applicability similar to scRNA-seq for resolving allele-specific CNAs. In contrast, probe-based spatial profiling or imaging technologies (e.g., Visium CytAssist, Visium HD) typically lack SNP-level resolution, restricting their utility to the detection of total copy number gains and losses (allele-agnostic CNAs).

Given the inherent sparsity and noise of spatial data, computational methods are indispensable for robust CNA inference. A suite of algorithms originally designed for scRNA-seq is routinely applied to ST data by treating spatial coordinates as independent cellular or multi-cellular samples, such as InferCNV [3] and CopyKAT [9] for allele-agnostic detection, alongside CaSpER [10], Numbat [11] and XClone [12] for allele-aware resolution. More recently, ST-specific tools are also emerging. For example, SlideCNA [13] and STmut [14] are derived from the existing tools InferCNV and CNVKit by including an interface for handling ST data. With more tailored designs, STARCH [15], its successor CalicoST [16], and SpaCNA [17] all leverage spatial autocorrelation via Markov random fields to enhance CNA analysis, among which Cali-coST has demonstrated impressive potential in detecting allele-specific CNAs. Despite this methodological proliferation, a critical gap remains: it is entirely unclear how reliably these tools perform across diverse ST platforms, CNA configurations, and tissue architectures. Existing methods are typically evaluated on limited datasets lacking definitive ground truth, leaving the field without a clear consensus on their accuracy, sensitivity, and platform suitability.

While CNA analysis from single-cell RNA-seq has been benchmarked against dozens of experimental datasets [18, 19], these datasets generally suffer from limited ground truth and/or unmatched cells in the DNA-seq. Instead, systematic benchmarking of ST-based CNA detection requires robust, ground-truth-aware datasets, ideally from a reliable simulator. However, current ST simulators, e.g., scDesign2 [20] and scDesign3 [21], predominantly operate at the count-matrix level, rendering them fundamentally incapable of simulating the read-level allelic variation needed to evaluate allele-specific CNA detection. While scReadSim can generate reads from a reference genome [22], it lacks the allelic information and hence cannot evaluate allele-specific CNA. To address this, we developed stCNAsim, a highly versatile, read-level simulator, explicitly engineered to model spatial CNAs across diverse ST chemistries. Utilizing stCNAsim, we generated 46 synthetic spatial datasets encompassing a wide parameter space of CNA types, tumor purities, sequencing depths, and spatial structures. Using this ground-truth compendium, we comprehensively benchmarked current single-cell- and spatial-based CNA inference algorithms. This study provides a foundational evaluation of existing tools, illuminating their respective strengths and methodological blind spots, and establishes a rigorous framework to guide the accurate spatial mapping of subclonal genomic complexity.

## 2 Results

### 2.1 Overview of the allele-aware spatial RNA-seq simulation and benchmarking framework

To enable systematic evaluation of copy number alteration detection from spatial transcriptomics data, we developed an allele-specific simulator, **stCNASim**, and coupled it with a comprehensive benchmarking design spanning various simulated datasets. The overall workflow is summarized in Fig. 1a,b. The simulator was designed to generate spatial RNA-seq datasets with controlled CNA ground truth while preserving realistic expression and allelic characteristics from empirical data. This allows direct assessment of CNA inference methods across multiple biologically relevant tasks, including CNA prediction, tumor identification, subclone inference, and robustness to spatial pattern variation.

**Fig. 1.**
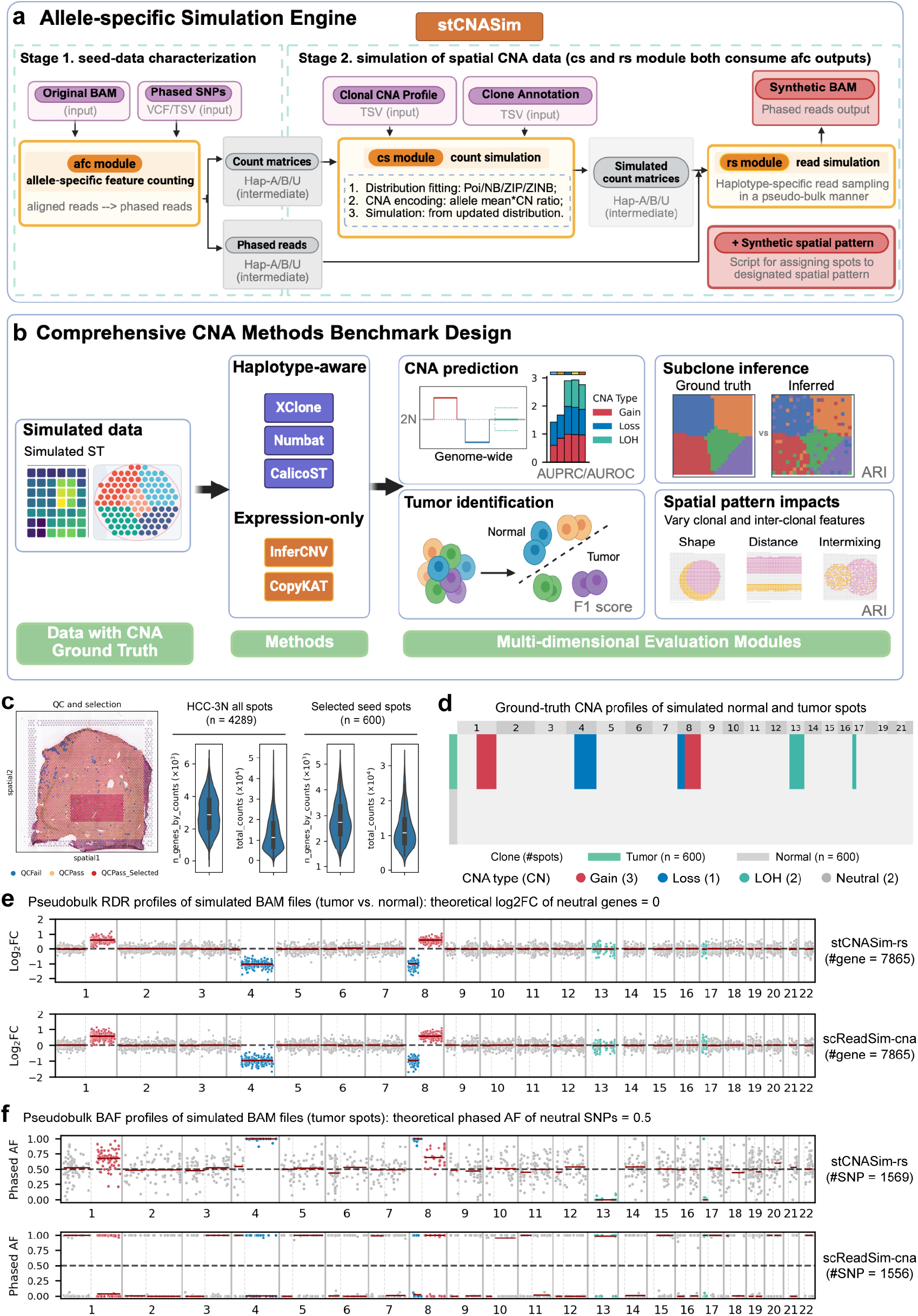
Overview of stCNASim, benchmarking experiment design, and CNA signals in simulated data. (a) Workflow of stCNASim: an allele-aware simulator for generating spatial transcriptomics reads under user-specified CNAs. (b) Schematic of the benchmarking framework used to evaluate CNA detection performance. (c) Seed data generated from the HCC-3N dataset used for simulation. (d) Ground-truth copy number (CN) states assigned to simulated normal and tumor spots. (e) Pseudobulk log2 fold-change (log2FC) profiles comparing tumor and normal spots. (f) Pseudobulk phased allele frequency (AF) profiles of tumor spots. Panels (a) and (b) were created with www.BioRender.com.

As input, stCNAsim requires a seed spatial transcriptomics dataset in BAM format (10x Genomics Visium with polyA selection is the current default), a phased VAF file for relevant SNPs, and a pre-defined CNA profile of clones together with their spatial locations for individual cells or spots. Then, stCNAsim will output a new BAM file and allele-specific count matrices across spatial spots and genes. Briefly, the entire simulation workflow is built upon allele-resolved processing across all core modules, organized into two major functional stages with four sequential core modules and an auxiliary spatial patterning module (Fig. 1a and Supplementary Fig. S1). Stage 1 focuses on seed data characterization to derive three allele-specific gene expression count matrices (allele-A, B, and unclear) by extracting from the seed BAM using the allele-feature-counting (*afc*) module. Specifically, this module will flag the haplotype for each gene-associated UMI by matching the input phased SNPs, and assign each UMI into one of the three disjoint groups: haplotype A (Hap-A), haplotype B (Hap-B), and an unknown (ambiguous) haplotype category (Hap-U). At the junction between Stages 1 and 2, core statistical modelling will be performed: 1) allele-specific probabilistic marginal distributions are fitted to the seed data (using library size and gene allelic mean) with multiple candidate distributions, e.g., negative binomial, and 2) the fitted distributions are adapted to the synthetic CNA states by encoding the CNA profile as allele-specific multiplicative shifts in the distribution mean parameters. With the revised marginal distributions of the three haplotype-specific UMI counts for each gene in each spot, Stage 2 generates new counts by sampling from the adjusted distributions by the count-simulator (*cs*) module, followed by UMI-level read resampling from the original seed BAM with SNP-level haplotype masking with the read-simulator (*rs*) module.

Collectively, these interconnected modules generate paired count-level and read-level spatial transcriptomic data with fully annotated, *in silico* ground-truth CNA states and clonal labels for downstream benchmarking. More details of the simulation framework can be found in the Methods section. Of note, morphological images, including H&E staining, are not applicable in this simulator.

### 2.2 Multi-metric evaluation demonstrates stCNASim as a unique and accurate allele-specific spatial reads simulator

To evaluate stCNASim, we compared its synthetic profiles against those generated by alternative simulators. We initiated this benchmarking process by constructing a seed dataset to serve as a shared input template for all simulation pipelines. This seed dataset was derived from a 10x Visium spatial transcriptomics profile of a hepatocellular carcinoma (HCC) specimen characterized by [23]. Specifically, we subset data from the HCC-3N normal tissue section, which originally contained 4,289 spatial spots. The resulting seed resource retains 600 high-quality normal spots and includes a panel of 8,866 phased single nucleotide polymorphisms (SNPs) compiled across all original spots (Fig. 1c; Supplementary Fig. S2; Methods). The complete seed spot barcode list with corresponding HCC-3N cluster annotations can be found in Supplementary Data 2, and the full set of phased SNPs is provided in Supplementary Data 3. Quality control analyses revealed that the full original tissue spots had a median of 2,905 expressed genes and a median of 11,340 total UMIs per spot. By comparison, our filtered seed spots yielded a median of 2,736 expressed genes and a median of 10,769 total UMIs per spot. This dataset is used as the default seed data unless otherwise specified.

Using these 600 seed spots and 8,866 phased SNPs as a reference template, we utilized stCNASim to generate a synthetic dataset comprising 600 normal and 600 tumor spots (Methods), with the ground-truth CNA profiles visualized in Fig. 1d. This synthetic dataset, designated the stCNASim-validation dataset, was employed to benchmark the count and read simulation performance of stCNASim against alternative simulators. Specifically, within the stCNASim-validation dataset, no CNAs were incorporated in the synthetic normal spots, while six unique CNA patterns were introduced into synthetic tumor spots: (1) two copy gain events on chr1q and chr8q, each assigned a copy number of 3; (2) two copy loss events on chr4q and chr8p, each assigned a copy number of 1; (3) two LOH events on chr13q and chr17p, each assigned a copy number of 2 with uniparental origin. We supplied these two clonal populations and six tumor-specific CNA events as simulation parameters to stCNASim, then executed the tool to produce the synthetic CNA-containing gene count matrix and matching BAM file.

stCNASim is aimed to recapitulate the intrinsic properties of real seed data, preserving its spot-level and gene-level features across copy-neutral genomic regions, while accurately encoding CNA-driven molecular signatures within altered segments: total count signals summarized as read depth ratio (RDR) values [11, 12], and allelic imbalance captured via B-allele frequency (BAF) measurements [11, 12]. To our knowledge, no existing method supports CNA-aware count and read simulations for spatial transcriptomics data. We therefore benchmarked stCNASim against adapted single-cell transcriptomics pipelines, specifically incorporating custom CNA-aware variants of the count simulator scDesign2 [20] and the read simulator scReadSim [22]. First, we evaluated the *afc* module by comparing its allele-agnostic UMI counts (aggregated across the A, B, and U matrices for stCNASim) against those from SpaceRanger v1.1.0 and STARsolo [24] on the input seed dataset. We observed nearly identical results across a diverse range of metrics, including library size, zero-value proportions, and bivariate relationships (such as gene mean versus zero proportion, gene variance, and the coefficient of variation; Supplementary Fig. S3). Similarly high concordance was achieved when evaluating data simulated by the *cs* and *rs* modules (Supplementary Figs. S4 and S5), demonstrating the robust, high-fidelity performance of all three stCNASim modules (*afc, cs*, and *rs*). Second, due to the lack of existing CNA count simulators, we adapted scDesign2 to incorporate CNA signals, creating a baseline variant termed scDesign2-cna (Supplementary Fig. S6). Overall, stCNASim and scDesign2-cna produced highly consistent count simulations for both normal and tumor cells. This consistency extended to the read depth ratio (RDR) across various CNA states (loss, gain, and LOH; Supplementary Fig. S7) and pseudo-bulk chromosomal profiles visualized via log2 fold-change (log2FC) contrasting tumor and normal spots (Supplementary Figs. S8–S9). However, regarding library size simulation, only stCNASim accurately reproduced a distribution comparable to the input seed data, whereas scDesign2-cna yielded a significantly narrower, over-concentrated distribution (Supplementary Fig. S7a). Another minor issue is that scDesign2-CNA’s simple multiplicative approach cannot faithfully model copy-number-induced variability in zero distributions; for example, the copy gain failed to decrease the zero proportion, as this strategy cannot amplify zero to a larger value (Supplementary Fig. S7e). Third, we compared stCNASim’s read-level simulator to scReadSim. Since scReadSim is not natively compatible with CNA analysis, we developed an adapted version, scReadSim-cna, by replacing its aligner (switching Bowtie2 to CellRanger to retain the `xf` tag) and updating its count generator to stCNASim (or scDesign2-cna to handle continuous values and CNA-induced shifts; Supplementary Figs. S10–S11). While both scReadSim-cna and stCNASim showed strong agreement in marginal read-count statistics and pseudo-bulk RDR (Fig. 1e and Supplementary Figs. S12–S13), scReadSim-cna failed to generate allele-specific reads, as it relies on a single reference genome rather than distinguishing paternal and maternal alleles (Fig. 1f and Supplementary Figs. S14–S18). In contrast, stCNASim accurately modeled allelic shifts across all CNA states (gain, loss, and LOH), establishing it as the only available tool capable of simulating allele-specific reads for spatial transcriptomics.

In summary, when evaluating global transcriptomic metrics and CNA-associated RDR signatures, stCNASim performs equivalently to the modified scReadSim-cna baseline, with both tools closely recapitulating seed dataset statistics. StCNASim, however, delivers an indispensable advantage for spatial CNA bench-marking: it generates biologically realistic, continuous haplotype-resolved allelic frequency distributions by explicitly integrating phased SNP information. In contrast, scReadSim-cna’s single-reference-haplotype framework introduces irreversible allelic skew that corrupts all BAF signals, making it inappropriate for validating allele-aware CNA detection workflows. Collectively, these results establish stCNASim as a more robust simulation tool for benchmarking analytical pipelines that jointly interrogate total copy-number changes and allelic imbalance from spatial transcriptomic BAM files. More details of the simulator evaluation can be found in the Supplementary Notes.

### 2.3 Stress-testing spatial CNA prediction using diverse stCNASim-generated datasets

We designed a comprehensive benchmark to assess five widely adopted CNA detection algorithms using 46 synthetic datasets generated with stCNASim (Fig. 1b). This benchmark encompassed two expression-only methods: InferCNV [3] and CopyKAT [9], as well as three allele-aware tools: Numbat [11], XClone [12], and CalicoST [16]. InferCNV and CopyKAT model copy-number alterations exclusively from total gene expression count matrices. The remaining three algorithms additionally integrate allelic frequency data via alternate-allele depth (AD) and total allelic depth (DP) matrices. XClone supports flexible modeling that incorporates allelic counts either independently or alongside total expression signals, whereas Numbat and CalicoST produce unified joint expression–allele results via integrated statistical frameworks. Notably, CalicoST is the only tool of the five algorithms explicitly designed for spatial transcriptomic data. It incorporates spatial autocorrelation into its CNA inference framework, while the remaining four methods were initially built for single-cell RNA-seq analysis and lack native support for leveraging spatial coordinate information. Here, we evaluated the performance of five CNA analysis tools in predicting copy-number states for a single tumor clone. To stress-test these tools, we varied key technical parameters to their extreme values, including the presence and proportion of reference cells, tumor purity, sequencing depth, and the occurrence of whole-genome duplication (WGD). The inferred genome-wide CNA profiles were then compared against ground-truth states—encompassing copy-number gains, losses, loss of heterozygosity (LOH), and WGD events—using the Area Under the Receiver Operating Characteristic (AUROC) and Area Under the Precision-Recall (AUPRC) metrics.

First, we evaluate the effects of the presence/absence of reference spots at different tumor purity levels. Reference spot provision is optional for InferCNV, CopyKAT, and CalicoST, yet mandatory for Numbat and XClone to complete CNA inference. To characterize how reference spot count shapes CNA detection accuracy, we generated nine synthetic datasets spanning tumor purities from 1% to 99% (Supplementary Fig. S19b). Across all datasets, the number of reference spots was fixed at 600, while the combined pool of normal and tumor spots remained constant at 1000; tumor spot counts ranged from 10 to 990 to create a continuous tumor purity gradient. These datasets were used to benchmark tumor purity effects on CNA detection performance throughout subsequent analyses.

Comparing performance with and without reference cells revealed that InferCNV, CopyKAT, and CalicoST consistently achieved higher CNA detection accuracy when a reference was utilized, particularly in samples with high tumor purity. In the absence of a reference, InferCNV and CopyKAT at 90% tumor purity incorrectly predicted inverted copy-number gain and loss states between tumor and normal spots, while CalicoST failed to detect any expected CNAs at 99% tumor purity (Fig. 2a–c; Supplementary Figs. S20–S24). When matched spatial normal spots are unavailable, external single-cell transcriptomic profiles of matching cell types offer a viable alternative reference. For instance, incorporating an external reference of 655 single-cell normal hepatocytes from a separate hepatocellular carcinoma (HCC) study (Lu et al., [25]) corrected InferCNV’s predictions at 90% tumor purity, regardless of whether the analysis included 1,000 spatial spots or was downsampled to 200 (Supplementary Figs. S20c,d). Furthermore, CalicoST demonstrated robustness to spatial separation, maintaining performance at 97% and 99% tumor purity even when a 20-spot spatial gap was introduced between the reference and target tumor regions (Supplementary Fig. S25).

**Fig. 2.**
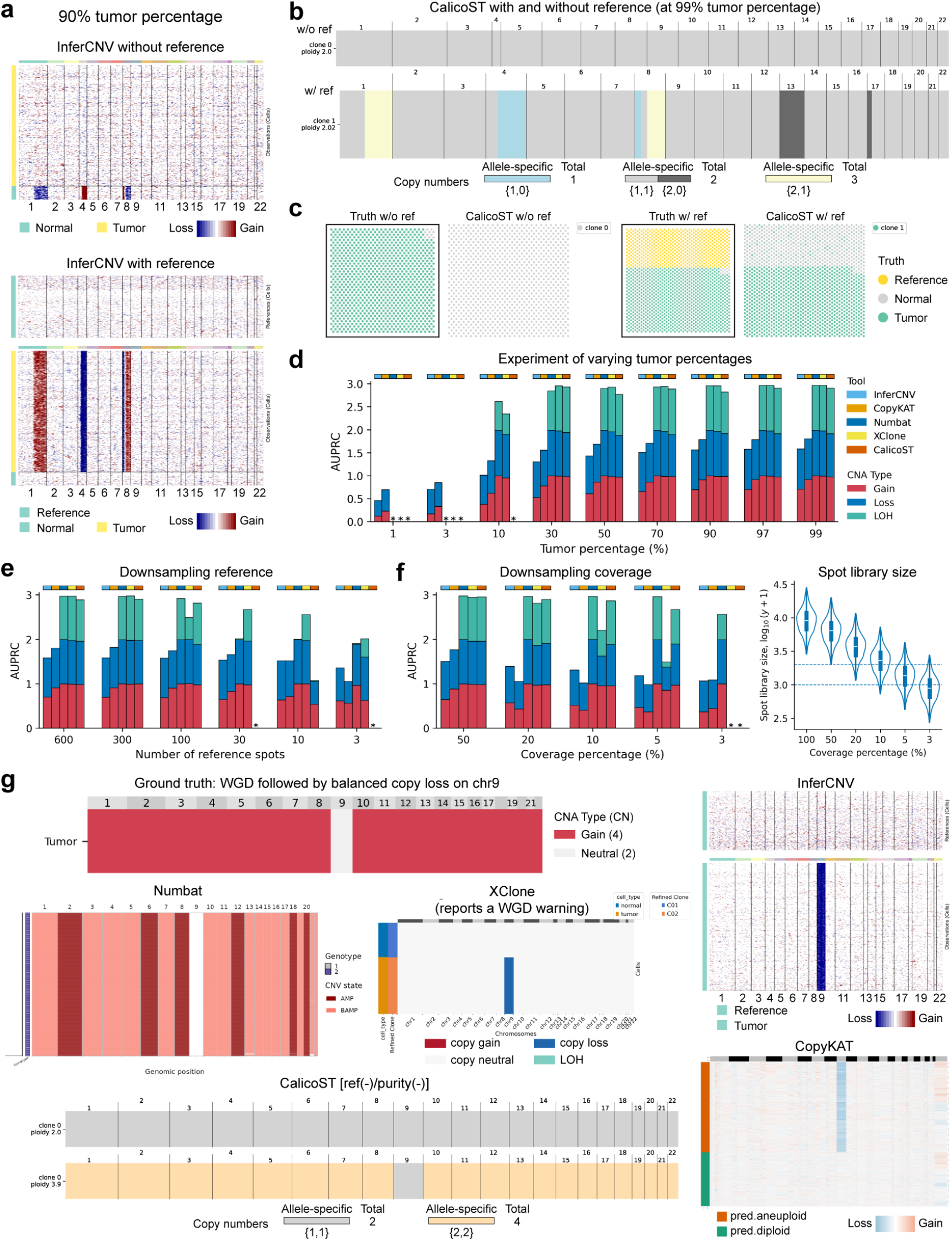
Benchmarking of CNA profile prediction with simulated data. (a-d) Evaluation across varying tumor percentages. (a) InferCNV-predicted CNA profiles with and without reference at 90% tumor percentage. (b) CalicoST-predicted CNA profiles with and without reference (i.e., under the [ref(+)/purity(+)] and [ref(-)/purity(-)] settings) at 99% tumor percentage. (c) Spatial distribution of ground truth and CalicoST-identified tumor spots (with/without reference) at 99% tumor percentage. (d) AUPRC of predicted CNAs (all tools using reference) versus tumor percentage. (e-f) AUPRC metrics under downsampling conditions: (e) reference spots and (f) sequencing coverage. (f; right panel) The two blue dashed lines mark spot library sizes (total UMI counts) of 1000 and 2000, respectively. The x-axis coverage is at the read level. (g) Prediction of whole-genome duplication (WGD) followed by balanced copy loss on chr9. Note: XClone reported a WGD warning; CalicoST results are shown only for the [ref(-)/purity(-)] setting due to failure of the [ref(+)/purity(+)] run. In AUPRC plots, an asterisk (*) indicates no CNA was detected or that a technical error occurred

Conversely, at the opposite extreme of low tumor purity (10%), CNA detection becomes highly challenging even when using a reference. At this threshold, only Numbat and XClone retained moderate predictive power (Fig. 2d; Supplementary Fig. S26 and S27). For example, Numbat’s 3-stack bar chart of AUPRC decreased from 2.840 (out of 3) at 30% purity to 2.612 at 10% purity, whereas CalicoST failed due to technical errors in [ref(+)/purity(+)] setting and missed CNA in [ref(-)/purity(-)] setting (Supplementary Fig. S22 and S23). At ultra-low purities of 1% and 3%, none of the tested methods yielded reliable results, though CopyKAT and InferCNV completed execution. The complete AUPRC and AUROC metrics at all purity levels for all tools run with reference spots are summarized in Fig. 2d and Supplementary Fig. S19c. At intermediate-to-high tumor purities (≥ 30%), the three allele-aware methods outperformed the expression-only tools overall. Among these methods, XClone achieved the highest average combined AUPRC (2.950), marginally surpassing Numbat (2.925). At low tumor fractions (≤ 10%), expression-only tools retained the practical advantage of consistently generating complete outputs, albeit with substantially reduced predictive power. The three allele-aware pipelines either encountered technical errors or exhibited degraded performance at low purity thresholds, likely driven by insufficient detectable allelic imbalance and copy-number shifts within sparse tumor populations.

Second, we evaluated tool sensitivity to reference spot abundance by sequentially reducing the number of available reference spots from 600 down to 3, while holding the total tumor spot count fixed at 1000 (Fig. 2e; Supplementary Fig. S28 - S29). Detection of copy gains and losses remained robust across all reference spot abundances for every tested algorithm. However, LOH calling accuracy declined sharply as reference spot numbers diminished. Numbat exhibited pronounced sensitivity to limited reference populations for clonal LOH detection, as illustrated by side-by-side comparison of Supplementary Fig. S28b and c. Among the three allele-aware tools, XClone exhibited markedly higher robustness to limited reference spot numbers relative to Numbat and CalicoST.

Third, we characterized how reduced sequencing depth alters CNA prediction performance by incrementally downsampling sequencing coverage to 5% of the original read depth (Fig. 2f; Supplementary Fig. S30). Numbat maintained the highest prediction robustness even at minimal coverage, with its pseudobulk aggregation strategy effectively mitigating low-depth noise. CalicoST retained interpretable CNA profiles at 5% coverage, yet clone assignment results exhibited weak, noisy signals across many tumor spots. XClone experienced the most severe degradation of CNA resolution as sequencing depth was depleted.

Last, we examined a special case: whole-genome duplication with balanced chr9 copy loss. Detecting whole-genome duplication (WGD) from transcriptomic sequencing data presents a well-established analytical challenge, as uniform ploidy doubling is difficult to disentangle from global increases in sequencing coverage. To benchmark algorithm performance on this complex scenario, we generated a specialized synthetic dataset featuring global WGD followed by focal balanced copy loss on chromosome 9 (Fig. 2g; Supplementary Fig. S31). This design produces a distinctive allelic signature indicative of an underlying WGD event: under a tetraploid baseline, the balanced chr9 copy loss generates an allelic frequency (AF) of 0.5, which differs starkly from the AF = 0 or 1 profiles typical of canonical unbalanced copy loss modeled with a diploid baseline. The two expression-only methods, InferCNV and CopyKAT, only detected the chr9 focal copy loss and failed to identify the genome-wide WGD event. Numbat successfully recovered both the WGD signature (manifest as universal chromosomal amplifications) and the focal balanced copy loss on chr9 (which appears copy-neutral under a diploid baseline). However, the tool erroneously classified balanced copy gains across six chromosomes, including chr2 and chr6, as unbalanced allelic alterations. XClone correctly identified the chr9 copy loss and output an explicit WGD warning within its runtime log files, despite not formally integrating the ploidy shift into its final copy-number segmentation. CalicoST only generated complete valid outputs under the reference-free [ref(-)/purity(-)] workflow; the reference-enabled [ref(+)/purity(+)] pipeline terminated with technical errors. When run with its 4-ploidy baseline setting, the [ref(-)/purity(-)] configuration accurately resolved both the global WGD signature and focal chr9 loss; no CNAs were detected when utilizing CalicoST’s default 2-ploidy baseline.

To summarize, single-cell-designed CNA inference algorithms are adaptable to spatial transcriptomics data. Compared to reference-free mode, supplying reference spots, either matched spatial normal tissue or external non-spatial single-cell normal profiles, consistently boosts CNA detection accuracy. For CalicoST, when the optimal reference-enabled [ref(+)/purity(+)] pipeline fails due to technical errors, the reference-free [ref(-)/purity(-)] workflow provides a viable fallback. All tested algorithms suffer degraded performance at low tumor purity thresholds. Numbat delivers superior resilience to depleted sequencing coverage, while XClone is the most robust algorithm when working with limited numbers of reference spots. LOH detection accuracy drops markedly for all tools when reference spot populations are sparse, whereas copy gain and loss calling remain largely unaffected by reference depletion. Only Numbat, XClone, and CalicoST (when configured with a 4-ploidy baseline) reliably resolve complex combinatorial events involving WGD superimposed on focal balanced copy loss.

### 2.4 Assessing tumor identification by simulating diverse tumor percentages and CNA complexities

One common utility of CNA detection is to identify tumor cells from the whole cell population in the spatial tumor microenvironment. Therefore, we generated synthetic datasets to assess the accuracy of the predicted CNA states in distinguishing tumor from normal cells. These datasets correspond to the aforementioned datasets spanning a range of tumor purities utilized for CNA prediction evaluation (Supplementary Fig. S19b). Each dataset contains one tumor clone alongside normal cells, with a total of 1,000 spots, and features the ground-truth CNA profile illustrated in Fig. 1d. To ensure a fair comparison, a reference set of 600 spots was utilized for all tools. Under a balanced mixing ratio (equal numbers of normal and tumor cells), all five tools accurately separated tumor from normal cells (Fig. 3a; Supplementary Figs. S32–S33).

**Fig. 3.**
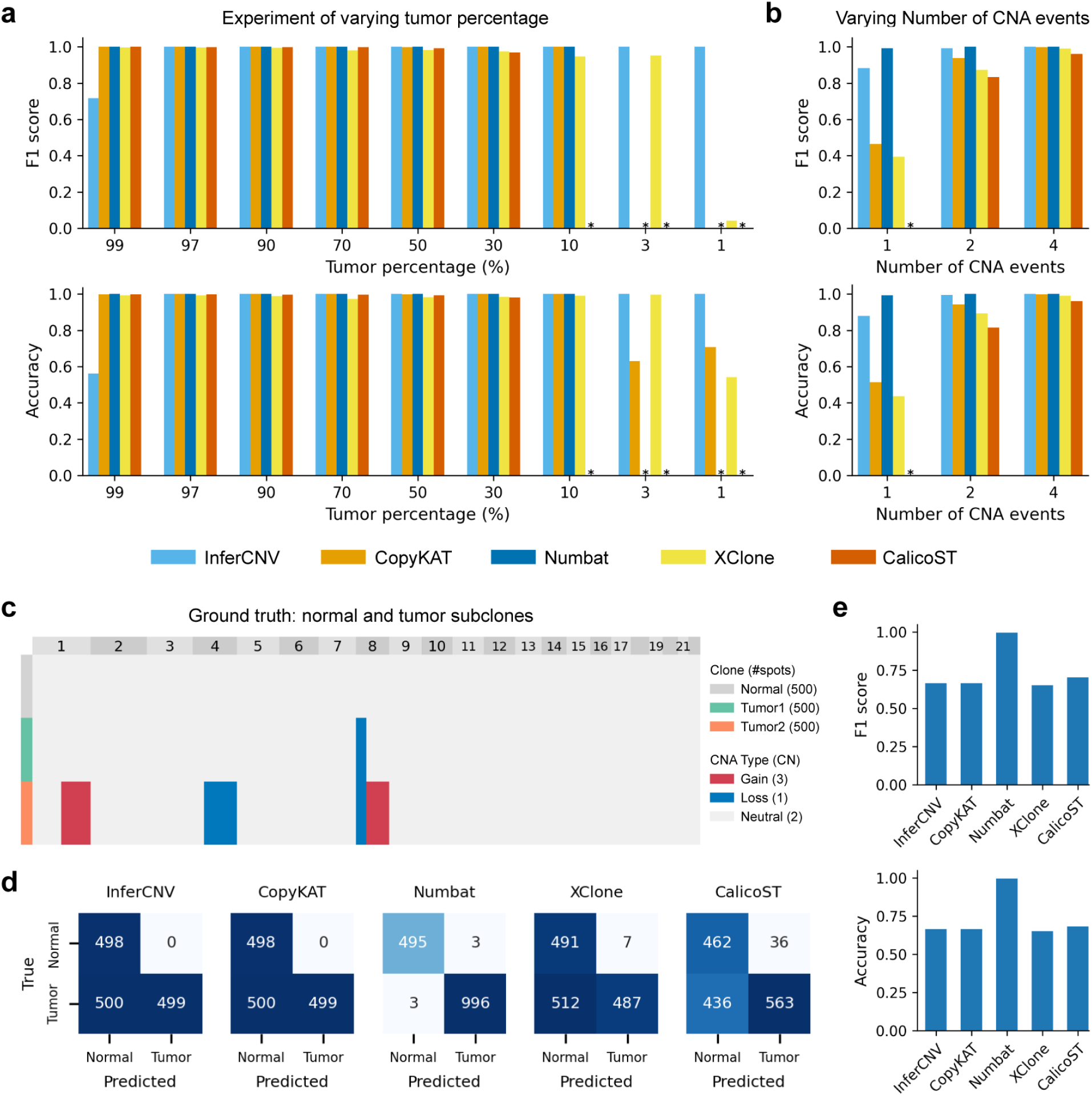
Benchmarking of tumor identification with simulated data. (a) Evaluation across varying tumor percentages. Each simulation contains 1 normal and 1 tumor clones. (b) Evaluation across varying number of CNA events in the tumor clone. Each simulation contains 1 normal and 1 tumor clones. (c-e) Experiment combining normal and tumor subclones with distinct CNA profiles. (c) Ground-truth CNA profiles of the simulated normal and tumor subclones. #spots: number of spots; CN: copy number. (d) Confusion matrices comparing ground-truth and predicted normal/tumor labels. (e) Performance metrics. ‘Tumor’ was treated as positive label in metric calculation. When plotting the F1 score and accuracy metrics, an asterisk (*) indicates no CNA was detected or a technical error occurred.

However, when the tumor proportion was reduced to extremely low levels, only InferCNV remained robust and accurate down to a 1% proportion (achieving both F1 score and accuracy >0.99). In contrast, the other methods failed to yield reasonable results at various detection thresholds: CalicoST failed at 10%, Numbat and CopyKAT at 3%, and XClone at 1% (Fig. 3a; Supplementary Fig. S33). Surprisingly, in scenarios where tumor cells were extremely dominant (99%), InferCNV exhibited degraded performance, misclassifying 437 out of 990 tumor spots into the normal group despite the use of reference cells (Fig. 3a; Supplementary Fig. S33). This is likely due to the inherent challenges of clustering under extreme class imbalance.

We next evaluated the impact of CNA complexity by gradually reducing the number of CNA events from four to two and one (Fig. 3b and Supplementary Fig. S34). Across all scenarios, Numbat demonstrated highly robust and accurate tumor cell detection, even with only a single CNA event (chr8p loss). InferCNV followed closely, showing decreased performance only in the single-CNA scenario, where it still maintained an F1 score of 0.882. Conversely, CopyKAT, XClone, and CalicoST exhibited substantial performance declines as the number of CNA events decreased, yielding F1 scores of 0.938, 0.872, and 0.833, respectively, for two CNA events, and 0.465, 0.396, and a technical failure, for a single CNA event.

Furthermore, we merged the tumor cells with four CNAs and those with one CNA to simulate a two-clone tumor and investigated whether the tools could still successfully identify all tumor cells (Figs. 3c–e; Supplementary Fig. S35). Notably, the clone harboring only a single CNA event (chr8p loss) shares a high similarity with normal cells in terms of absolute copy number differences. This high similarity explains why, when mixed solely with normal cells, this clone went undetected by all methods except Numbat. Within this two-clone mixture, Numbat remained highly robust, identifying nearly all tumor cells across both distinct subclones (F1 score = 0.997). Unsurprisingly, the other benchmarked methods failed to effectively detect the single-CNA clone.

### 2.5 Evaluation of subclone inference using simulated allele-specific CNAs

Next, we investigated whether the predicted CNA states could resolve the sub-clonal structures within tumor cell populations. To evaluate this, we again utilized stCNASim to generate simulated datasets under challenging scenarios, specifically focusing on loss of heterozygosity (LOH) and allele-specific copy number alterations. In the first scenario, we simulated two subclones that shared four clonal (trunk) CNA events (gains on chr7p and 11q, and losses on chr9p and 16q), while clone 2 harbored two unique subclonal LOH events on chr1p and 6q (Fig. 4a). As expected, the two allele-agnostic methods, InferCNV and CopyKAT, failed to separate these subclones because they completely ignore allelic information (Fig. 4b; Supplementary Fig. S36). In contrast, all allele-aware methods successfully distinguished these two equally proportioned clones. Notably, CalicoST achieved the highest clustering accuracy, yielding a superior Adjusted Rand Index (ARI) and misclassifying only 9 out of 1,000 spots, compared to 51 and 63 misclassified spots for Numbat and XClone, respectively.

**Fig. 4.**
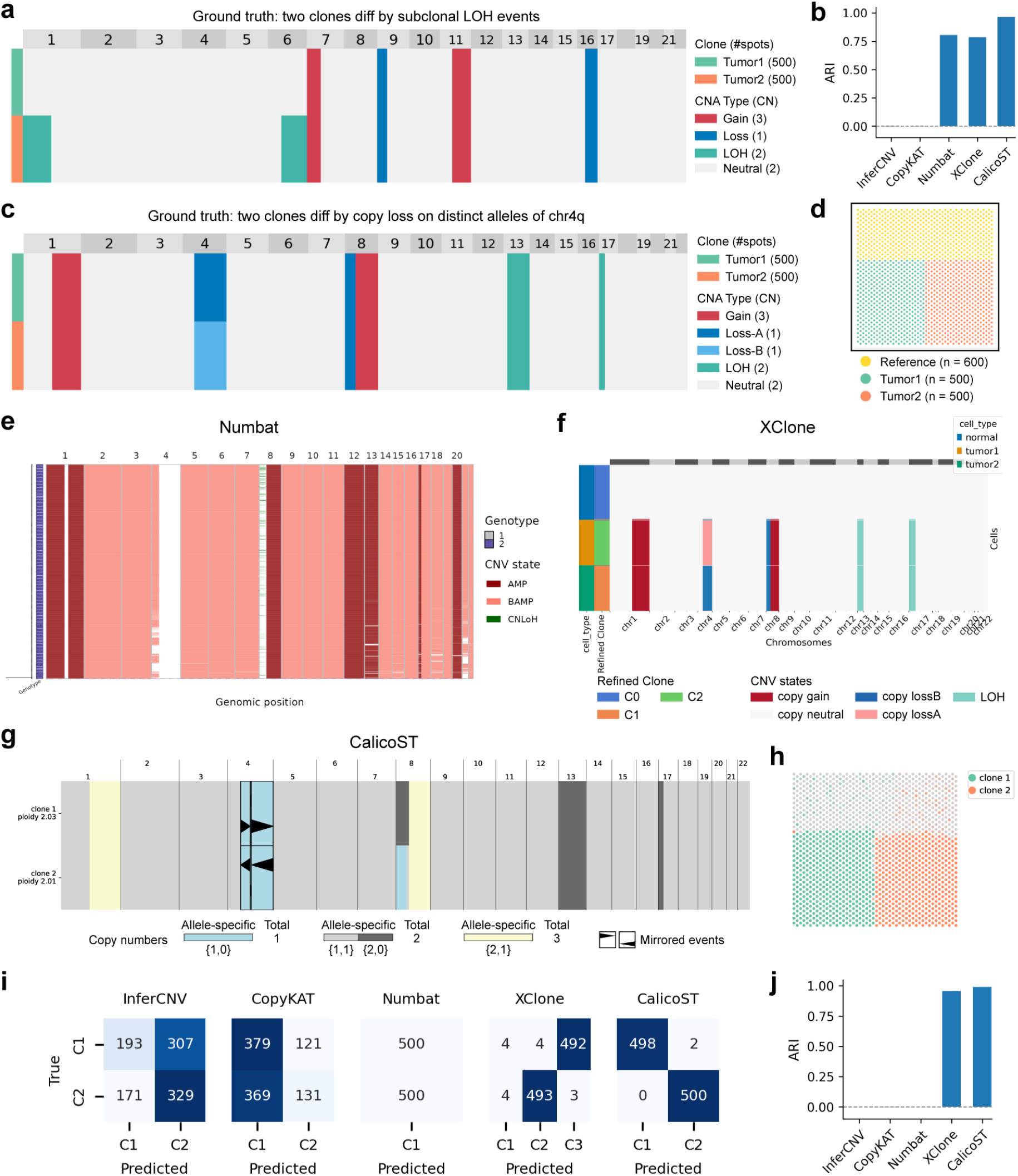
Benchmarking of subclone inference with simulated data. (a-b) Benchmarking using simulated data where two tumor clones diff by subclonal LOH events on chr1p and chr6q. (a) Ground-truth CNA profiles of two simulated tumor clones. (b) ARI metrics for ground-truth versus predicted clone labels. (c-j) Benchmarking using simulated data where two tumor clones diff by copy loss on distinct alleles of chr4q. (c) Ground-truth CNA profiles of two simulated tumor clones. (d) Spatial distribution of ground-truth clones. (e-g) Predicted CNA profiles by: Numbat (e), XClone (f), and CalicoST (g). (h) Spatial distribution of clones identified by CalicoST. (i) Confusion matrices comparing ground-truth and predicted clone labels. (j) ARI metrics for ground-truth versus predicted clone labels.

In the second scenario, clones 1 and 2 shared five clonal CNA events but featured distinct allele-specific losses on chr4q (Fig. 4c–j). Consistent with the first scenario, InferCNV and CopyKAT were unable to distinguish these two subclones (Fig. 4i; Supplementary Fig. S37). Surprisingly, Numbat also failed to differentiate between the two distinct lost alleles in these clones. Conversely, CalicoST and XClone successfully resolved the two subclones with high accuracy (Fig. 4e–j). A third simulation involving allele-specific LOH on chr13q further confirmed Numbat’s limitation in detecting subclones characterized by allelic divergence on the same chromosome. In contrast, CalicoST and XClone retained strong partitioning capabilities; out of 1,000 spots, CalicoST made only 2 errors, while XClone made 43 errors (Supplementary Fig. S38).

### 2.6 Benchmarking the effect of cell spatial patterns on CNA calling

Notably, although InferCNV, CopyKAT, Numbat, and XClone can be applied to ST data, they do not incorporate spatial information for CNA calling. Conversely, CalicoST is specifically tailored for ST data and leverages a hidden Markov random field (HMRF) to model spatial correlation [16]. However, the precise impact of integrating the spatial information on CNA calling remains uncharacterized. To address this, we generated simulated datasets featuring diverse spatial architectures using stCNASim. Specifically, we generated a 2-clone tumor and normal mixture with a total of 2000 spots, among which the two tumor clones share copy gain on chr1q and 8q and loss on chr8p, while clone 2 has unique copy loss on chr4q and LOH on 13q and 17p (Fig. 5a). Since InferCNV, CopyKAT, Numbat, and XClone do not use spatial coordinates, we found they perform reasonably well in both CNA prediction and subclone detection (slightly lower performance for CopyKAT in CNA state prediction and XClone in subclone identification; Fig. 5b-c and Supplementary Fig. S39).

**Fig. 5.**
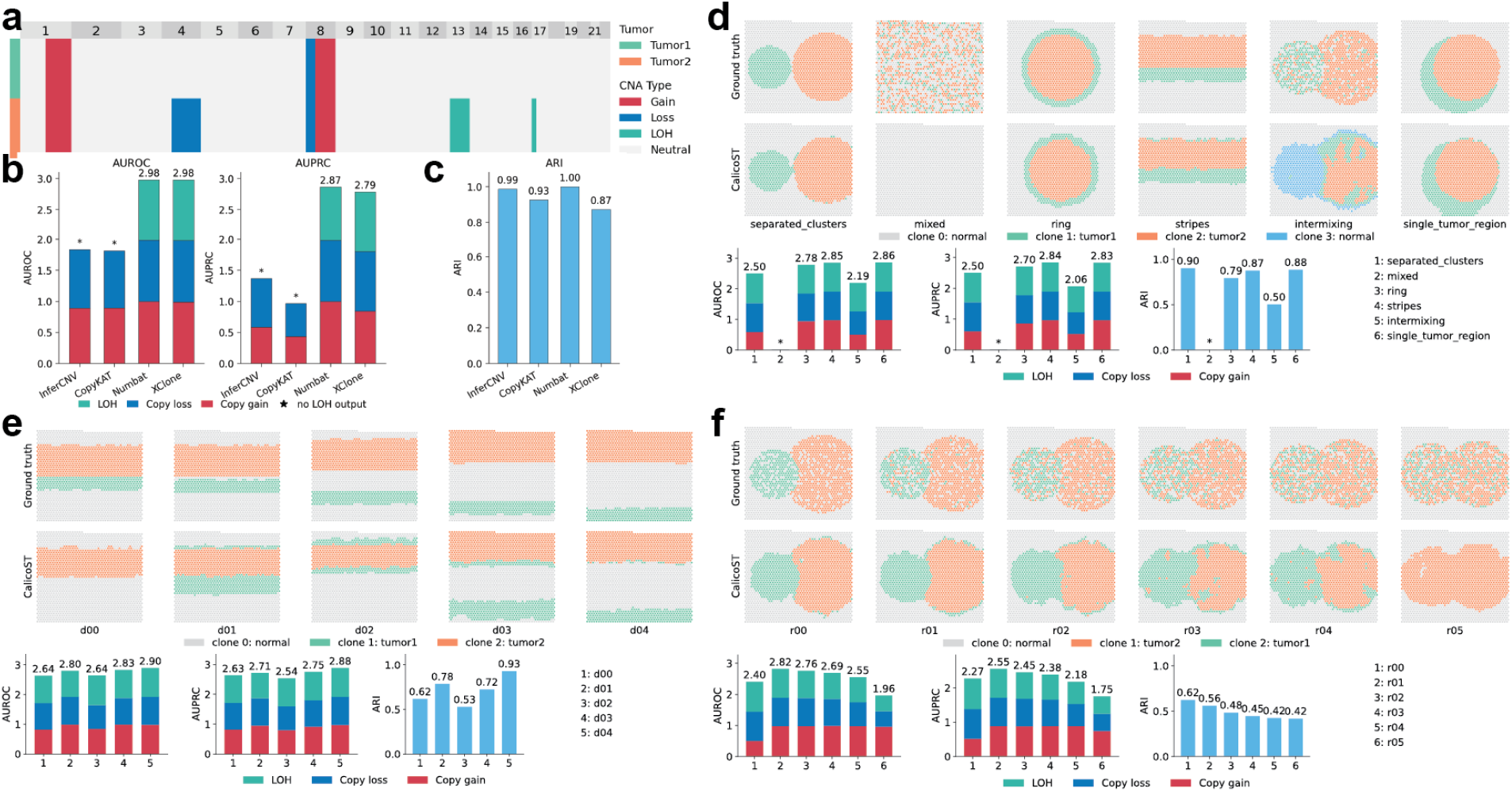
Benchmarking the effect of varying cellular spatial patterns on CalicoST using simulated data. (a) Ground-truth CNA profiles of two simulated tumor clones. (b) Inference accuracy (AUROC and AUPRC) of CNAs inferred by InferCNV, CopyKAT, Numbat, and XClone. (c) ARI evaluating clonal assignment accuracy for InferCNV, CopyKAT, Numbat, and XClone relative to ground truth. (d-f) Ground-truth versus CalicoST-predicted cellular spatial patterns, along with corresponding CNA inference performance (AUROC/AUPRC) and clonal assignment accuracy (ARI), evaluated under varying: (d) tumor clone shapes, (e) inter-clonal distances, and (f) cell intermixing rates across tumor clones.

By fixing the clone CNA profiles and systematically varying only the input cell coordinates in six different spatial distributions (separated cluster, mixed, ring, stripes, intermixing, and single-tumor region; Fig. 5d and Supplementary Fig. S40), we evaluated how distinct spatial cell arrangements influence CalicoST in terms of CNA calling and subclone inference. Because the HMRF model operates on the assumption that adjacent spatial spots exhibit greater transcriptional similarity, we hypothesized that CalicoST would demonstrate superior performance under structurally simplified spatial cell patterns, such as homogeneous tumor clusters with well-defined boundaries. Our evaluation largely validated this hypothesis. In the most extreme scenario, where the spatial distribution of normal and tumor cells was completely randomized and lacked discernible structure, CalicoST failed to detect any CNA events or identify tumor subclones (scenario 2 for full mixed; Fig. 5d). Conversely, when tumor cells formed coherent clusters with distinct geometries devoid of cell intermixing, both the CNA calling and subclone inference of CalicoST were generally more accurate, despite degraded performance in clustering tumor clones in intermixing and ring settings (respective ARI: 0.5 and 0.79; Fig. 5d).

Furthermore, as the distance between the two distinct tumor clones increased in the stripe scenarios (Fig. 5e and Supplementary Fig. S41), we observed a moderate increasing trend in the performance of CalicoST. Finally, incremental increases in the interclonal cell mixing rate led to a progressive degradation in both CNA calling accuracy and tumor subclone inference resolution (Fig. 5f and Supplementary Fig. S42).

## 3 Conclusion, guidelines and discussion

By using real sequencing data as the seed, our simulator stCNASim retains realistic technical and biological properties, including gene expression distributions, allele-specific read coverage, and SNP-level phasing information. Collectively, stCNASim delivers paired count-level and read-level spatial transcriptomic datasets with fully annotated *in silico* ground-truth CNA states and clonal labels, all generated through end-to-end allele-specific processing. This unique design makes it a robust, dedicated platform for comprehensive evaluation of spatial CNA detection and analysis tools. To the best of our knowledge, there is currently no alternative simulation tool designed to generate allele-specific copy number alterations (CNAs) in single-cell or spatial transcriptomics (ST) data. While existing packages offer partial capabilities—such as scDesign3, which supports covariates in generating count matrices, and scReadSim, which generates reads from a reference genome—they lack the ability to model allele- or haplotype-specific events. This gap positions stCNASim as a pivotal resource to ensure that proposed computational models in this domain are reliable and their underlying theories are testable.

By systematically varying simulation parameters to encompass extreme biological and technical conditions, we performed a rigorous stress test on five representative CNA inference tools. Their performances are summarized in Supplementary Data 11. Briefly, our benchmarking revealed several key insights:

- **Adaptability of single-cell methods:** The four single-cell-based methods adapt remarkably well to ST data overall, despite completely ignoring spatial coordinates.
- **Spatial integration:** The ST-specific method, CalicoST, successfully leverages spatial autocorrelation, though its performance degrades under high spatial intermixing of tumor and normal cells.
- **Context-specific strengths:** Each tool exhibits distinct advantages depending on the scenario. Infer-CNV is highly effective at tumor identification when the tumor cell proportion is extremely low; CopyKAT offers superior computational efficiency; XClone is the most robust when reference cell counts are limited; Numbat excels at detecting tumor cells and subclones; and CalicoST is a versatile tool for resolving diverse allele-specific events.
- **Vulnerabilities under stress testing:** Each tool also revealed specific limitations; InferCNV and Copy-KAT failed to capture allele-specific events, while CalicoST, Numbat, and XClone exhibited sensitivity to low tumor purity, mirrored subclones, and low sequencing coverage, respectively.

Ultimately, while no single tool dominates across all evaluation metrics, certain methods perform exceptionally well in specific challenging scenarios. This benchmarking framework and our corresponding guidelines aim to assist researchers in selecting the most appropriate tool based on their specific biological complexity and technical configurations.

Despite these insights, several limitations of our simulation and benchmarking framework warrant discussion. First, our evaluation focused strictly on single-slide simulations. Multi-slice experimental designs, in both 2D and 3D, are increasingly utilized to map the tumor microenvironment (TME) and clonal architecture across larger tissue volumes. Second, although we incorporated multi-clonal settings, we did not model temporal trajectories or evolutionary branching. Future iterations incorporating multiple clones with explicit evolutionary structures will further enhance therapeutic and clinical relevance. Third, our simulations were restricted to a 55-*µ*m spot resolution (matching standard Visium data). We did not evaluate other spatial platforms with different resolutions, such as Slide-seq V2 (10 *µ*m) or Patho-DBiT (50 *µ*m). Fourth, we did not account for intra-spot cell-type heterogeneity. Future studies could integrate paired single-cell and spatial RNA-seq data as seeds to simulate spatial spots containing diverse cell-type mixtures at varying proportions. Fifth, we omitted explicit modelling of intra-spot gene-gene correlations, inter-spot spatial correlations and cell-cell interactions. Future module optimizations integrating these covariance structures will facilitate studies of molecular and cellular regulation in spatial biology. Finally, because our simulator relies on existing sequencing reads as seeds, it does not support the de novo generation of haplotypes. However, this is unlikely to be a major limitation, given that current cancer genomics studies focus heavily on intra-patient comparisons.

Looking ahead, we expect stCNASim to standardize the evaluation of future CNA detection tools, particularly in resolving the spatial and temporal dynamics of tumor evolution. Moreover, given its capacity to generate highly diverse synthetic datasets at scale, stCNASim could serve as a valuable training engine for supervised deep learning frameworks—akin to the role of synthetic data in training tools like DeepVariant for SNV genotyping—paving the way for the development of robust, off-the-shelf, deep-learning-based CNA prediction models.

## 4 Methods

### 4.1 stCNASim: an allele-aware spatial RNA-seq CNA simulator

To benchmark and validate pipelines for allele-specific copy number alteration (CNA) analysis in spatial transcriptomics, we developed stCNASim, a modular simulator that generates synthetic spatial transcriptomics datasets with fully known ground-truth CNA profiles. The simulator accepts three core inputs: an aligned seed BAM file, a set of phased heterozygous single-nucleotide polymorphisms (SNPs), and a user-defined clonal CNA profile. It outputs simulated alignment files with embedded CNA signals and predefined clonal spatial architectures.

The full simulation pipeline implements allele-resolved logic across all core components and is partitioned into two primary functional stages, comprising four sequential core modules plus an auxiliary spatial patterning module. Throughout the following text, we use the terms haplotype and allele interchangeably.

#### 4.1.1 The preprocessing (*pp*) module

The first stage, seed-data characterization, establishes allele-specific analytical foundations via two key modules: the pre-processing (*pp*) module and the allele-specific feature counting (*afc*) module. This stage mainly takes two inputs: a BAM alignment file and a list of phased SNPs, which encode the two parental haplotypes using binary labels 0 and 1. Such haplotype-resolved SNP data can be derived from multiple bulk, single-cell and spatial omics technologies, such as scDNA-seq [26], single-cell transcriptomics [11, 12] and spatial transcriptomics [16].

The *pp* module validates, filters, and standardizes all input annotation files, ensuring data integrity, format compliance, and genomic coordinate consistency prior to downstream counting and simulation routines. A central preprocessing step involves curating gene feature annotations via two operations: (1) deduplicating gene records with identical gene symbols by retaining only the first encountered entry; and (2) retaining only genes residing on user-specified chromosomes (default: human autosomes 1-22). Notably, stCNASim only extracts chromosomal coordinates (chromosome, start, end positions), gene names, and strand orientation from gene annotations; gene structural features, including exon boundaries, are not incorporated as inputs. The simulator does not natively resolve overlapping gene loci by default; handling of reads mapping to multiple overlapping genes is fully delegated to upstream alignment pipelines such as SpaceRanger (through the `GN` and `xf` tags).

#### 4.1.2 The allele-specific feature counting (*afc*) module

The *afc* module resolve the allelic origins of aligned sequencing reads and UMIs of the input seed BAM file by aggregating the haplotype information from all their harboring phased SNPs. For every gene, the *afc* module assigns each UMI overlapping the gene into one of the three disjoint groups: haplotype A (Hap-A, corresponding to label 0 in the input phased SNPs), haplotype B (Hap-B, corresponding to label 1), and an unknown (ambiguous) haplotype category (Hap-U). This module ultimately outputs allele-stratified (i.e., Hap-A, B, U) sets of gene-wise phased UMIs and associated spot-by-gene count matrices, both of which serve as empirical templates for downstream simulation.

##### Step 1: Data preparation and quality control

1. **Parallel batching of genes**. Genes are loaded and split into batches for parallel processing. Within each batch, every gene, including those with overlapping genomic intervals, is processed separately. This divide-and-conquer strategy enables efficient counting on large datasets with tens of thousands of genes.
2. **Read filtering**. For each gene, all reads intersecting its genomic span are extracted from the input seed BAM file and discarded if they satisfy any of the following exclusion criteria:
  - mapping quality (MAPQ) < 20;
  - aligned length < 30nt;
  - aligned fraction (mapped bases / total read length) < 0.5;
  - SAM flag marking any bit of UNMAP, SECONDARY, QCFAIL;
  - unpaired fragments in paired-end sequencing;
  - UMI barcodes containing ambiguous ‘N’ bases;
  - `GN` tag not match the query gene name;
  - strand orientation incompatible with the annotated gene strand;
  - 10x Genomics-specific `xf` tag not equal to 17 or 25. These two values mark the read is confidently mapped to the transcriptome.
3. **SNP filtering**. Input phased SNPs are pre-filtered to retain only informative ones for subsequent haplotype assignment. SNPs are excluded if either criterion holds: (1) its total aggregated UMI count across all spots < `min_count` (default 1), or (2) its minor allele frequency < `min_maf` (default 0). The default parameters apply lenient filtering, presupposing that input phased SNPs are well-supported and informative.

##### Step 2: SNP-based haplotype assignment via hierarchical inference

To resolve the haplo-type state of each UMI, allelic assignment proceeds through three nested hierarchical aggregation layers: individual phased SNPs - single reads - full UMIs.

1. **SNP level**: For any read spanning a given SNP coordinate, we first assign a SNP-level haplotype state by comparing the observed base at the SNP position against the reference and alternate alleles linked to each parental haplotype. Four mutually exclusive SNP-level states are defined:
  - **A**: Observed base matches the reference haplotype (haplotype index 0).
  - **B**: Observed base matches the alternative haplotype (haplotype index 1).
  - **O (Other)**: Observed base matches neither the reference nor alternative allele and thus lacks support for either parental haplotype.
  - **U (Unknown)**: No base is fetched, e.g., the SNP lies within a split read.
2. **Single read (sread) level**: A single read can cover multiple SNPs, each contributing an independent haplotype vote. The read’s overall haplotype state is determined by the consensus of all SNP votes. The five possible states are:
  - **A**: No SNPs are in state B; and at least one SNP is in state A.
  - **B**: No SNPs are in state A; and at least one SNP is in state B.
  - **D (Dual)**: The sread carries both A-state and B-state SNPs.
  - **O (Other)**: No SNPs are in state A or B; and at least one SNP is in state O.
  - **U (Unknown)**: Either (1) the read covers no phased SNPs; or (2) No SNPs are in state A, B or O; and at least one SNP is in state U.
3. **UMI level**: All sreads sharing the same cell barcode and UMI sequence form one UMI-level read group. The final UMI haplotype state is assigned via consensus across all its constituent sreads. The five UMI-level states are:

- **A**: No sreads are in state B or D; and at least one sread is in state A.
- **B**: No sreads are in state A or D; and at least one sread is in state B.
- **D**: Either (1) at least one sread is in state D, or (2) the UMI group contains both A-state and B-state sreads.
- **O**: No sreads are in state A, B, or D; and at least one sread is in state O.
- **U**: All sreads are in state U.

While this three-layer hierarchy could be conceptually simplified by directly aggregating SNP-level calls to the UMI level, we retain the full layered structure to preserve modularity and extensibility. Specifically, the sread level constitutes a technically meaningful analytical unit that provides an informative intermediate tier for haplotype assignment. Furthermore, this framework delivers extensibility by supporting stage-specific threshold tuning, such as setting minimum consensus voting rates, at each step of the inference hierarchy.

##### Step 3: Allele-specific UMI counting

Prior to quantification, multi-mapping UMIs (those assigned to multiple distinct genes) are identified and discarded to avoid ambiguous count contributions. We then quantify unique UMIs partitioned across the five haplotype states (A, B, D, O, U) for every spatial spot and annotated gene, yielding five allele-specific spot-by-gene count matrices. Notably, only the count matrices corresponding to states A, B, and U are retained for downstream count simulation; matrices for ambiguous D and O states are discarded entirely.

##### Step 4: Generation of per-gene allele-specific seed BAM files

In addition to generating allele-stratified count matrices, the *afc* module exports reads originating from gene-specific allele-resolved UMIs into individual partitioned BAM files, each aggregating pseudobulk reads across all seed spots. In line with the preceding counting step, only reads annotated to alleles A, B, and U are preserved. These partitioned BAMs serve as the allele-specific read pool for per-gene resampling operations implemented in the downstream read simulation (*rs*) module.

#### 4.1.3 The count simulation (*cs*) module

The second stage, spatial CNA simulation, further extends allele-specific modeling to simulate new CNA-encoded counts for each allele with the count simulation (*cs*) module and generate new BAM file by allele-resolved UMI resampling with the read simulation (*rs*) module.

The *cs* module constitutes the statistical core of stCNASim. Its mandatory inputs comprise three components: (1) the allele-specific spot-by-gene count matrices (corresponding to alleles A, B, and U), which are derived from seed data and generated by the *afc* module, (2) clonal CNA profiles that define locus-specific copy numbers for both alleles (A and B) within each clone, and (3) clone annotations specifying each clone’s source seed cell type and number of spatial spots to simulate. The module outputs synthetic allele-specific count matrices embedded with clone-specific CNA profiles for all target clones via three key operations. In brief, it first fits gene-wise marginal distributions to allelic count profiles of seed spots; it then translates allele-specific copy number changes into multiplicative scaling factors for distribution mean parameters; finally, it conducts per-gene count simulation from the the adjusted distributions, producing synthetic allele-resolved spot-by-gene count matrices for each simulated clone.

Essentially, the input count data takes the form of an allele × spot × gene tensor, with each spot annotated by its corresponding seed cell type label. Distribution fitting runs independently for every allele and seed cell type, producing gene-wise distribution parameters stratified jointly by allelic identity and cell type. Accordingly, the tensor is partitioned along its three axes into batches of count vectors. Each vector corresponds to a distinct allele, cell type, and gene triplet and contains UMI counts from all spots assigned to the matching cell type. These vectors serve as the fundamental unit for distribution fitting; each is processed in isolation to derive its matched set of gene-wise distribution parameters. In contrast to the top-down tensor partitioning used for distribution fitting, synthetic count generation adopts a bottom-up assembly strategy. During simulation, per-gene count sampling is performed independently for each allele and clone. For a given allele-clone pair, gene-wise counts are sampled from CNA-calibrated distributions to produce a synthetic count vector matching the spot count of the simulated clone. All resulting vectors are then aggregated across all constituent genes, clones, and alleles to assemble the complete synthetic allele × spot × gene tensor, with each spot annotated by its corresponding clone label.

Furthermore, two core independence assumptions underpin the entire count fitting and simulation pipeline: (1) gene independence: marginal distributions are fitted and sampled per gene in isolation, meaning gene-gene correlation and covariance structures are not explicitly modeled. (2) spatial independence: spatial autocorrelation between adjacent spots is omitted from both distribution fitting and count simulation routines. Notably, while the *cs* module is primarily designed for allelic stratified count simulation, it also implements a standalone utility to perform conventional haplotype-agnostic CNA simulation using total gene expression matrices.

The module’s core computational pipeline is detailed below in four sequential steps. We formalize the mathematical notation adopted throughout this module as follows:

- *s* ∈ {1, …, *S*} : Index of seed or simulated spatial spots.
- *g* ∈ {1, …, *G* }: Index of genes.
- *t* ∈ {1, …, *T* }: Index of seed cell types.
- *c*∈ {1, …, *C* }: Index of simulated clones. *t*(*c*) denotes the source seed cell type from which clone *c* inherits distribution parameters.
- *a* ∈ {A, B, U }: Allele channel (A, B, and U).
- **A, B, U** ∈ ℕ^*S×G*^: Spot-by-gene count matrices specific to alleles A, B, and U, respectively.
- **X** = **A** + **B** + **U** ∈ ℕ^*S×G*^: Aggregated total expression count matrix across all three allelic channels.
- **Ã**, 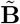, **Ũ**: Simulated allele-specific spot-by-gene count matrices.
- *ℓ*_*s*_: Library size (total UMI count across all genes) of spot *s*.
- *z*_*s*_: Discrete membership label for spot *s*; *z*_*s*_ ∈ {1, …, *T*} for seed cell type assignment; *z*_*s*_ ∈ {1, …, *C*} for simulated clone assignment.
- *n*_*t*_ = ∑_*s*_ I{*z*_*s*_ = *t}*: Total spots belonging to seed cell type *t*.
- *n*_*c*_ = ∑_*s*_ I{*z*_*s*_ = *c}*: Total spots allocated to simulated clone *c*.
- 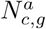 : Absolute copy number of allele *a* at gene *g* within clone *c*, with a diploid baseline of 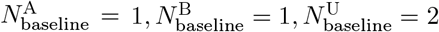
- 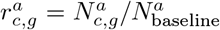: Copy number ratio for allele *a* at gene *g* in clone *c*, normalized against the diploid baseline.

##### Step 1: Data preparation

(1) spot quality control: seed spots are excluded from downstream modeling if their library size < 1000 or the number of detected genes < 100 in the total expression matrix **X**. (2) count matrix partitioning: each allele-specific spot-by-gene count matrix output by the *afc* module is split into cell-type-specific sub-matrices to facilitate gene-wise marginal distribution fitting.

##### Step 2: Marginal distribution fitting

The *cs* module first separately fits cell-type-specific marginal count distributions to gene-wise alleles A, B, and U counts derived from the *afc* outputs. It automatically selects the optimal distribution from four candidate models, including Poisson (Poi), negative binomial (NB), zero-inflated Poisson (ZIP), and zero-inflated negative binomial (ZINB). Its selection strategy mirrors the independent mode implemented in scDesign2 [20], with one key modification: mean parameters are adjusted by library size.

Specifically, for gene *g* in seed cell type *t*, the allelic count 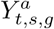 (abbreviated 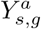 hereafter for simplicity of notation) of allele *a* ∈ {A, B, U} in spot *s* follows one of the four candidate distributions, with spot-specific size-factor-adjusted mean 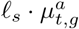, cell-type-specific dispersion 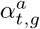 and zero-inflation probability 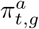:

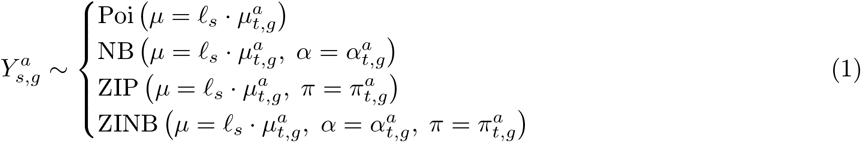

The size factor *ℓ*_*s*_ corresponds to the library size (i.e., total UMIs across all genes) of spot *s*, defined as:

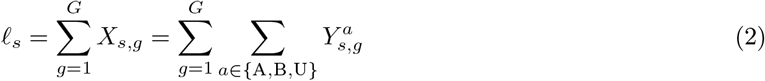

The probability mass function of ZINB, parameterized by mean *µ*, dispersion *α*, and zero inflation *π*, is a mixture of a point mass at zero and a negative binomial distribution:

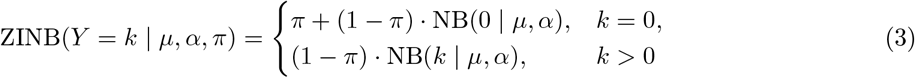

Poi, NB, and ZIP represent constrained special cases of the full ZINB model, formalized as:

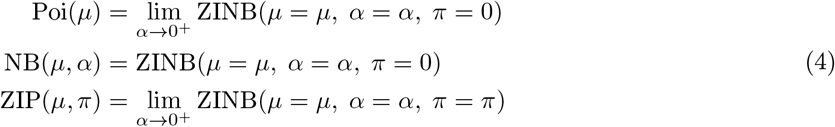

Parameter estimation for all four distributions is supported via dedicated classes available within the Python `statsmodels` package.

A model selection procedure is performed to select the optimal distribution from the four candidate models, leveraging a heuristic strategy and generalized likelihood ratio (GLR) tests.

1. **Objective**: Select the optimal marginal distribution from the four candidate models (Poi, NB, ZIP, ZINB) for the vector of allelic counts 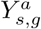 of gene *g* across all *n*_*t*_ spots assigned to cell type *t*.
2. **Dispersion check** : Compute the sample mean 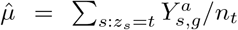 and sample variance 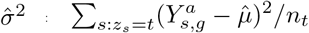.
3. **Underdispersed case** 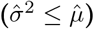: Fit both Poisson (Poi) and zero-inflated Poisson (ZIP) models. Perform a GLR test with 1 degree of freedom to determine if zero-inflation is significant. If the p-value < 0.05, select ZIP; otherwise, select Poisson.
4. **Overdispersed case** 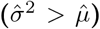: Fit both negative binomial (NB) and zero-inflated negative binomial (ZINB) models. Perform a GLR test with 1 degree of freedom to determine if zero-inflation is significant. If the p-value < 0.05, select ZINB; otherwise, select NB.
5. **Technical issues**: Genes encountering model-fitting failures from technical issues, such as optimization non-convergence, default to the Poisson distribution.
6. **Lowly-expressed genes**: Genes with fewer than `min_nonzero_num` non-zero spots are excluded from fitting and assigned a mean value of zero, treated as transcriptionally silent. Default thresholds are 1 non-zero spot for allele A and B, and 3 for allele U.

##### Step 3: Copy number ratio calculation

CNA states are encoded via allele-specific copy number ratios, where the baseline copy numbers for alleles A, B, and U are set to 1, 1, and 2, respectively. This design enables the simulator to model both total copy number alterations and allele-specific events, such as loss of heterozygosity (LOH). For example, consider an LOH event characterized by loss of allele B, which converts the wild-type AB genotype to a tumor-specific AA genotype; the corresponding allele-specific copy number ratios for alleles A, B, and U are 2, 0, and 1, respectively. More examples of allelic CN ratios can be found in Supplementary Data 4.

Input clonal CNA profiles define segment-level absolute copy numbers for alleles A and B. Per-gene copy ratios are derived by intersecting genomic CNA segments with gene annotation coordinates. Genes with no overlap to any CNA segment retain a copy-neutral ratio of 1.0, while overlapping genes inherit the absolute copy numbers and copy ratios of their respective CNA segments.

For a gene *g* within clone *c* that overlaps a CNA segment and bears allelic copy numbers 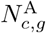 (allele A) and 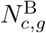 (allele B) inherited from that segment, the copy number 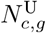 of allele U is the sum of the two haplotypes:

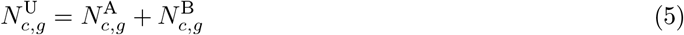

Consequently, the bounded copy ratio 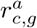 for each allele *a* ∈ {A, B, U} of gene *g* in clone *c* is given by:

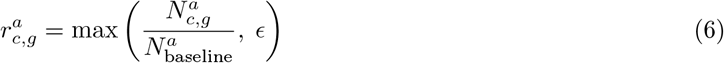

where the diploid baselines for the three alleles are 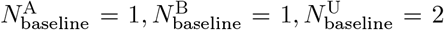. The leakage rate *ϵ* (default: 0.01) establishes a non-zero frequency floor for completely deleted alleles, accounting for real-world sequencing errors and mapping artifacts that produce background reads.

##### Step 4: CNA-modulated count simulation

Gene-wise baseline allelic means obtained from distribution fitting are rescaled using allele- and clone-specific copy ratios to adjust the marginal distributions, and synthetic allelic counts are subsequently sampled from these updated distributions. Ultimately, the *cs* module outputs three separate spot-by-gene count matrices for alleles A, B, and U, encoded with the user-specified CNA profiles and clonal structures.

##### Size factor simulation

Library size distributions are first characterized from the seed spots and subsequently used to sample new library sizes for the synthetic spots. For seed cell type *t*, library size *ℓ*_*s*_ of spot *s* follows a log-normal distribution with shift 1:

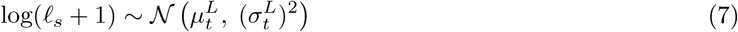

For clone *c* originating from source seed cell type *t*(*c*), *n*_*c*_ synthetic library sizes 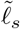 are then sampled from the fitted distribution corresponding to *t*(*c*):

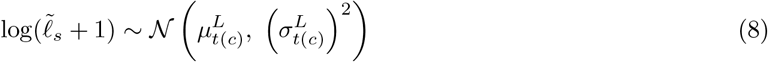

Sampled library sizes are clamped to a bounded range [*L*_min_, *L*_max_] to prevent unrealistic outliers, where 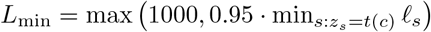 and 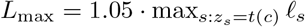.

##### CNA-induced parameter rescaling

The fitted baseline allelic mean of each gene is scaled by two multiplicative factors: the allele-specific copy ratio 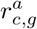 and a global sequencing coverage scaling term *ρ*. For clone *c* derived from source cell type *t*(*c*), the rescaled mean 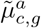 for gene *g* is:

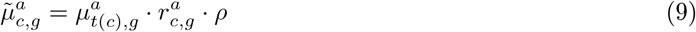

Here *ρ* (default 1.0) modulates the ratio between the average library size of all synthetic spots and that of the original seed data. Dispersion (*α*) and zero-inflation (*π*) parameters are inherited unchanged from the source cell type, reflecting the assumption that CNA events alter the mean expression level but not the dispersion and zero-inflation structure of the gene.

##### Gene-wise count sampling

Within each clone, synthetic allelic counts are sampled independently per gene from the rescaled ZINB distribution, modulated by spot-specific simulated library sizes. For a synthetic spot *s* assigned to clone *c* with library size 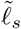, the simulated allelic count 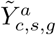 (abbreviated 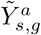 hereafter for simplicity of notation) for allele *a* at gene *g* is sampled as:

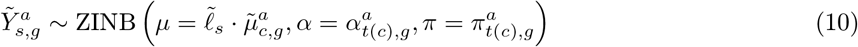

Unified ZINB sampling is universally applicable even when the gene’s optimal fitted model is Poisson, NB, or ZIP, as all three simpler distributions represent constrained ZINB sub-cases.

##### Output assembly

Full-length allelic spot-by-gene count matrices are assembled via vertical concatenation of all clone-level simulated matrices. All synthetic count matrices corresponding to alleles A, B, and U are finally integrated into one unified AnnData object [27], annotated with clone identity labels.

#### 4.1.4 The read simulation (*rs*) module

The *rs* module generates synthetic allele-resolved sequencing reads stored in standard BAM format, requiring two inputs: (1) per-gene, per-allele partitioned seed BAM files produced by the *afc* module; and (2) the three allele-specific simulated count matrices output from the *cs* module. Rather than generating reads *de novo*, which would not preserve realistic characteristics such as base quality scores and fragment length distributions, the module performs resampling of original UMI read groups extracted from the seed BAM files. For every gene-allele pair, seed UMIs are sampled with replacement to match the target simulated UMI abundance defined in the *cs* output matrices. Sampled reads are SNP-masked to enforce haplotype consistency, and re-barcoded with synthetic spot identifiers and UMI sequences before final BAM compilation.

##### Step 1: Data preparation

Unique barcodes are allocated to simulated spots via random generation or sampling from a user-supplied whitelist. Genes are split into batches for parallel processing.

##### Step 2: Per-gene UMI resampling with haplotype SNP masking

For every gene-allele pair, seed UMI read groups are sampled with replacement from the corresponding partitioned BAM file, until the resulting per-spot UMI counts align with the target values defined in the count matrices simulated by the *cs* module. If the target count exceeds the number of available seed UMIs, the same seed UMI may be sampled multiple times - each replicate will receive distinct new barcodes and a replicate index suffix in the read query name. All reads associated with each resampled seed UMI undergo three post-processing steps.

- **Haplotype SNP masking**. To enforce haplotype consistency between simulated reads and the original UMI-level consensus haplotype, we adjust base calls across all aligned positions of each read: at SNP positions (all post-QC phased SNPs covered by the read and identified by the *afc* module), the bases are replaced with the SNP-alleles matching the target haplotype to resolve technical artifacts including sequencing errors. At all other read positions, bases are overwritten using reference sequences retrieved from the reference FASTA file. This adjustment reduces noisy genotyping signals that would otherwise distort haplotype inference when CNA detection tools perform reference phasing on the simulated BAM.
- **BAM tag refresh**. Each resampled seed UMI read group is assigned a new unique spot barcode and UMI sequence, with UMIs constrained to be distinct within each synthetic spot. The read tags storing spot barcodes (`CB` ) and UMI sequences (`UB` ) are first archived in auxiliary backup tags for complete read provenance tracking, then overwritten with the synthetic identifiers. Additionally, the raw uncorrected barcode tags (`CR` and `UR` ) are updated to align with the newly generated barcode sequences.
- **Read query name modification**. The read query name is appended with a gene index and replicate index suffix to ensure global uniqueness when the same seed UMI is sampled multiple times.

##### Step 3: Allele-specific BAM assembly

All per-gene per-allele synthetic BAM files are concatenated, coordinate-sorted and indexed to generate the final output BAM file. The resulting simulated BAM file is designed to preserve the technical characteristics of the seed sequencing data, including (1) CIGAR strings and aligned positions that partially recapitulate gene exon-intron structures; (2) base quality scores; (3) fragment length distributions and (4) UMI architectures - encompassing the read counts and distribution pattern along the gene locus.

#### 4.1.5 The spatial patterning module

In addition to its four core modules (*pp, afc, cs*, and *rs*), the simulator features an auxiliary spatial patterning module that generates structured spatial tumor architectures. Users first specify a spatial distribution signature, including clonal shape, inter-clonal distance and intermixing rate, for each simulated clone, after which the tool assigns synthetic spots to spatial coordinates within each clonal compartment. Molecular identities (barcodes, clone labels, expression and BAM content) remain fixed across spatial conditions, and only the spot coordinates are altered. Notably, the four core modules perform count and read simulation without incorporating spatial information, whereas spatial structure is appended as a post-processing step via this dedicated module. This modular separation implies that the simulator does not explicitly model spatial autocorrelation in gene expression values or sequencing read features.

##### Coordinate lattice and clone assignment

Spatial layouts are constructed on a Visium-like odd-*r* hexagonal lattice with fixed nearest-neighbor spacing *s* = 1. For row *r* and column *c*, spot coordinates *x* and *y* are

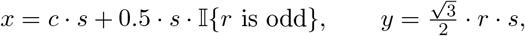

after which coordinates are centered at the origin. Users specify a spatial signature through three controllable factors: clonal geometry, interclonal distance and interclonal mixing rate. Clone labels are assigned by geometric scores on this lattice under exact cardinality constraints; barcodes within each clone type are then randomly permuted and mapped one-to-one onto the labeled spots. Outputs follow a Space Ranger– compatible format (spot coordinates and spot-level clone annotations), enabling direct use with spatial transcriptomics analysis tools.

### 4.2 Seed data generation

All simulated datasets analyzed in this study were derived from a single unified seed dataset, which was built from the 10x Visium spatial transcriptomics data of HCC-3N - a normal tissue section isolated from the HCC-3 hepatocellular carcinoma (HCC) specimen reported in [23].

#### 4.2.1 Seed spots selection and BAM file generation

The raw HCC-3N dataset contained 4,289 spatially resolved in-tissue spots covering the entire tissue section. Raw sequencing reads were preprocessed with SpaceRanger v1.1.0 and aligned to the GRCh38 human reference genome. For seed data construction, we first computed per-spot quality control metrics, including the number of detected genes, total UMI counts, and the fraction of mitochondrial UMIs, using the function `scanpy.pp.calculate_qc_metrics()` . Spots with poor quality were excluded according to the following criteria: `n_genes_by_counts` < 1000 or > 6000, `total_counts` < 2000 or > 35000, or `pct_counts_mt` > 20. After filtering, a spatially contiguous panel of 600 high-quality normal spots were selected as seed spots for downstream simulation and benchmarking. The corresponding BAM file and gene-by-spot expression count matrix were subsequently subset to include solely the 600 selected spots; BAM file subset was performed using samtools v1.23.1 [28].

#### 4.2.2 Phased SNPs generation

Phased SNP sets were derived from the BAM file of the 4,289-spot HCC-3N 10× Visium dataset [23] using a reference-based phasing pipeline wrapped within the xcltk package [12]. Specifically, genotyping of around 7M common SNPs in the BAM file was first performed with cellsnp-lite [29], configured with the parameters `–minCOUNT 11` and `–minMAF 0.1` . The resulting SNP genotypes were then subjected to statistical reference phasing via Eagle2 [30] to produce the final phased SNP panel. Note that only autosomal SNPs on chromosomes 1-22 were retained throughout this workflow.

### 4.3 Simulating the validation and benchmarking datasets using stCNASim

All synthetic datasets generated by stCNASim, including data for simulator validation and CNA inference benchmarking, were built upon the unified seed dataset described in Section 4.2. Every simulation leveraged the GRCh38 human reference genome, paired with gene annotations derived from the CellRanger-provided GTF file (version `GRCh38-3.0.0` ). Genes with invalid or duplicate names, alongside those mapping outside autosomes 1-22, were excluded from downstream analysis. This filtering workflow retained a final panel of 32,295 genes from the original 33,539 annotated entries. With the seed dataset, GRCh38 reference genome, and curated gene annotations serving as core inputs, stCNASim generates tailored synthetic spatial tran-scriptomic datasets configured with task-specific CNA profiles and clonal architectures. The complete list of all simulated datasets utilized in this study can be found in Supplementary Data 5, and their corresponding ground-truth CNA profiles applied during simulation are provided in Supplementary Data 6.

### 4.4 Multi-metric evaluation of stCNASim

#### 4.4.1 Calculation of RDR-related metrics

The RDR-related spot-wise and gene-wise metrics for simulator evaluation were computed on both total-expression and allele-resolved count matrices, stratified by cell types (seed data) or clone labels (simulated data). We denote an evaluated spot-by-gene count matrix as **Y**^*S×G*^, with entry *Y*_*s,g*_ corresponding to spot *s* ∈ {1, …, *S}* and gene *g* ∈ {1, …, *G}*.

##### Spot-wise metrics

For each spot *s*, the library size is defined as 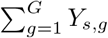, and the gene zero proportion equals 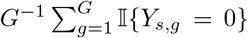. Spot-level metrics were calculated exclusively for total-expression matrices and omitted from allele-stratified count matrices.

##### Gene-wise metrics

For each gene *g*, we calculated four statistics - Mean: 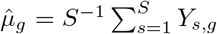; Variance: 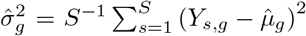; Coefficient of variation (CV): 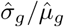, and CV is set to 0 when 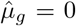 Spot zero proportion: 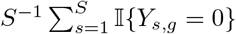.

##### Smoothed pairwise gene-statistic relationships

We generated continuous smoothed curves to visualize the association between the three gene-wise statistics (variance, CV, zero proportion) versus genewise mean expression. Smoothing was implemented via k-nearest-neighbor kernel regression (default *k* = 100) with Gaussian distance weighting.

##### Pseudo-bulk log2 fold change (log2FC) between two groups

Consider two groups with identically sized count matrices **Y**^(1)^ and **Y**^(2)^, where 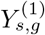 and 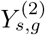 represent counts for spot *s* and gene *g* in each cohort. Let 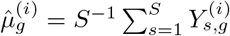 denote the mean expression of gene *g* in group *i* ∈ {1, 2}. The log2FC of group 2 relative to group 1 is defined piecewise:

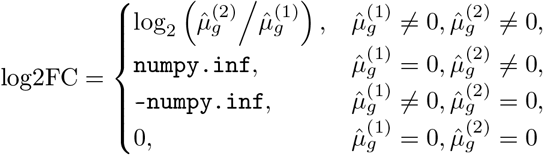

#### 4.4.2 Calculation of BAF-related metrics

The BAF-related gene-wise and SNP-wise metrics were calculated for total allelic depth (DP) count matrices **D** = **A** + **B**, where **A** and **B** store UMI counts for the two phased haplotypes A and B. These gene-level and SNP-level DP matrices were stratified by cell types (seed data) or clone labels (simulated data).

##### Gene-wise metrics

Let **D**^*S×G*^ be a spot-by-gene DP matrix with entry *D*_*s,g*_ for spot *s* ∈ {1, …, *S}* and gene *g* ∈ {1, …, *G}*. For each gene *g*, we calculated three statistics - DP mean: 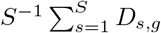; DP spot zero proportion: 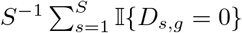; Phased allele frequency 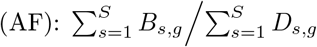.

##### SNP-wise metrics

Let **D**^*S×P*^ denote the spot-by-SNP depth matrix with entry *D*_*s,p*_ for spot *s* ∈ {1, …, *S}* and SNP *p* ∈ {1, …, *P}*. For each SNP *p*, we calculated four statistics - DP mean: 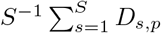 DP spot zero proportion: 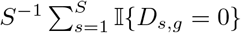; Phased AF: 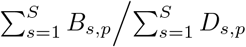. Raw unphased AF: 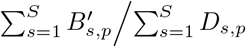 where 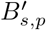 corresponds to raw UMI count of the unphased alternative allele (i.e., the `ALT` field in the VCF file) of SNP *p* in spot *s*.

#### 4.4.3 Median library size normalization for total-expression matrices

For a total-expression count matrix, its normalization by median library size was performed with the Scanpy function `scanpy.pp.normalize_total(target_sum = median_library_size)`, where `median_library_size` is the median spot library size calculated across the full input matrix.

### 4.5 Configuration and parameters of CNA inference methods

Throughout this benchmarking work, we evaluated five established CNA detection tools: InferCNV v1.18.1, CopyKAT v1.1.0, Numbat v1.4.2, XClone v0.4.2, and CalicoST v1.0.0. Their specific configurations and key parameters are detailed below:

#### InferCNV

We executed the R function `infercnv::run()` with the core parameters set to `cutoff=0.1, cluster_by_groups=TRUE, denoise=TRUE, HMM=TRUE` . When incorporating reference spots, we supplied the reference cell types or clone labels to the `ref_group_names` parameter in the `CreateInfercnvObject()` function; otherwise, the parameter was set to `NULL` . For datasets containing subclonal structures, we evaluated InferCNV’s within-group clustering against global clustering by setting `cluster_by_groups=TRUE` and `cluster_by_groups=FALSE`, respectively. For the spatial patterning experiments, InferCNV was run once with known normal references and `cluster_by_groups=TRUE`, and the HMM-based discrete CN-state calling step was disabled (`HMM=FALSE` ), retaining the denoised continuous expression residual matrix used for AUROC/AUPRC scoring.

#### CopyKAT

We ran the R function `copykat::copykat()` using the default parameters `id.type=` “`S`”`, ngene.chr=5, win.size=25, KS.cut=0.1, distance=` “`euclidean`”. Barcodes corresponding to diploid reference spots were passed to the `norm.cell.names` parameter for reference-aided runs; this argument was omitted for reference-free analysis.

#### Numba

We first performed preprocessing according to the tool’s user guide, using the pipeline wrapped in the pre-compiled `pileup_and_phase.R` script to extract allelic signals. Subsequently, we executed the R function `numbat::run_numbat()` for CNA inference, using the key parameters `t=1e-5, gamma=20` . Following the tool’s GitHub documentation, reference spots were explicitly excluded prior to executing the core inference routine. For the spatial patterning baseline, we used no-filter preprocessing that retains normal spots in the allele and count matrices, aggregates them into a reference expression profile, and then runs `run_numbat()` without stripping those spots from the input matrices.

#### XClone

Preprocessing to generate spot-by-gene total-expression and allelic count matrices was performed via the recommended `xcltk` Python package. We then executed the tool’s RDR, BAF, and Combine modules using `xclone.model.run_RDR()`, `xclone.model.run_BAF()`, and `xclone.model.run_combine()`, respectively, setting `clustering_method=` “`baf`” in the Combine module. For the dataset with mirrored copy loss events, we set `merge_loss=False, customizedplotting=True` to visualize the two distinct alleles. Similarly, we configured `merge_loh=False, customizedplotting=True` for the dataset featuring mirrored LOH events.

#### CalicoST

We evaluated CalicoST across four distinct configurations based on the utilization of reference spots and the activation of the spot-level tumor purity estimation step: [ref(+)/purity(+)], [ref(+)/purity(-)], [ref(-)/purity(+)], and [ref(-)/purity(-)]. When reference spots were used (i.e., in the [ref(+)/purity(+)] and [ref(+)/purity(-)] modes), the file containing the reference barcodes was passed to the `normalidx_file` parameter; otherwise, it was set to `None` . For the main spatial patterning evaluation, CalicoST was run in the [ref(-)/purity(-)] configuration (`normalidx_file=None`, `tumorprop_file=None`, `tumorprop_threshold=0.001` ), consistent with homogeneous simulated spots and to avoid purity-step failures on unstructured layouts. All other parameters followed the user guide available in its GitHub repository (https://github.com/raphael-group/CalicoST).

### 4.6 Multi-task benchmarking framework for CNA inference methods

We implemented a comprehensive benchmarking pipeline (`bcd` package) to systematically evaluate five copy number aberration (CNA) detection methods, InferCNV [3], CopyKAT [9], Numbat [11], XClone [12], and CalicoST [16], across three complementary evaluation tasks: (1) genome-wide CNA profile prediction, (2) tumor versus normal cell classification, and (3) subclonal structure identification. Each task follows a consistent workflow: tool-specific data extraction and preprocessing, ground-truth formatting, spot/gene overlap harmonization, and metric computation. Reference (normal) spots, as specified by user-provided labels, were excluded from all evaluation steps prior to metric calculation unless otherwise specified.

#### 4.6.1 Benchmarking CNA profile prediction

The performance of each tool for CNA profile prediction was evaluated independently across three categories: copy gain, copy loss, and LOH. For each tool and CNA type, we adopted the Area Under the Precision-Recall Curve (AUPRC) and Area Under the Receiver Operating Characteristic curve (AUROC) as evaluation metrics. These metrics were computed by comparing each tool’s predictions against ground-truth profiles. Ground truths consist of predefined *in silico* CNA states for simulated datasets, or profiles inferred from matched omics data for real biological samples.

##### Step 1: Tool-specific data extraction and preprocessing

Because each method generates predictions with distinct file formats and genomic resolutions, we extracted and standardized these outputs into unified spot-by-gene matrices.

- **InferCNV**: Raw output consisted of a continuous spot-by-gene matrix of smoothed relative expression values normalized against a diploid reference baseline. Deviations from this baseline (values above or below 1) indicate copy number gains or losses. The same matrix was reused for all CNA types. Standardization: expression values were sign-flipped when evaluating copy loss.
- **CopyKAT**: Raw output was a continuous spot-by-gene matrix containing log-scale relative expression values against a diploid baseline. Deviations from the baseline (values above or below 0) indicate evidence of gains or losses. This single matrix was shared across all CNA types. Standardization: for copy-loss evaluation, values were sign-inverted (multiplied by -1) so that higher scores represent stronger evidence of loss.
- **Numbat**: Raw outputs comprised spot-by-segment posterior probability matrices for each CNA type. Standardization: genomic segments were mapped to gene-level coordinates using gene annotation files to yield per-CNA-type, spot-by-gene probability matrices. Unlike other tools, Numbat only returns results for segments where CNAs were detected and excludes all other segments. To address this discrepancy, we expanded Numbat’s spot-by-gene probability matrices to the full transcriptome-wide gene set and filled missing entries with a value of 0 for genes residing within unreported segments.
- **XClone**: Raw outputs contained spot-by-gene CNA posterior probabilities, providing separate matrices for each CNA category (gain, loss, and LOH).
- **CalicoST**: Raw outputs included clonal gene copy number estimates alongside spot-level clone labels and tumor proportions. Standardization: for CalicoST configurations that do not output tumor fractions, clone 0 was assigned a fraction of 0.0, and all other detected clones received 1.0. Per-entry CNA probabilities within the final spot-by-gene matrix were computed by multiplying deterministic CNA states (binary values of 0 or 1 derived from clonal gene-level copy number calls) by the corresponding per-spot tumor fractions, yielding separate matrices for gain, loss, and LOH.

##### Step 2: Ground truth construction

Ground-truth CNA states were initially defined from segment-level CNA annotations. These segment-level annotations were further projected to the gene level by intersecting genomic segments with annotated gene coordinates. Specifically, genes overlapping a CNA segment inherited the corresponding segment CNA state, whereas non-overlapping genes were assigned a copy-neutral state. Genes that overlap multiple distinct segments were excluded from downstream analysis. This procedure yielded binary spot-by-gene matrices, in which a value of 1 denotes the presence of a CNA event and 0 indicates a copy-neutral state. Independent binary matrices were constructed separately for gain, loss, and LOH events.

##### Step 3: Harmonization

The following harmonization procedures were performed independently for each CNA type (gain, loss, and LOH): (1) All reference spots were removed from both tool-generated and ground-truth matrices. (2) Remaining spots were intersected across all tools and ground-truth data, while genes were pairwise intersected between each individual tool and ground-truth profile. (3) Each tool matrix and its matched ground-truth matrix was subset to the overlapping spots and genes, ensuring identical matrix dimensions for rigorous element-wise comparison.

##### Step 4: Evaluation metrics calculation

Evaluation metrics were calculated independently for the three CNA categories: copy gain, copy loss, and LOH. Threshold-dependent binary classification metrics were computed on flattened matrices specific to each CNA type. We sampled up to 1,000 distinct threshold cutoffs: predictions with scores greater than or equal to a given cutoff were classified as positive, and all remaining cases were classified as negative. AUPRC was derived from precision and recall values across all sampled thresholds. Similarly, AUROC was calculated based on corresponding true positive rates (TPR) and false positive rates (FPR).

#### 4.6.2 Benchmarking tumor/normal classification

To evaluate the performance of each tool in binary classification of tumor versus normal spots, we used five evaluation metrics: accuracy, precision, recall, F1-score, and the adjusted Rand index (ARI). These metrics were calculated by comparing classification outputs from each tool against ground-truth labels. Ground truth labels were predefined *in silico* labels for simulated datasets, or derived from well-validated annotations for real biological samples.

##### Step 1: Tool-specific data extraction and classification logic

For each tool, classification results were extracted directly when available; otherwise, raw outputs were converted into binary tumor/normal assignments. A standardization procedure was implemented to harmonize the data format across all tools. All reference spots were removed from raw outputs prior to standardization.

- **InferCNV**: The tool does not directly output tumor labels. Standardization: Hierarchical clustering with Ward’s criterion (ward.D2) and Euclidean distance was applied to the spot-by-gene relative expression matrix. The `cutree` function was then used to partition the spots into two clusters. Within each cluster, per-spot CNA scores were calculated as the mean absolute deviation (MAD) of all genes relative to a reference expression baseline of 1.0. Finally, the cluster with the higher mean CNA score was designated as the tumor clone, while the remaining cluster was identified as the normal clone.
- **CopyKAT**: Raw outputs correspond to per-spot ploidy calls. Standardization: spots labeled aneuploid were classified as tumor, and diploid spots were classified as normal.
- **Numbat**: We directly adopted the tool’s native per-spot binary tumor/normal assignments.
- **XClone**: We directly adopted the tool’s native per-spot binary tumor/normal assignments.
- **CalicoST**: Raw outputs consist of per-spot tumor proportions. Standardization: for CalicoST configurations that do not report tumor fractions, clone 0 was assigned a tumor fraction of 0.0, and all other identified clones were assigned 1.0. K-means clustering (*k* = 2) was applied to tumor proportion values to partition all spots into two clusters. The cluster with a higher mean tumor proportion was classified as the tumor clone, and the other cluster as the normal clone.

##### Step 2: Ground truth construction

Clone-level ground-truth tumor/normal labels were mapped to spot-level labels according to per-spot clone assignments. All reference spots were excluded from the evaluation set.

##### Step 3: Harmonization

Predicted labels from each tool and ground-truth labels were subset to the intersection of spots across all tools and the ground truth.

##### Step 4: Evaluation metrics calculation

We computed the five metrics by comparing predicted labels against ground truth, with tumor defined as the positive class. TP: true positive; FP: false positive; FN: false negative; *N* : total number of evaluated spots. Accuracy = (TP + TN)*/N* ; Precision = TP*/*(TP + FP); Recall = TP*/*(TP+ FN); F1-score = 2. (Precision .Recall)*/*(Precision + Recall). ARI: measures the similarity between predicted and true cluster assignments, adjusted for chance.

#### 4.6.3 Benchmarking subclonal structure identification

To evaluate each tool’s performance in subclonal structure identification, we adopted the adjusted Rand index (ARI) as the evaluation metric. This metric was computed by comparing each tool’s subclone identification outputs against ground-truth labels.

##### Step 1: Tool-specific data extraction and subclone identification

Let *k* denote the number of ground-truth subclones among the evaluated spots. For each tool, per-spot clone labels were extracted directly where available. When explicit labels were absent, raw outputs were converted to *k* clone assignments. A standardization workflow was implemented to harmonize data formats across all tools, and all reference spots were excluded from downstream evaluation.

- **InferCNV**: Standardization: Hierarchical clustering with Ward’s criterion (ward.D2) and Euclidean distance was performed on the spot-by-gene relative expression matrix. The `cutree` function was subsequently used to partition evaluated spots into *k* clusters to generate per-spot clone assignments. While InferCNV supports native global subclustering across all evaluated spots under the setting `cluster_by_groups=FALSE` (e.g., via built-in hierarchical or Leiden clustering on the relative expression matrix), the post-hoc clustering above adopts matching configurations to its internal hierarchical clustering. This post-hoc clustering was initially implemented to obtain subclustering results for the `cluster_by_groups=TRUE` setting, which is optimized for CNA profile visualization.
- **CopyKAT**: Raw outputs contained hierarchical clustering results. Standardization: The `cutree` operation was applied to partition evaluated spots into *k* clusters.
- **Numbat**: Raw outputs included per-spot clone labels.
- **XClone**: Raw outputs included per-spot clone labels.
- **CalicoST**: Raw outputs included per-spot clone labels.

##### Step 2: Ground truth

Ground-truth clone labels correspond to predefined *in silico* assignments for simulated datasets, or well-validated clone annotations for real biological samples. All reference spots were excluded from the evaluation set.

##### Step 3: Harmonization

Predicted labels from each tool and ground-truth labels were subset to the shared set of spots present across all tools and the ground truth.

##### Step 4: Evaluation metrics calculation

The primary evaluation metric is the adjusted Rand index (ARI), which quantifies the agreement between predicted clone assignments and ground-truth labels, corrected for chance. The ARI ranges from -1 to 1, where higher values indicate stronger similarity.

### 4.7 Stress-testing spatial CNA prediction using stCNASim-synthetic datasets

The data generation process has been described in Section 2.3 and Supplementary Notes. Additional details for specific datasets are provided below.

#### 4.7.1 Construction of datasets spanning a gradient of tumor purity

The workflow used to generate nine synthetic datasets covering tumor proportions ranging from 1% to 99% was visualized in Supplementary Fig. S19a-b. Specifically, we first constructed a base synthetic dataset using stCNASim, comprising 2,600 spatial spots: 600 reference spots, 1,000 normal spots, and 1,000 tumor spots carrying predefined clonal CNA signatures. We then subsampled normal and tumor spots without replacement from their corresponding clonal populations within the base dataset to generate distinct purity levels. Across all derived datasets, the count of reference spots was held fixed at 600, and the aggregate number of normal plus tumor spots was maintained at 1,000. The number of sampled tumor spots varied incrementally from 10 to 990, establishing a continuous gradient of tumor purity for downstream benchmarking.

#### 4.7.2 HCC scRNA-seq data processing

To assess whether external single-cell RNA-seq (scRNA-seq) data can serve as a reference for CNA analysis in spatial transcriptomics, we leveraged a hepatocellular carcinoma (HCC) scRNA-seq dataset reported by Lu et al. [25]. We isolated the “HCC-03N” normal sample from the raw count matrix to produce a 2601 × 25712 cell-by-gene expression matrix. We then computed per-cell quality control (QC) metrics, including number of detected genes, total UMI counts, and the proportion of mitochondrial UMIs, using the Scanpy function `scanpy.pp.calculate_qc_metrics()` . Cells failing QC were filtered out according to the following cutoffs: `n_genes_by_counts` < 1000 or > 3000, `total_counts` < 2000 or > 30000, or `pct_counts_mt` > 20. This filtering retained 655 high-quality cells. We next excluded genes with duplicated symbols and kept only those shared with our synthetic spatial transcriptomics dataset (at 90% tumor purity). After gene filtering, 19,228 unique genes remained. We then combined barcodes from the 655 single cells and 1000 spatial spots (100 normal, 900 tumor) into a unified count matrix, resulting in a final 1665 × 19228 barcode-by-gene count matrix.

Using this combined dataset, we supplied the 655 single-cell hepatocytes as a reference to infer CNAs across all 1000 spatial spots via InferCNV. We additionally constructed a second dataset by random sub-sampling from the above matrix: 600 single-cell hepatocytes, 20 normal spots, and 180 tumor spots. This yielded a new combined 800 × 19228 barcode-by-gene count matrix. For this second experiment, we ran InferCNV on all cells and spots jointly to predict CNAs without designating a reference population.

#### 4.7.3 Reference spot downsampling experiment

We first built a base synthetic dataset via stCNASim, consisting of 600 reference spots alongside 1,000 tumor spots encoded with predefined clonal CNA profiles. We then generated five additional datasets by subsampling reference spots without replacement from this base cohort to produce a gradient of reduced reference counts. For all downsampled datasets, the number of tumor spots was fixed at 1,000, while the retained reference spot count was sequentially reduced from 600 down to 3.

#### 4.7.4 Coverage downsampling experiment

The base dataset was generated by stCNASim and contained 600 reference spots plus 1,000 tumor spots carrying predefined clonal CNA profiles. We subsampled sequencing reads from the base BAM file using the command `samtools view -h -b -s <fraction>`, yielding five additional datasets with progressively reduced read coverage fractions.

### 4.8 Benchmarking the impact of cell spatial cell patterns on CNA calling

#### 4.8.1 Dataset generation

Using identical seed data derived from the HCC-3 dataset, we employed stCNASim to generate a simulated cohort comprising normal cells and two tumor clones (tumor1 and tumor2). Clones tumor1 and tumor2 share three CNA events: a copy-number gain on chr1q, a copy-number loss on chr8p and a copy-number gain on chr8q. In addition, tumor2 harbors three private events: a copy-number loss on chr4q, LOH on chr13q and LOH on chr17p. We initially simulated 1000 cells of each type. To construct spatial patterns with unbalanced tumor compartments while retaining a large normal background, tumor cells were randomly downsampled to 300 tumor1 and 700 tumor2 cells (seed 12345); all 1000 normal cells were retained. The resulting 2000-cell molecular dataset—including expression counts, BAM content and clone identities—was held fixed across all spatial conditions below.

Because the core simulation stages of stCNASim do not emit spatial coordinates, spot positions were assigned with the spatial patterning module (Section 4.1.5). Cells were placed on a Visium-like hexagonal lattice (nearest-neighbor spacing of 1 arbitrary unit) such that each barcode mapped to a unique spot within its clonal compartment. Only coordinates changed between conditions; this design isolates the effect of spatial architecture on downstream CNA calling. We evaluated three scenarios: varying clonal geometries, varying interclonal distance and varying interclonal cell mixing.

##### Varying clonal shapes

Six architectures were generated under identical molecular composition:

1. **Separated clusters:** tumor1 and tumor2 form two well-separated compact compartments surrounded by normal cells.
2. **Mixed:** normal and tumor cells are randomly distributed across the slide with no spatial structure.
3. **Ring:** tumor clones occupy concentric annular compartments, with tumor1 enriched at a smaller radius than tumor2 and normal cells filling the remainder of the lattice.
4. **Stripes:** tumor1 and tumor2 form parallel horizontal stripe-shaped compartments adjacent across a midline.
5. **Intermixing:** two predominant left/right tumor compartments (mainly tumor1 and tumor2, respectively) with a minority of each type infiltrating the opposite compartment.
6. **Single tumor region:** tumor1 and tumor2 occupy a single contiguous tumor domain, with a smaller tumor1 pocket adjacent to a larger tumor2 region.

##### Varying interclonal distance

Holding stripe geometry fixed, we generated five configurations in which the center-to-center gap between the two tumor stripes increased over 0, 9.0, 18.1, 27.1 and 36.1 lattice units (spot spacing = 1). At gap 0 the stripes abut across the midline; larger gaps insert an expanding band of intervening normal spots.

##### Varying interclonal cell mixing rate

Six configurations with left/right tumor compartments were generated under progressively stronger cross-compartment infiltration. The mixing rate *r* is defined as the fraction of each tumor type relocated into the opposite clone’s primary compartment (round(*r* · 300) tumor1 spots in the tumor2 region and round(*r* · 700) tumor2 spots in the tumor1 region). We used *r* ∈ {0, 0.08, 0.16, 0.24, 0.32, 0.40}.

#### 4.8.2 Running CNA analysis tools on simulated datasets

Because InferCNV, CopyKAT, Numbat and XClone do not incorporate spatial information, each was run once on the fixed 2000-cell molecular cohort to provide a non-spatial performance baseline. Gene-expression counts for InferCNV, CopyKAT and Numbat were obtained from the shared starsolo-derived count matrix (exported to 10x Genomics format); XClone used allele-aware read-depth (RDR) and B-allele frequency (BAF) features generated from the simulated BAM with xcltk, and Numbat additionally used BAM-derived allele pileups and phasing. Analyses followed each tool’s standard pipeline with the 1000 normal cells designated as the reference. Reported XClone metrics used BAF-based clone clustering.

For CalicoST, each spatial condition was analyzed from the simulated BAM together with a Space Ranger–style directory containing the shared expression matrix and pattern-specific spatial coordinates. Tumor-proportion (purity) estimation was omitted in the final benchmarking runs by setting `tumorprop_file` to `None`, consistent with the simulation design in which each spot is purely normal or purely tumor, and to avoid purity-step failures observed for unstructured or otherwise problematic layouts. When purity was bypassed, `tumorprop_threshold` was set to 0.001; remaining CalicoST settings followed the recommended configuration.

## 5 Data availability

The 10x Visium spatial transcriptomics data of human hepatocellular carcinoma (HCC) [23] were downloaded from Genome Sequence Archive (GSA) under accession number HRA000437, encompassing three sample subsets: HCC-3N, HCC-3L, and HCC-3T. The scRNA-seq profiles of human HCC from a separate study [25] were downloaded from GEO under accession number GSE149614.

## 6 Code availability

The stCNASim simulator presented in this manuscript can be accessed at the GitHub repository https://github.com/hxj5/stCNASim. All codes and associated materials required to reproduce the manuscript’s analyses are available at https://github.com/hxj5/bcd.

## 7 Acknowledgements

We thank Prof. Jin Gu for sharing the HCC data, and the Centre for PanorOmic Sciences (CPOS) at HKU for providing computing facilities. This project is supported by the National Natural Science Foundation of China (No. 62222217), the Research Grants Council of the Hong Kong SAR, China (No. 17126725), the InnoHK initiative of the Innovation and Technology Commission of the Hong Kong Special Administrative Region Government, and the University of Hong Kong (a startup fund and a seed fund, Y.H.; a Presidential PhD Scholarship, J.Q.). R. H. is supported by a Cancer Research Institute Immuno-Informatics Postdoctoral Fellowship (CRI Award 14614).

## 8 Contributions

Y.H. conceived the study, and R.H. initialized the analyses. X.H. implemented the allele-aware spatial RNA-seq CNA simulator stCNASim and performed all simulator validation assessments. X.H. and J.Q. designed the multi-task benchmarking pipeline for spatial CNA inference. X.H. carried out all benchmarking analyses, with spatial patterning experiments led independently by J.Q.. X.H., Y.H. and J.Q. wrote the manuscript with input from R.H.. All authors read, commented on, and approved the final manuscript.

## Notes

### Competing Interest Statement

The authors have declared no competing interest.

