## Supplementary Information for "stCNASim: Allele-aware spatial RNA-seq simulator enables systematic benchmarking of copy number inference"

### 1 Supplementary Figures

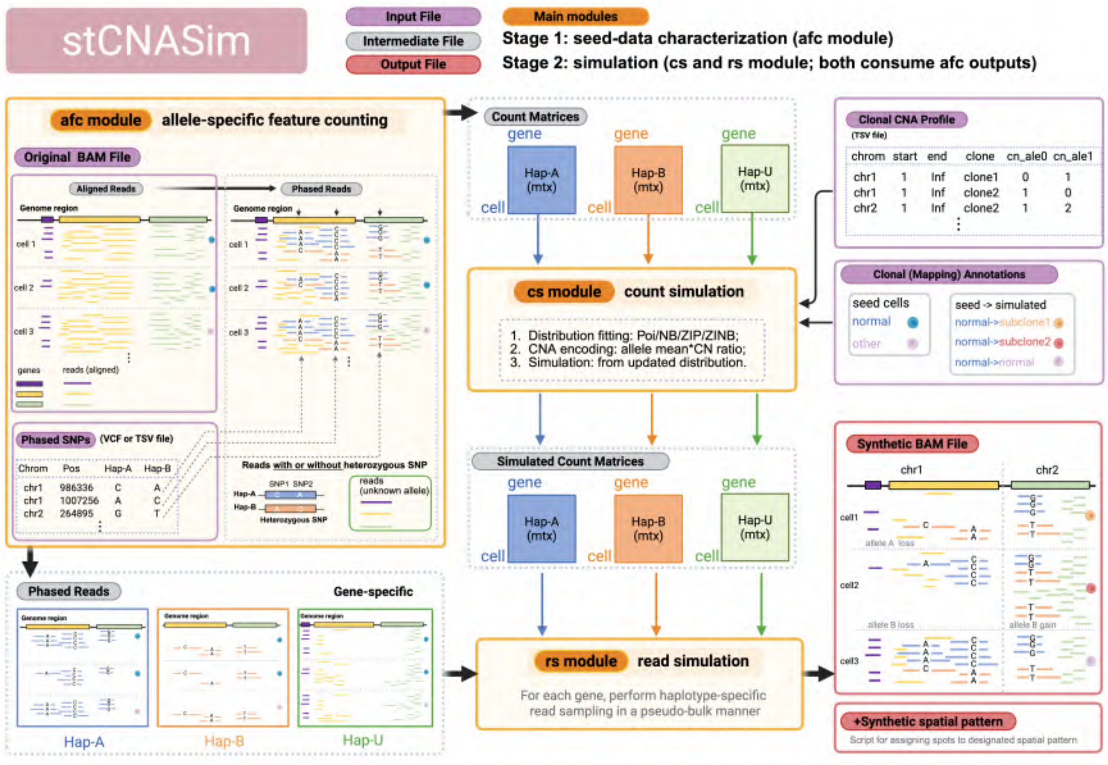


Figure S1: Overview of stCNASim, an allele-aware spatial RNA-seq simulator designed for comprehensive benchmarking of copy number alterations analysis.

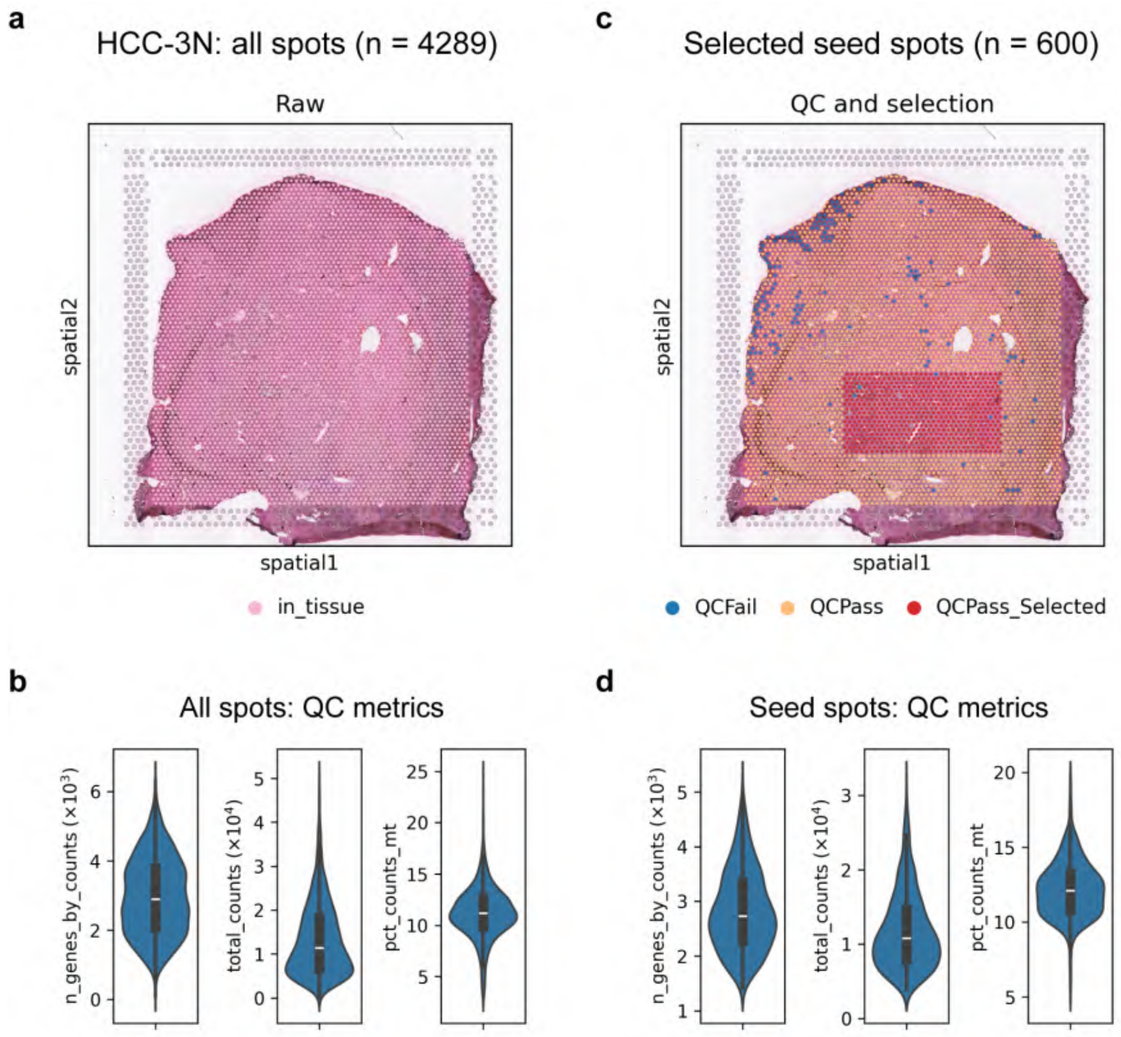


Figure S2: Seed data generation from the HCC-3N dataset. (a-b) Spatial distribution of HCC-3N in-tissue spots (n = 4289) on the H&E image and their corresponding quality control (QC) metrics. (c-d) Spatial distribution of selected seed spots (n = 600) on the H&E image and their associated QC metrics. QC metrics were calculated using `scanpy.pp.calculate_qc_metrics()`. A contiguous set of spots was selected after filtering out poor-quality data based on the following criteria: `n_genes_by_counts` $<$ 1000 or $>$ 6000; `total_counts` $<$ 2000 or $>$ 35,000; or `pct_counts_mt` $>$ 20. Panel b-d are related to and adapted from Fig. 1c.

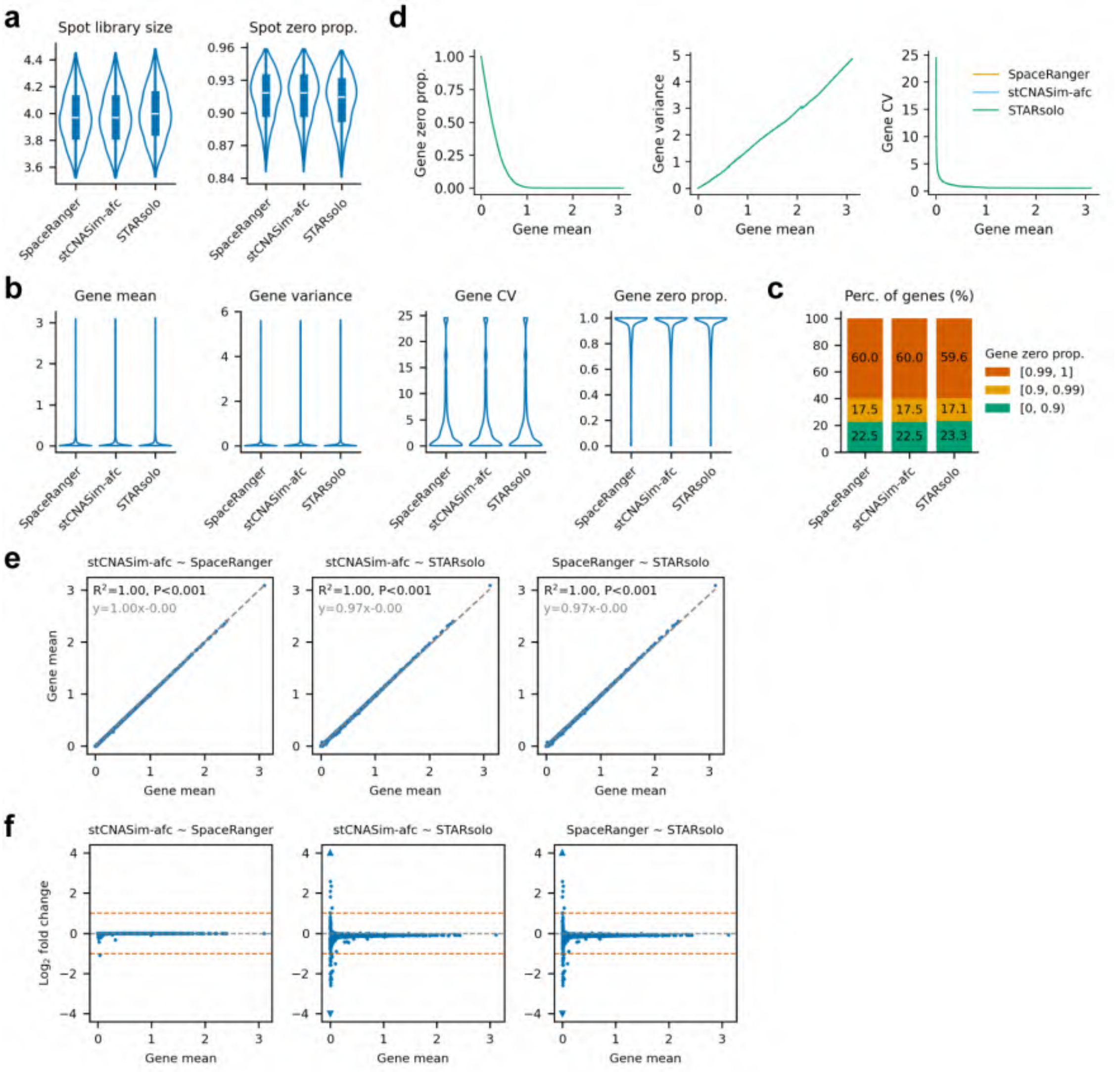


Figure S3: Internal modules: comparison of feature counting results (gene expression count matrices) on seed BAM file, generated by SpaceRanger, stCNASim-afc module, and STARsolo. The stCNASim-afc count matrix was constructed by aggregating its three haplotype-specific count matrices (i.e., $\mathbf{A} + \mathbf{B} + \mathbf{U}$). The three matrices share the same shape: 600 spots and 32272 genes. (a) Distributions of spot-wise library size and zero proportion. (b) Distributions of gene-wise mean, variance, coefficient of variation (CV), and zero proportion. (c) Distribution of genes stratified by distinct zero proportions. 'Perc.' denotes percentage. (d) Smoothed pairwise relationships between three gene-wise statistics vs. gene-wise mean across all genes. Curves were generated via k-nearest neighbor kernel regression with Gaussian distance weighting on the x-axis variable (default k=100). When plotting gene-wise CV against mean, genes with zero mean were excluded. (e) Linear regression analysis of gene expression means across different tools. Reported metrics include coefficient of determination ($R^2$), p-value (P), and regression equation. (f) Scatter plots illustrating log2 fold changes in gene mean expression between each pairwise groups, over the gene mean of the baseline (latter) group. Small triangular markers at the plot margins represent out-of-window data points. Two vermillion dashed lines denote log2 fold change values of 1 and -1, respectively. Note that gene-wise mean, gene-wise variance and spot-wise library size are transformed to the log10(1+x) scale, where x represents the original statistic value. 'prop.' denotes proportion.

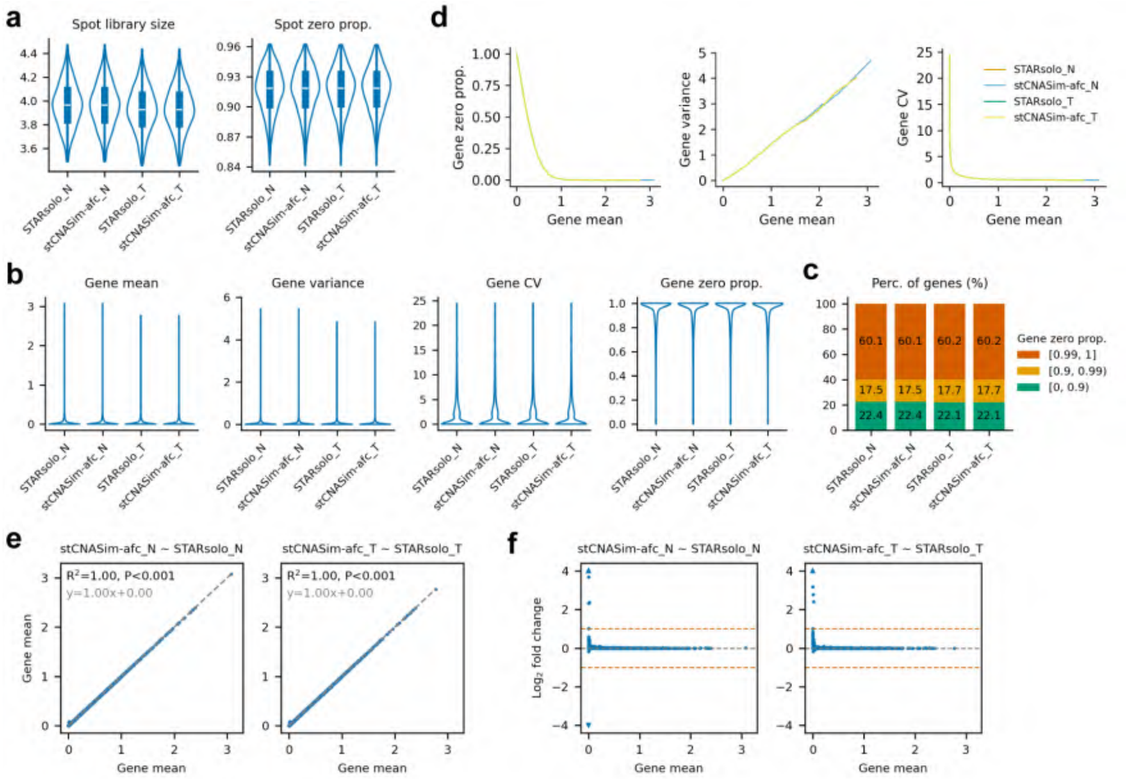


Figure S4: Internal modules: comparison of feature counting results (gene expression count matrices) on simulated BAM file, generated by the stCNASim-afc module and STARsolo. The stCNASim-afc count matrix was constructed by aggregating its three haplotype-specific count matrices (i.e., $\mathbf{A} + \mathbf{B} + \mathbf{U}$). Suffix 'N' and 'T' denote normal and tumor spots, respectively. The four matrices share the same shape: 600 spots and 32272 genes. (a) Distributions of spot-wise library size and zero proportion. (b) Distributions of gene-wise mean, variance, coefficient of variation (CV), and zero proportion. (c) Distribution of genes stratified by distinct zero proportions. 'Perc.' denotes percentage. (d) Smoothed pairwise relationships between three gene-wise statistics vs. gene-wise mean across all genes. Curves were generated via k-nearest neighbor kernel regression with Gaussian distance weighting on the x-axis variable (default k=100). When plotting gene-wise CV against mean, genes with zero mean were excluded. (e) Linear regression analysis of gene expression means across different tools. Reported metrics include coefficient of determination ($R^2$), p-value (P), and regression equation. (f) Scatter plots illustrating log2 fold changes in gene mean expression between each pairwise groups, over the gene mean of the baseline (latter) group. Small triangular markers at the plot margins represent out-of-window data points. Two vermillion dashed lines denote log2 fold change values of 1 and -1, respectively. Note that gene-wise mean, gene-wise variance and spot-wise library size are transformed to the log10(1+x) scale, where x represents the original statistic value. 'prop.' denotes proportion.

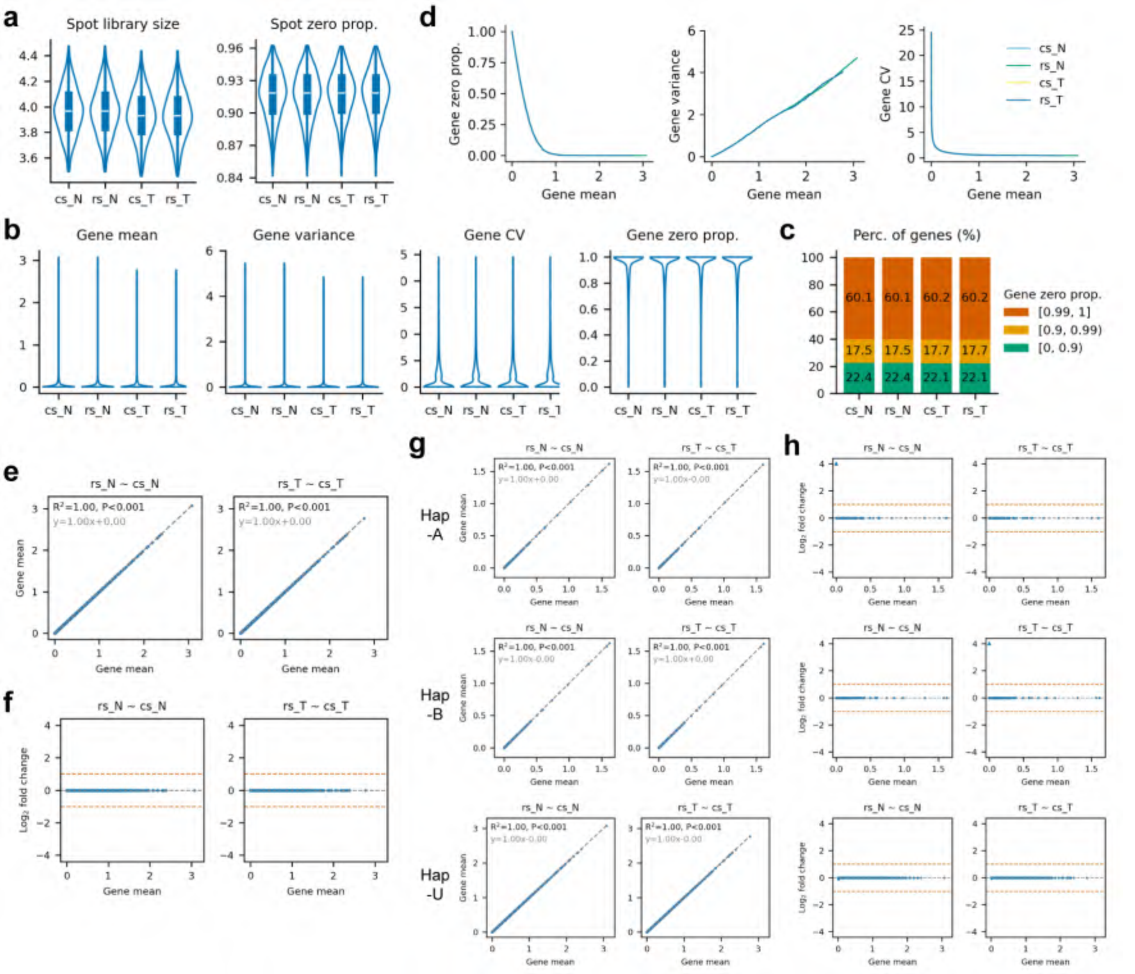


Figure S5: Internal modules: comparison of gene expression and haplotype-specific count matrices for simulated normal and tumor spots, generated by the stCNASim *cs* and *rs* modules. Suffix 'N' and 'T' denote normal and tumor spots, respectively. The four matrices share the same shape: 600 spots and 32295 genes. (a) Distributions of spot-wise library size and zero proportion. (b) Distributions of gene-wise mean, variance, coefficient of variation (CV), and zero proportion. (c) Distribution of genes stratified by distinct zero proportions. 'Perc.' denotes percentage. (d) Smoothed pairwise relationships between three gene-wise statistics vs. gene-wise mean across all genes. Curves were generated via k-nearest neighbor kernel regression with Gaussian distance weighting on the x-axis variable (default k=100). When plotting gene-wise CV against mean, genes with zero mean were excluded. (e-f) Analysis on the aggregated **X** count matrices. (e) Linear regression analysis of gene expression means across different tools. Reported metrics include coefficient of determination ($R^2$), p-value (P), and regression equation. (f) Scatter plots illustrating log2 fold changes in gene mean expression between each pairwise groups, over the gene mean of the baseline (latter) group. Small triangular markers at the plot margins represent out-of-window data points. Two vermillion dashed lines denote log2 fold change values of 1 and -1, respectively. (g-h) Similar analysis as (e-f), on three haplotype-specific count matrices **A**, **B**, and **U**. Note that gene-wise mean, gene-wise variance and spot-wise library size are transformed to the log10(1+x) scale, where x represents the original statistic value. 'prop.' denotes proportion.

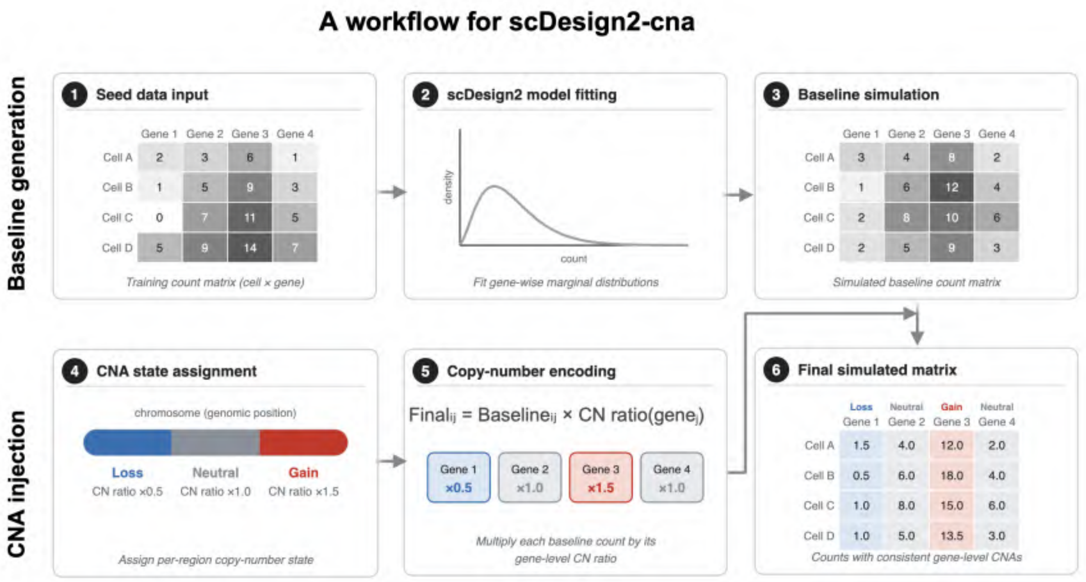


Figure S6: Benchmarking of count simulation: a workflow for scDesign2-cna. The scDesign2 simulation pipeline was adapted for CNA count generation: baseline counts were produced via conventional scDesign2 simulation, and every gene's baseline values were multiplied by a gene-specific copy number (CN) ratio corresponding to its copy number state.

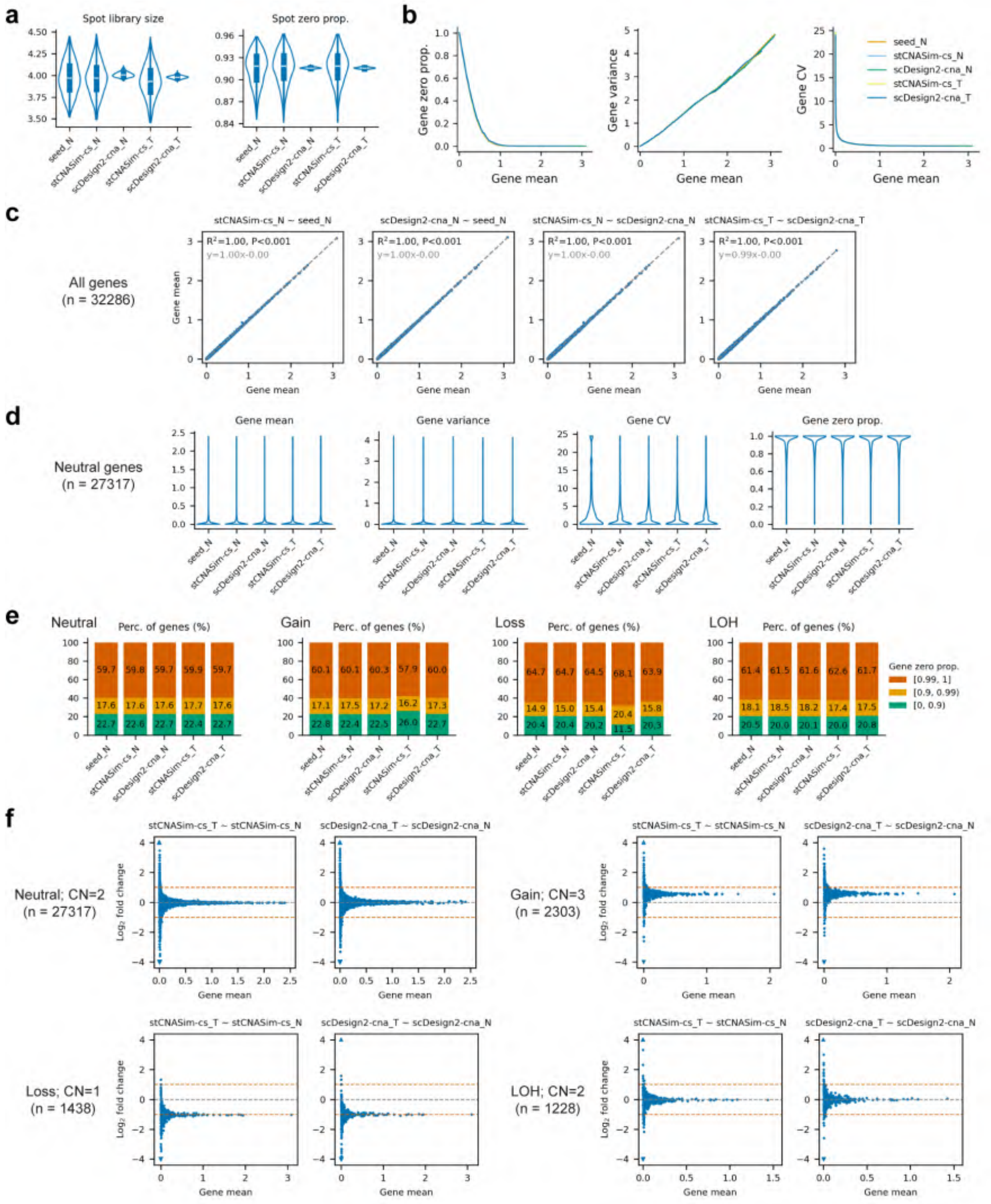


Figure S7: Benchmarking of the stCNASim count simulation module (continued on next page).

Figure S7: Benchmarking of the stCNASim count simulation module (stCNASim-cs), by comparing its simulated gene expression count matrix to (1) the seed matrix (seed) generated by feature counting with stCNASim *afc* module; and (2) the matrix simulated by modified scDesign2-independent mode (scDesign2-cna). Suffix 'N' and 'T' denote normal and tumor spots, respectively. The five matrices share the same shape: 600 spots and 32286 genes. (a) Distributions of spot-wise library size and zero proportion. (b-c) Analysis using all genes (n = 32286). (b) Smoothed pairwise relationships between three gene-wise statistics vs. gene-wise mean across all genes. (c) Linear regression analysis of gene expression means across different tools. (d) Analysis restricted to genes overlapping copy-neutral regions (n = 27317). Distributions of four gene-wise metrics. (e-f) Stratified analysis using genes overlapping regions with four distinct CNA states, analyzed separately. (e) Distribution of genes stratified by distinct zero proportions. 'Perc.' denotes percentage. (f) Scatter plots illustrating log2 fold changes in gene mean expression between each pairwise groups, over the gene mean of the baseline (latter) group. CN: copy number. Note that gene-wise mean, gene-wise variance and spot-wise library size are transformed to the log10(1+x) scale, where x represents the original statistic value. 'prop.' denotes proportion.

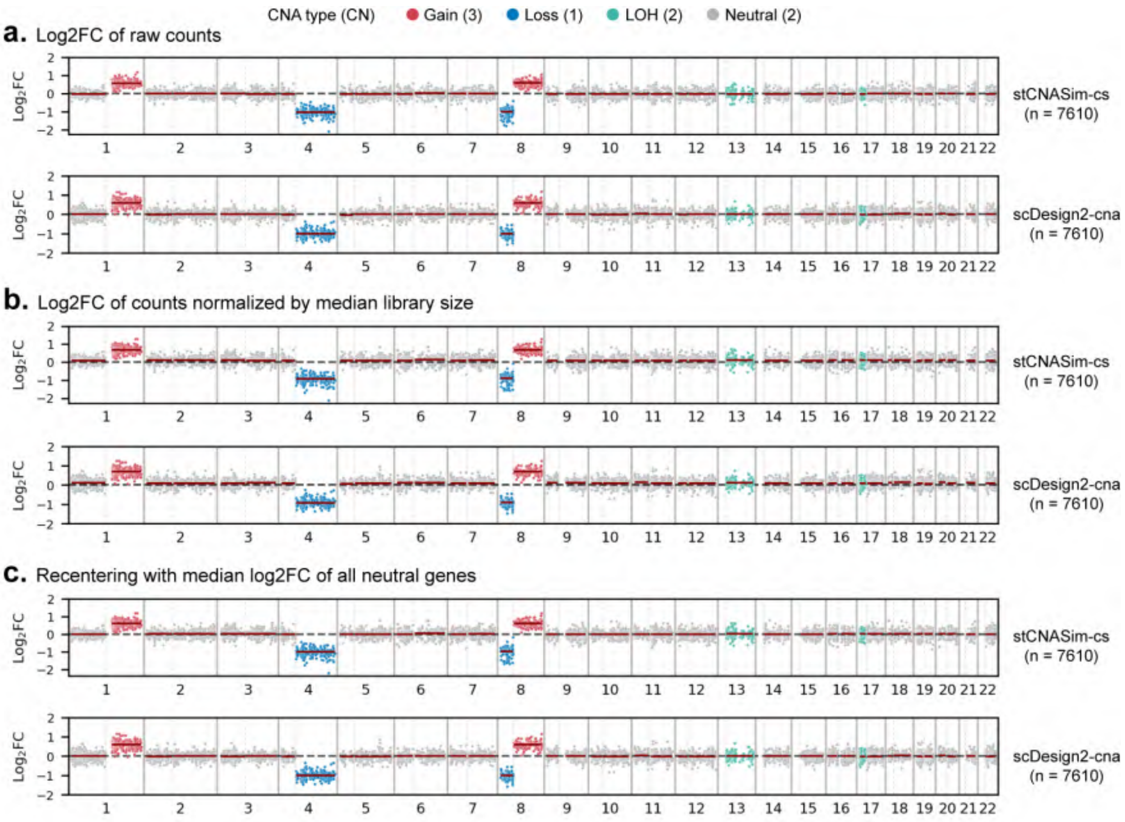


Figure S8: Benchmarking of count simulation: pseudobulk log2 fold-change (log2FC) profiles between tumor and normal spots derived from stCNASim-cs and scDesign2-cna simulations. Genes were pre-filtered by excluding those with fewer than three expressing cells or a mean expression level below 0.1 in the seed data. (a) Log2FC values computed from raw expression counts. (b) Log2FC values based on counts normalized by median library size; median library sizes were independently estimated using all simulated spots for each method, i.e., 600 normal spots for seed data and 1200 normal and tumor spots for stCNASim-cs and scDesign2-cna simulated data. (c) Re-centering of panel (b) log2FC values via subtraction of the median log2FC of all neutral genes. CN: copy number.

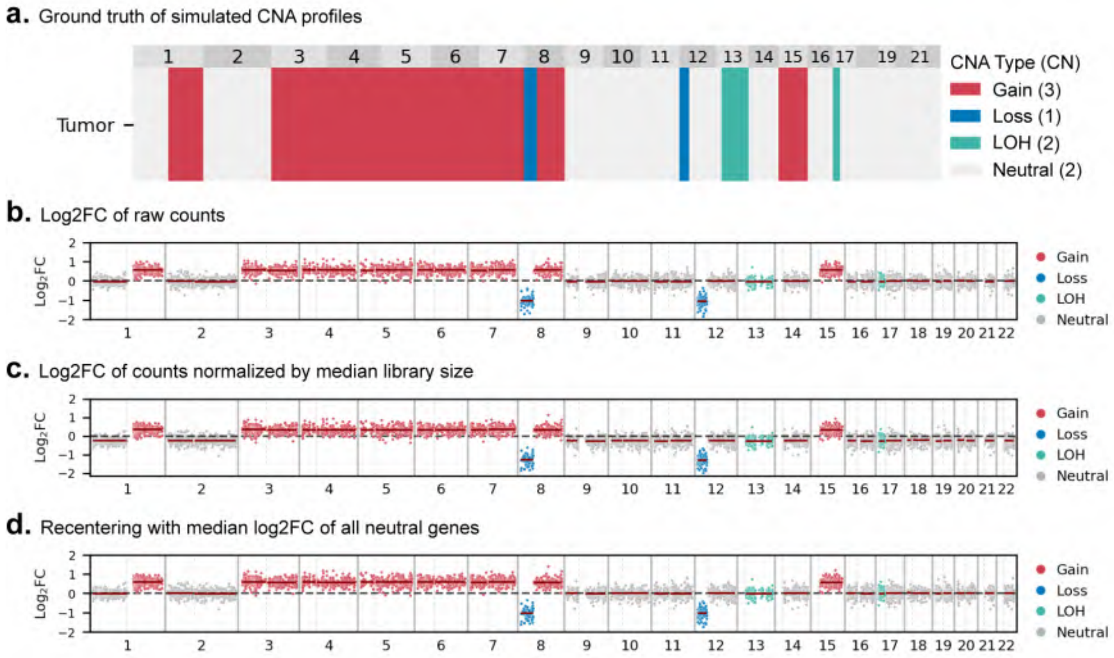


Figure S9: Benchmarking of count simulation: pseudobulk log2 fold-change (log2FC) profiles of a stCNASim simulated dataset with strong compositional effects. (a) Ground truth of simulated CNA profiles. CN: copy number. (b-d) Log2FC profiles between tumor and normal spots derived from stCNASim-cs simulation. Genes were pre-filtered by excluding those with fewer than three expressing cells or a mean expression level below 0.1 in the seed data. (b) Log2FC values computed from raw expression counts. (c) Log2FC values based on counts normalized by median library size; median library sizes were estimated using all simulated spots, i.e., 1200 normal and tumor spots. (d) Re-centering of panel (b) log2FC values via subtraction of the median log2FC of all neutral genes.

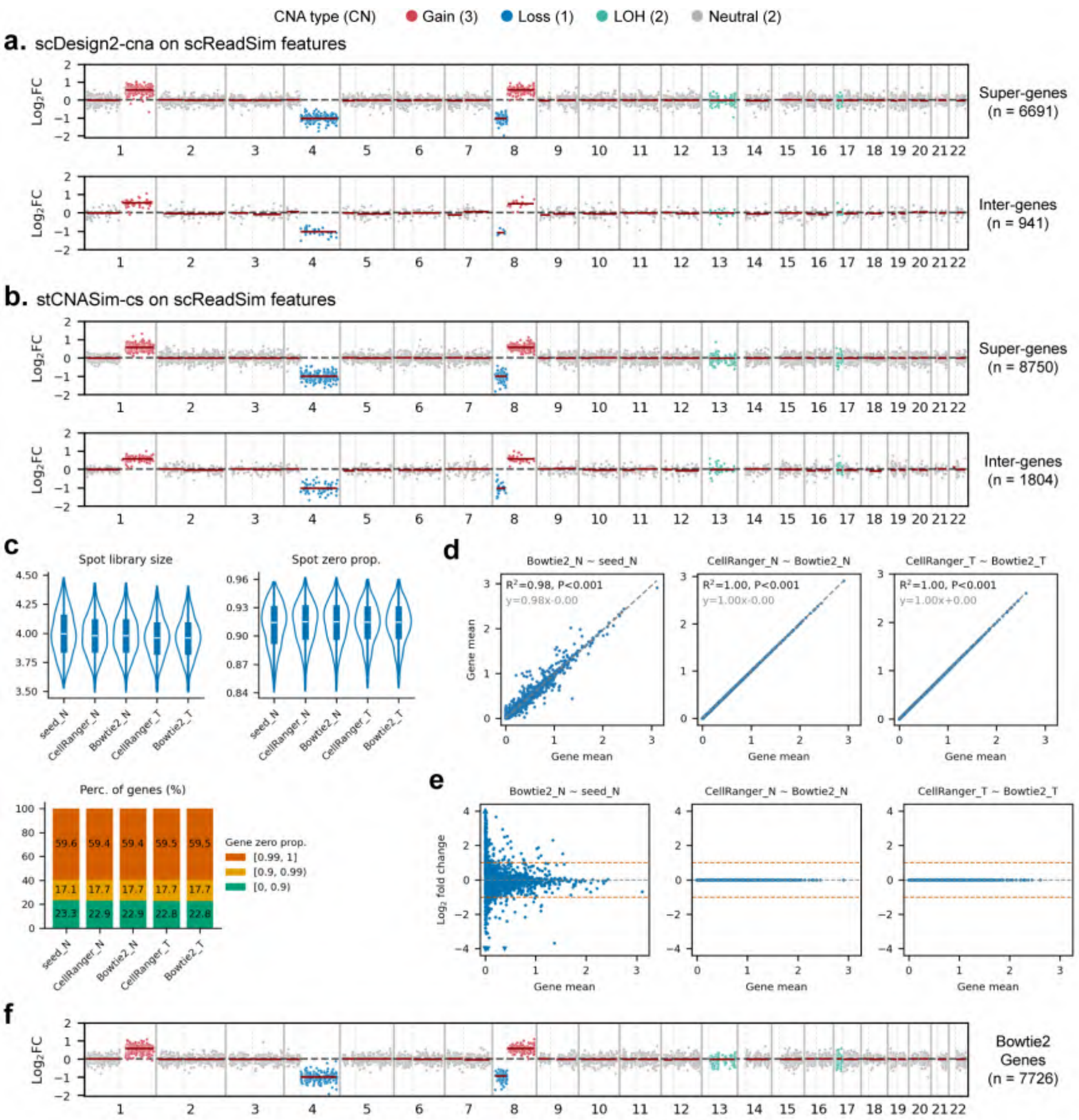


Figure S10: Benchmarking of read simulation: comparison of modified scReadSim workflow configurations for CNA read simulation. (a-b) Log2FC profiles between count matrices of simulated tumor and normal spots prior to read simulation. Count matrices were generated by running scDesign2-cna (a) and stCNASim-cs (b) on the scReadSim features (super-genes and inter-genes). For log2FC calculation, features were pre-filtered by excluding those with fewer than three expressing cells or a mean expression level below 0.1 in the simulated normal spots (a) or seed normal spots (b). Notably, the count matrix simulated by scDesign2-cna is incompatible with scReadSim due to `dtype` mismatch. (c-f) Comparison of the BAM files simulated via scReadSim downstream of stCNASim-cs. FASTQ files generated by scReadSim were separately aligned to the hg38 reference genome using Bowtie2 and CellRanger. All BAM files (including the seed BAM file) were quantified using STARsolo feature counting. The five matrices share the same shape: 600 spots and 32272 genes. Suffix 'N' and 'T' denote normal and tumor spots, respectively. (c) Spot-wise and gene-wise metrics. (d) Linear regression analysis of gene expression means across different tools. (e) Scatter plots illustrating log2 fold changes in gene mean expression between each pairwise groups, over the gene mean of the baseline (latter) group. (f) Log2FC values (tumor vs. normal) computed from raw expression counts of the Bowtie2-aligned BAM file. CN: copy number.

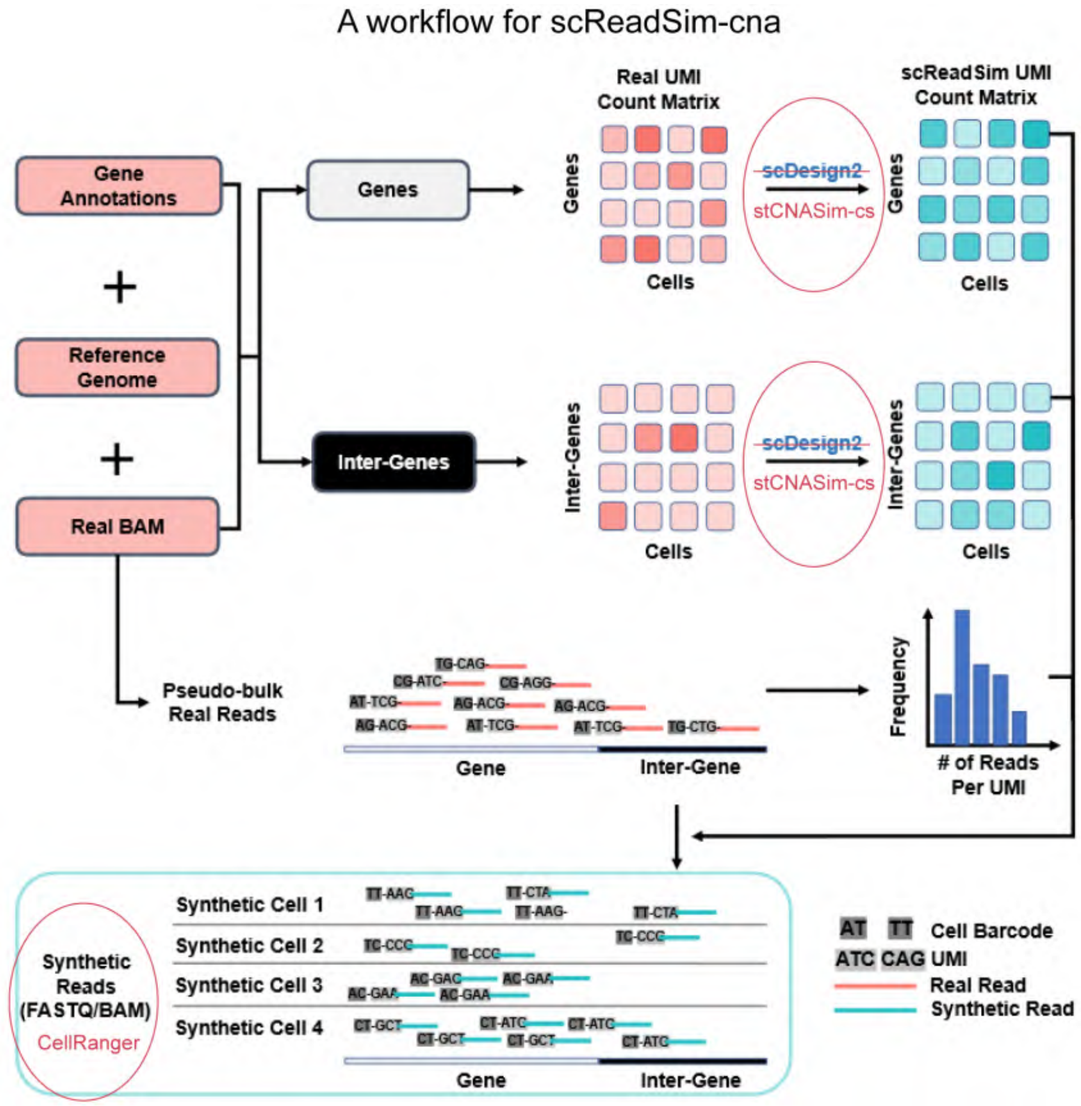


Figure S11: Benchmarking of read simulation: a workflow for scReadSim-cna. We modified the standard scReadSim simulation pipeline for CNA-aware read generation with two key adjustments (marked by red circles): (1) stCNASim-cs was adopted for CNA count simulation to replace scDesign2; and (2) CellRanger was used to align synthetic FASTQ files to BAM files, rather than Bowtie2. This modification generates the `xf` tag within the output BAM file, a mandatory input for the stCNASim allele-specific feature counting (*afc*) module. Figure adapted from Figure 1a of the original scReadSim manuscript (Yan et al., 2023 [1]).

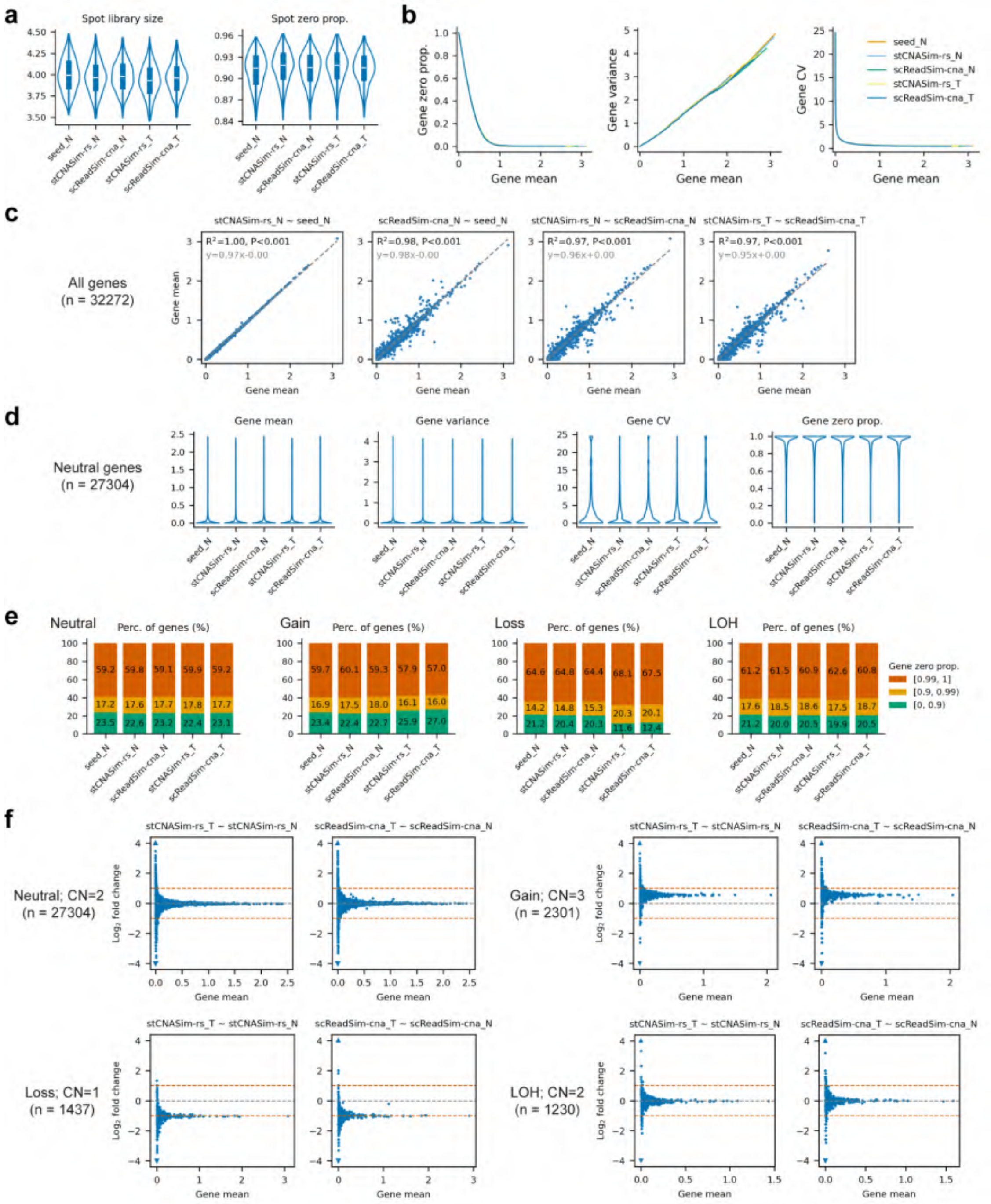


Figure S12: Benchmarking the RDR signals generated by the read simulation module of stCNASim (continued on next page).

Figure S12: Benchmarking the RDR signals generated by the read simulation module of stCNASim (stCNASim-rs). Suffix 'N' and 'T' denote normal and tumor spots, respectively. All BAM files (including the seed BAM file) were quantified using STARsolo feature counting. The five matrices share the same shape: 600 spots and 32272 genes. (a) Distributions of spot-wise library size and zero proportion. (b-c) Analysis using all genes (n = 32272). (b) Smoothed pairwise relationships between three gene-wise statistics vs. gene-wise mean across all genes. (c) Linear regression analysis of gene expression means across different tools. (d) Analysis restricted to genes overlapping copy-neutral regions (n = 27304). Distributions of four gene-wise metrics. (e-f) Stratified analysis using genes overlapping regions with four distinct CNA states, analyzed separately. (e) Distribution of genes stratified by distinct zero proportions. *Perc.* denotes percentage. (f) Scatter plots illustrating log2 fold changes in gene mean expression between each pairwise groups, over the gene mean of the baseline (latter) group. CN: copy number. Note that gene-wise mean, gene-wise variance and spot-wise library size are transformed to the log10(1+x) scale, where x represents the original statistic value. 'prop.' denotes proportion.

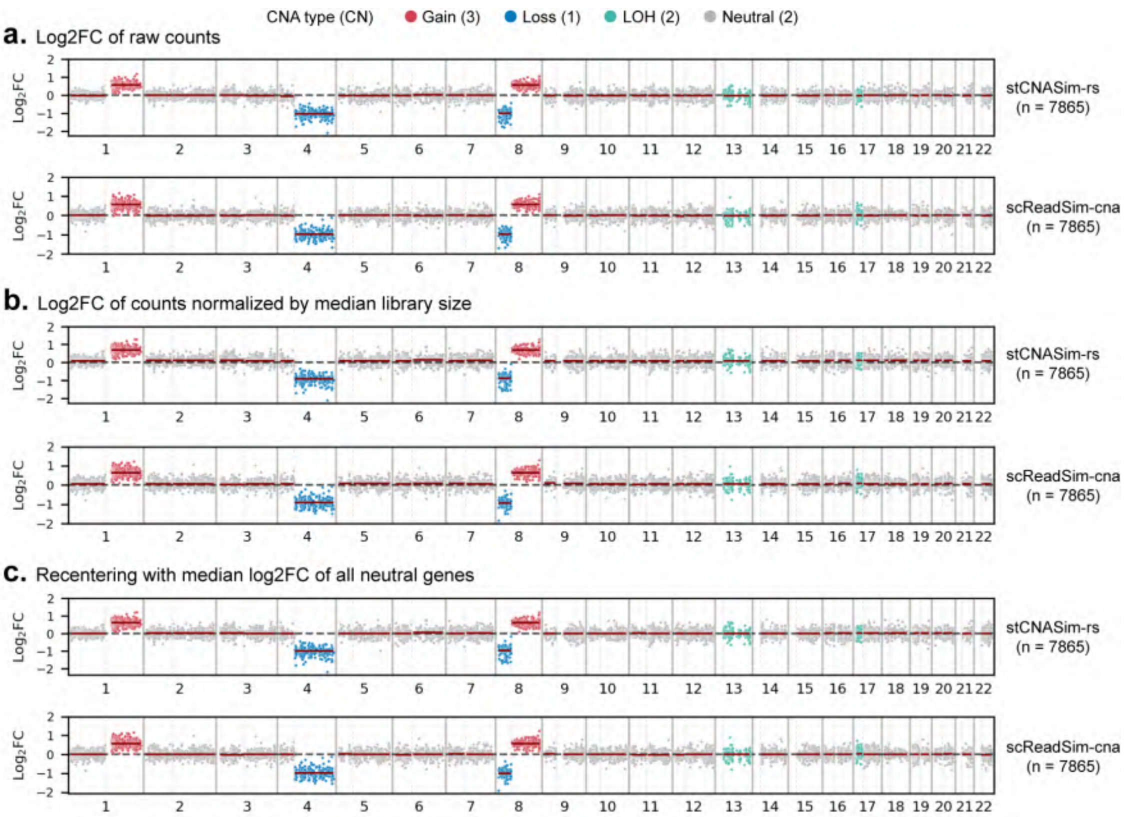


Figure S13: Benchmarking the RDR signals generated by the read simulation module of stCNASim (stCNASim-rs): pseudobulk log2 fold-change (log2FC) profiles between tumor and normal spots derived from stCNASim-rs and scReadSim-cna simulations. Genes were pre-filtered by excluding those with fewer than three expressing cells or a mean expression level below 0.1 in the seed data. (a) Log2FC values computed from raw expression counts. (b) Log2FC values based on counts normalized by median library size; median library sizes were independently estimated using all simulated spots for each method, i.e., 600 normal spots for seed data and 1200 normal and tumor spots for stCNASim-rs and scReadSim-cna simulated data. (c) Re-centering of panel (b) log2FC values via subtraction of the median log2FC of all neutral genes. CN: copy number. Panel a is related to and adapted from Fig. 1e.

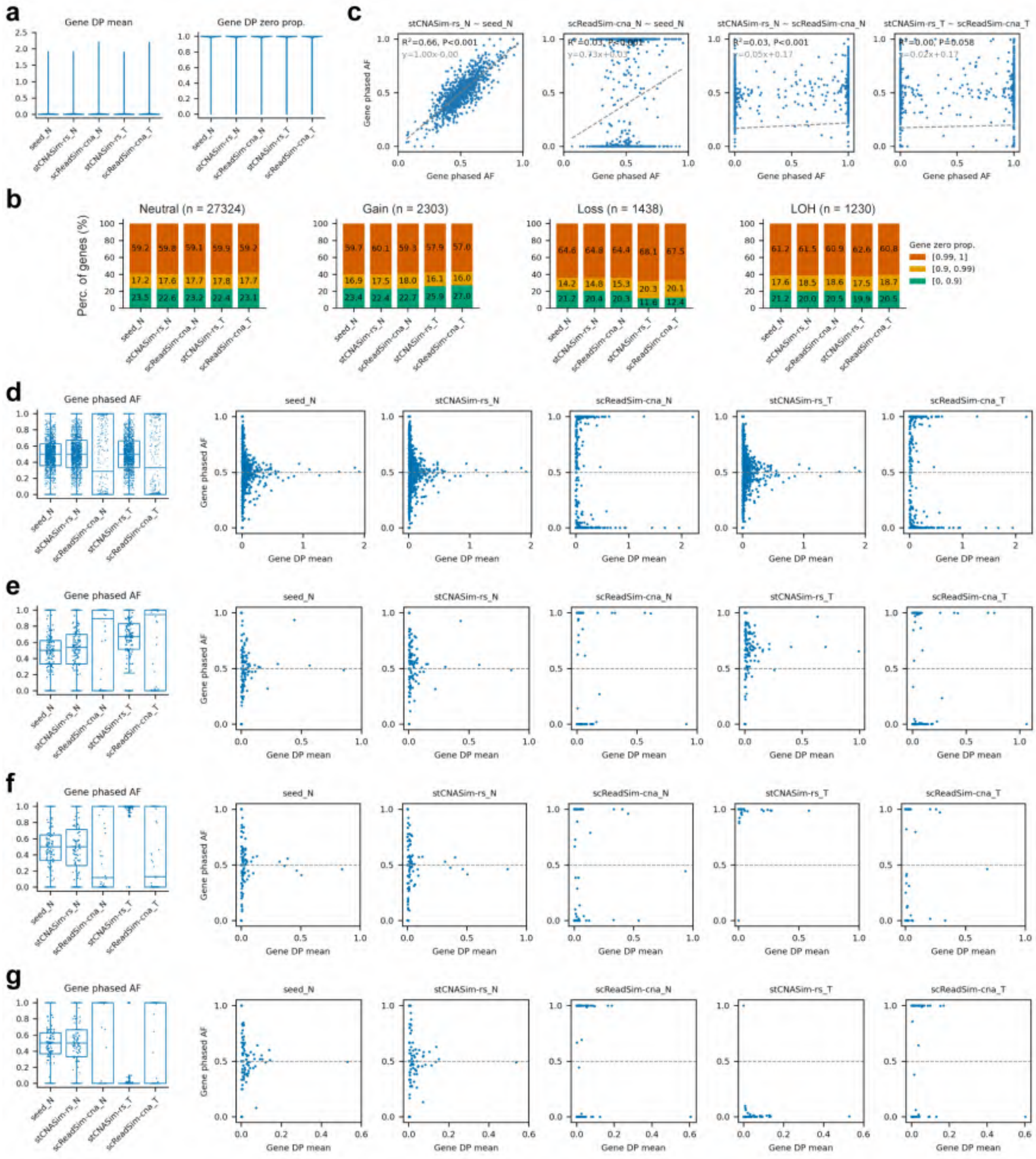

Figure S14: Benchmarking the gene-level BAF signals generated by the read simulation module of stCNASim (stCNASim-rs) (continued on next page).

Figure S14: Benchmarking the gene-level BAF signals generated by the read simulation module of stCNASim (stCNASim-rs). Suffix 'N' and 'T' denote normal and tumor spots, respectively. All BAM files (including the seed BAM file) were quantified via the stCNASim *afc* allelic feature counting. The five matrices share the same shape: 600 spots and 32295 genes. (a) Distributions of two gene-wise metrics, evaluated across all 32295 genes. (b) Distribution of genes stratified by distinct zero proportions. 'Perc.' denotes percentage. (c) Linear regression analysis of gene phased AF across different tools. In each group, genes whose aggregated DP $< 10$ or AF is `NaN` were excluded prior to regression. (d-g) Stratified analysis using genes overlapping regions with four distinct CNA states, analyzed separately: copy neutral CN=2 (d), copy gain CN=3 (e), copy loss CN=1 (f), and LOH CN=2 (g). For each panel set (d-g), the left sub-panel shows the distribution of gene-wise phased AF, whereas the right sub-panel presents the scatter plots illustrating the gene phased AF over the gene DP mean. CN: copy number. Note that gene-wise DP mean is transformed to the log10(1+x) scale, where x represents the original statistic value. AF: allele frequency. DP: depth of both alleles (i.e., $\mathbf{A}+\mathbf{B}$). 'prop.' denotes proportion.

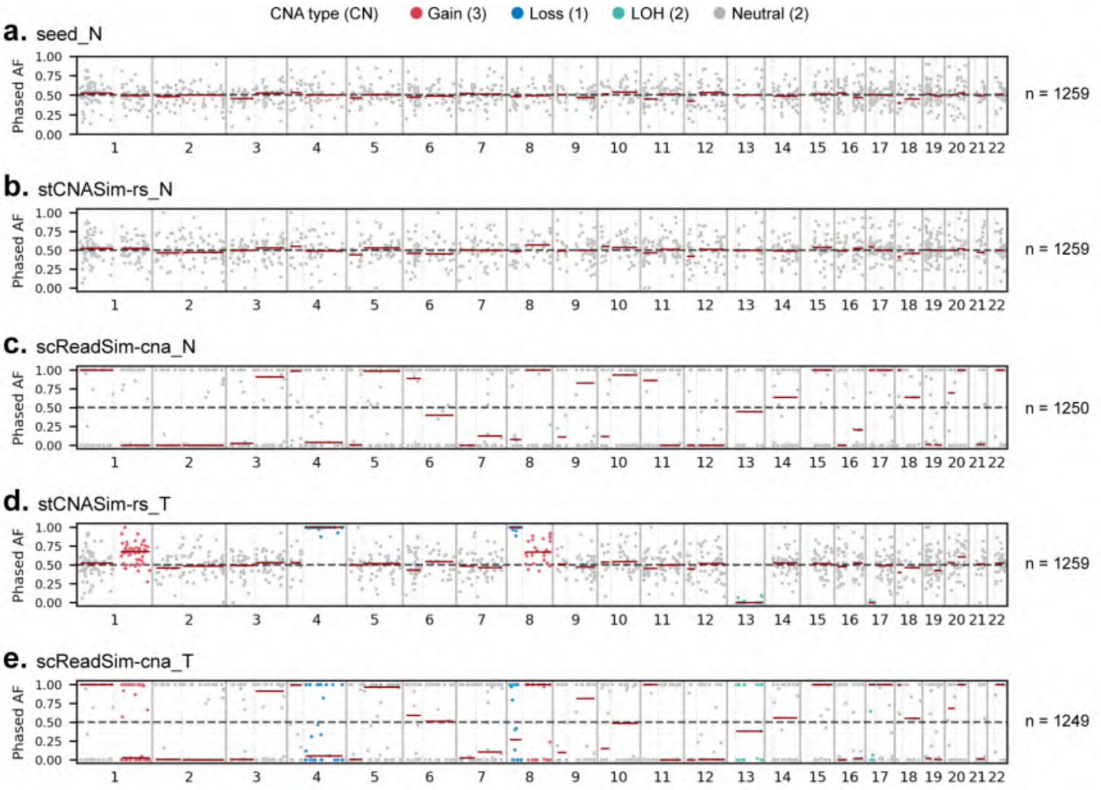


Figure S15: Benchmarking the gene-level BAF signals generated by the read simulation module of stCNASim (stCNASim-rs): pseudobulk phased allele frequency (AF) profiles derived from stCNASim-rs and scReadSim-cna simulations. Genes were pre-filtered by excluding those with aggregated DP $< 10$, or AF falling outside the 0.1-0.9 range in the seed data. Genes yielding non-numeric (`NaN`) BAF values were further discarded per analytical group. CN: copy number.

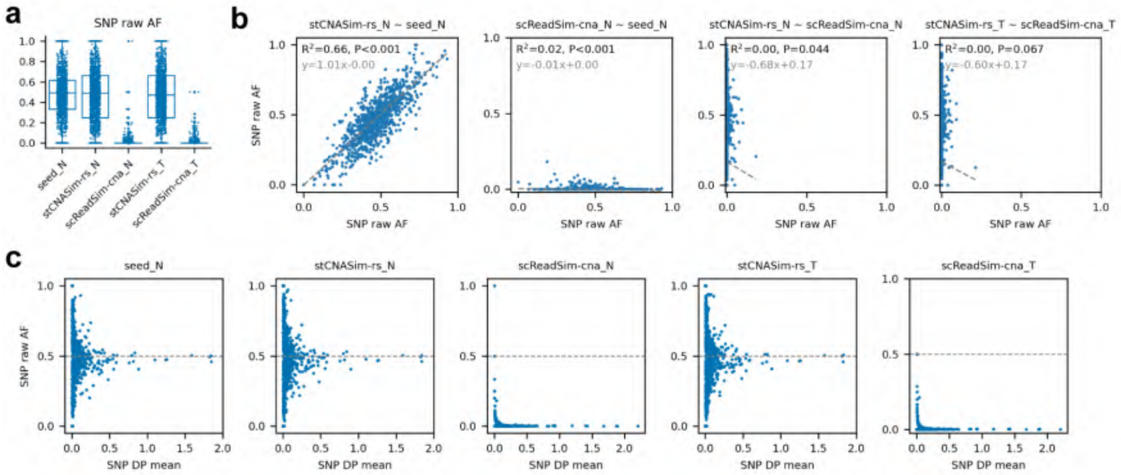


Figure S16: Statistics of the SNP-level raw BAF signals generated by the read simulation module of stCNASim (stCNASim-rs). Suffix 'N' and 'T' denote normal and tumor spots, respectively. All BAM files (including the seed BAM file) were pileup via the cellsnp-lite mode 1a, with the input 8866 phased SNPs. Analyses were restricted to a subset of 7381 SNPs residing within copy-neutral genomic regions. The five resulting matrices share the same shape: 600 spots and 7381 SNPs. (a) Distribution of SNP-wise raw AF. (b) Linear regression analysis of SNP raw AF across different tools. In each group, SNPs whose aggregated DP $< 10$ or AF is `NaN` were excluded prior to regression. (c) Scatter plots illustrating the SNP raw AF over the SNP DP mean. Note that SNP-wise DP mean is transformed to the log10(1+x) scale, where x represents the original statistic value. AF: allele frequency. DP: depth of both alleles (i.e., $\mathbf{A} + \mathbf{B}$).

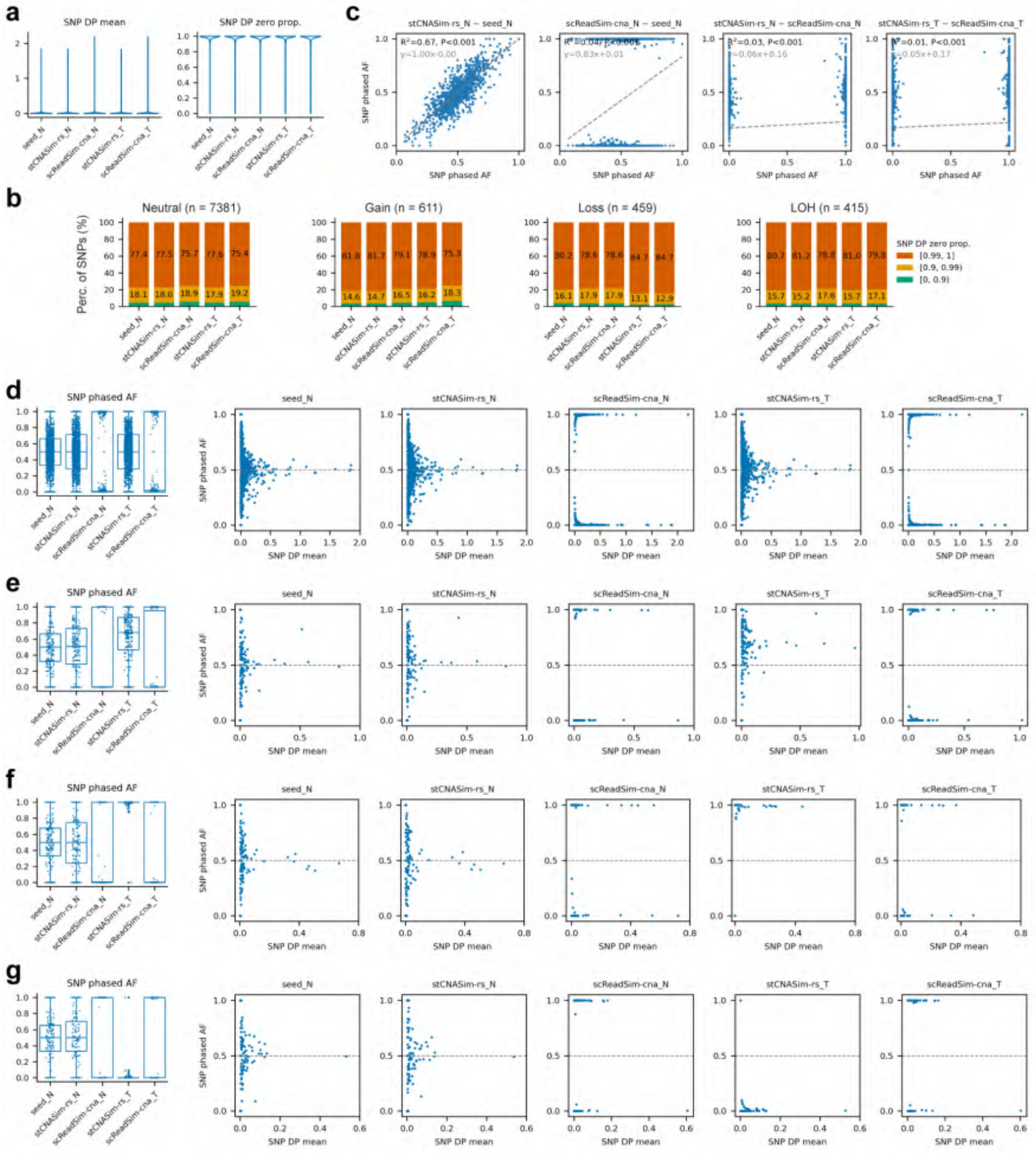

Figure S17: Benchmarking the SNP-level BAF signals generated by the read simulation module of stCNASim (stCNASim-rs) (continued on next page).

Figure S17: Benchmarking the SNP-level BAF signals generated by the read simulation module of stCNASim (stCNASim-rs). Suffix 'N' and 'T' denote normal and tumor spots, respectively. All BAM files (including the seed BAM file) were pileup via the cellsnp-lite mode 1a, with the input 8866 phased SNPs. SNP-wise phased AF was obtained by flipping raw AF values according to each SNP's phasing information. The five resulting matrices share the same shape: 600 spots and 8866 SNPs. (a) Distributions of two SNP-wise metrics, evaluated across all 8866 SNPs. (b) Distribution of SNPs stratified by distinct zero proportions. 'Perc.' denotes percentage. (c) Linear regression analysis of SNP phased AF across different tools. In each group, SNPs whose aggregated DP $<$ 10 or AF is `NaN` were excluded prior to regression. (d-g) Stratified analysis using SNPs located within regions with four distinct CNA states, analyzed separately: copy neutral CN=2 (d), copy gain CN=3 (e), copy loss CN=1 (f), and LOH CN=2 (g). For each panel set (d-g), the left sub-panel shows the distribution of SNP-wise phased AF, whereas the right sub-panel presents the scatter plots illustrating the SNP phased AF over the SNP DP mean. CN: copy number. Note that SNP-wise DP mean is transformed to the log10(1+x) scale, where x represents the original statistic value. AF: allele frequency. DP: depth of both alleles (i.e., $\mathbf{A} + \mathbf{B}$). 'prop.' denotes proportion.

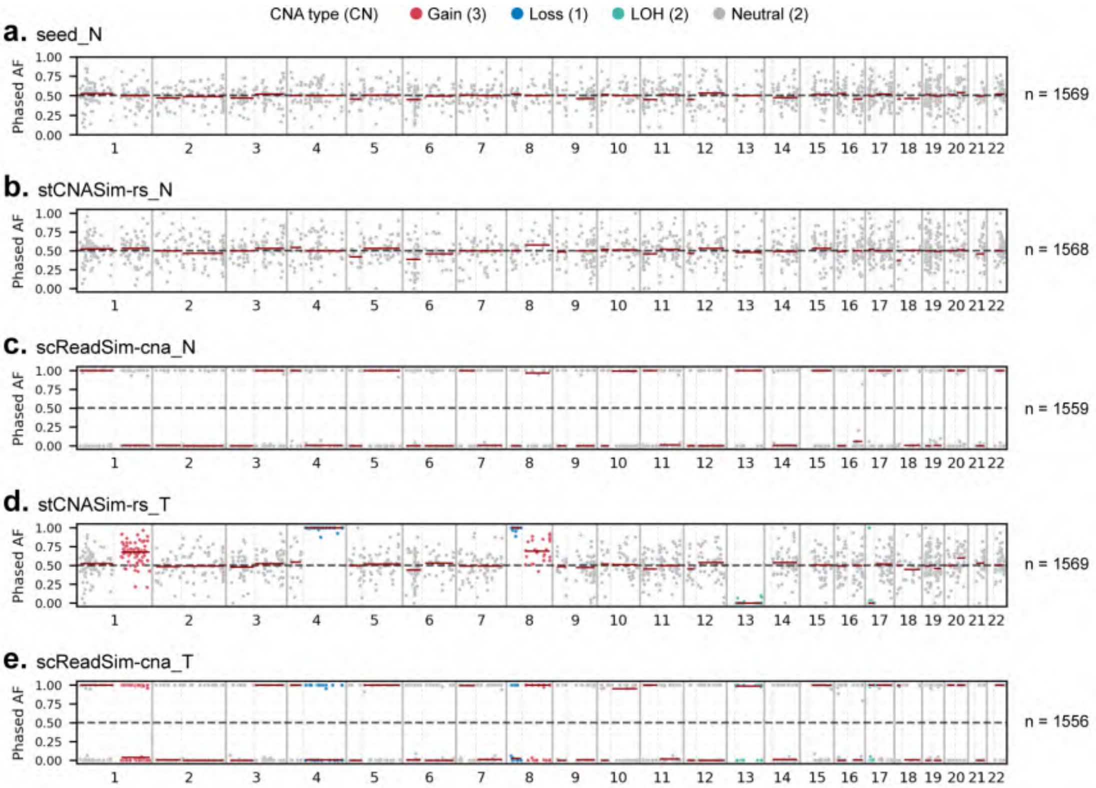


Figure S18: Benchmarking the SNP-level BAF signals generated by the read simulation module of stCNASim (stCNASim-rs): pseudobulk phased allele frequency (AF) profiles derived from stCNASim-rs and scReadSim-cna simulations. SNPs were pre-filtered by excluding those with aggregated DP $<$ 10, or AF falling outside the 0.1-0.9 range in the seed data. SNPs yielding non-numeric (`NaN`) BAF values were further discarded per analytical group. CN: copy number. Panel d-e are related to and adapted from Fig. 1f.

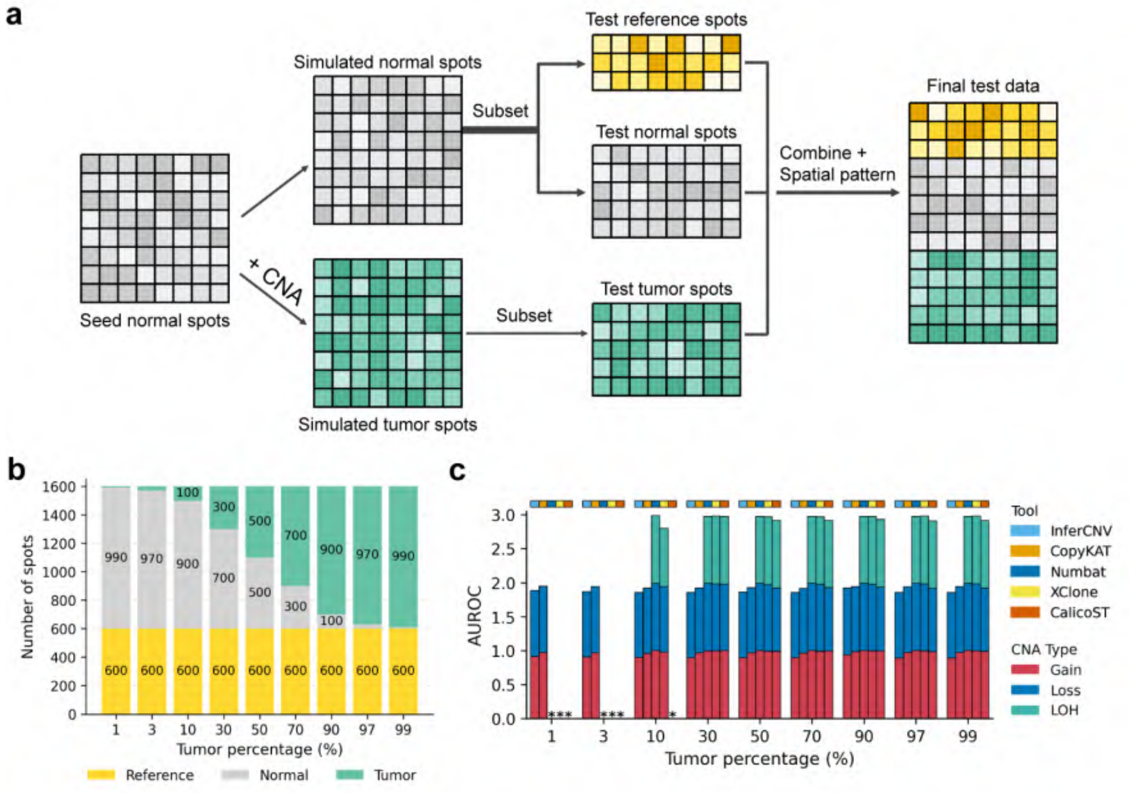


Figure S19: Overview of CNA prediction experiments across varying tumor percentages (simulated data). (a) Illustration of the data generation process. (b) Distribution of reference, normal, and tumor spots across tumor percentages. (c) AUROC of CNA profiles predicted by the five tools at each percentage. An asterisk (*) indicates no CNA was detected or a technical error occurred.

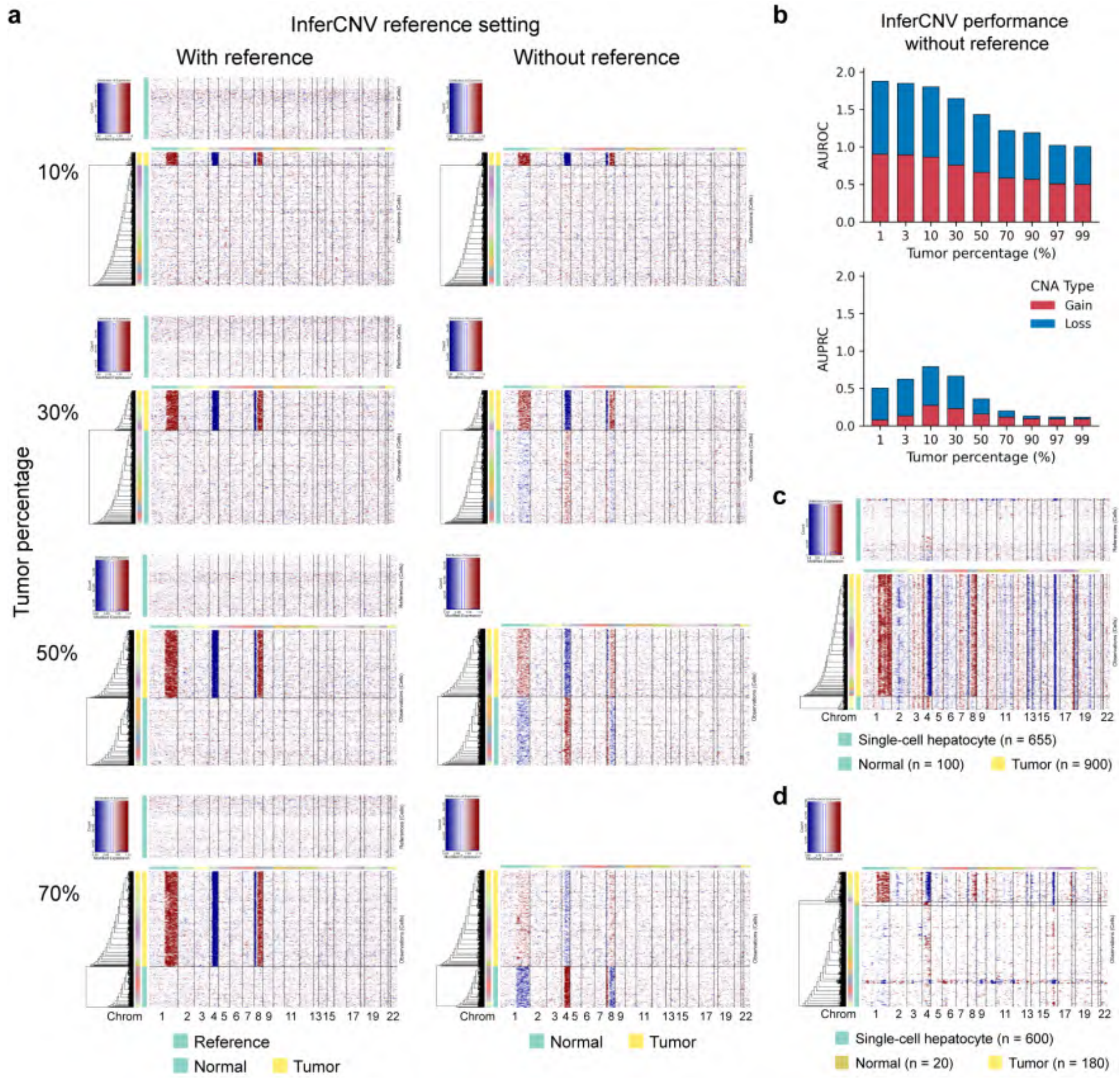


Figure S20: InferCNV performance in CNA prediction with and without reference spots (simulated data). (a) InferCNV predicted CNA profiles across various tumor percentages and reference settings. (b) AUROC and AUPRC of CNA profiles predicted by InferCNV when no reference is provided. (c) InferCNV predicted CNA profile using external hepatocyte cells from a scRNA-seq dataset as a reference (at 90% tumor percentage). (d) InferCNV predicted CNA profile when scRNA-seq hepatocyte cells are added to the analysis but not utilized as a reference.

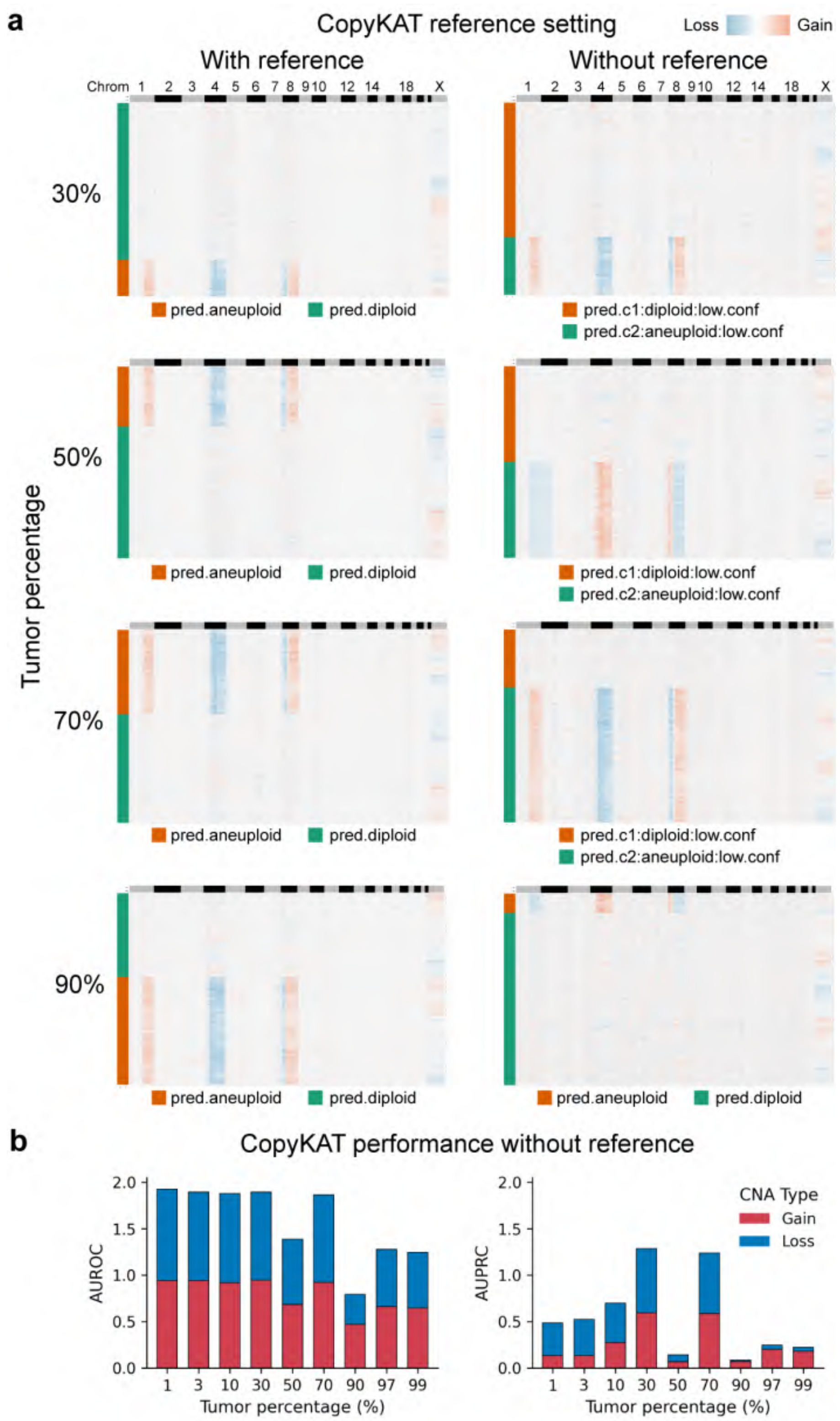


Figure S21: CopyKAT performance in CNA prediction with and without reference spots (simulated data). (a) CopyKAT predicted CNA profiles across various tumor percentages and reference settings. (b) AUROC and AUPRC of CNA profiles predicted by CopyKAT when no reference is provided.

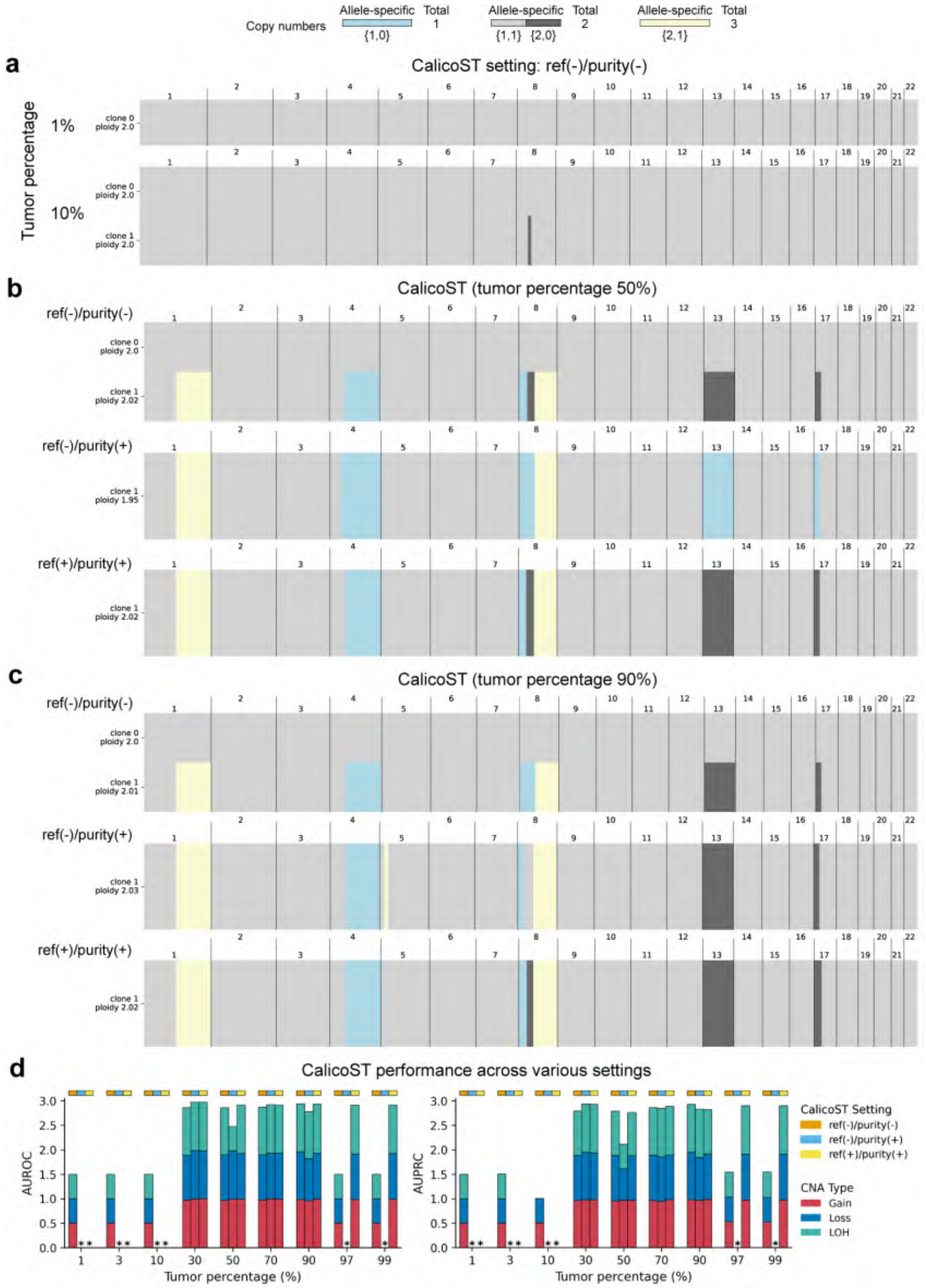


Figure S22: CalicoST performance in CNA prediction across various settings (simulated data). Specifically, ref(+/-) denotes the use of reference spots; purity(+/-) indicates whether the `purity` step was performed. At all tumor percentages, errors occur under the ref(+)/purity(-) configuration (with reference but without purity estimation). (a) CalicoST predicted CNA profiles at 1% and 10% tumor percentage [ref(-)/purity(-)]. (b-c) CalicoST predicted profiles at 50% (b) and 90% (c) tumor percentage across all settings. (d) AUROC and AUPRC metrics for each setting. An asterisk (*) indicates no CNA was detected or a technical error occurred.

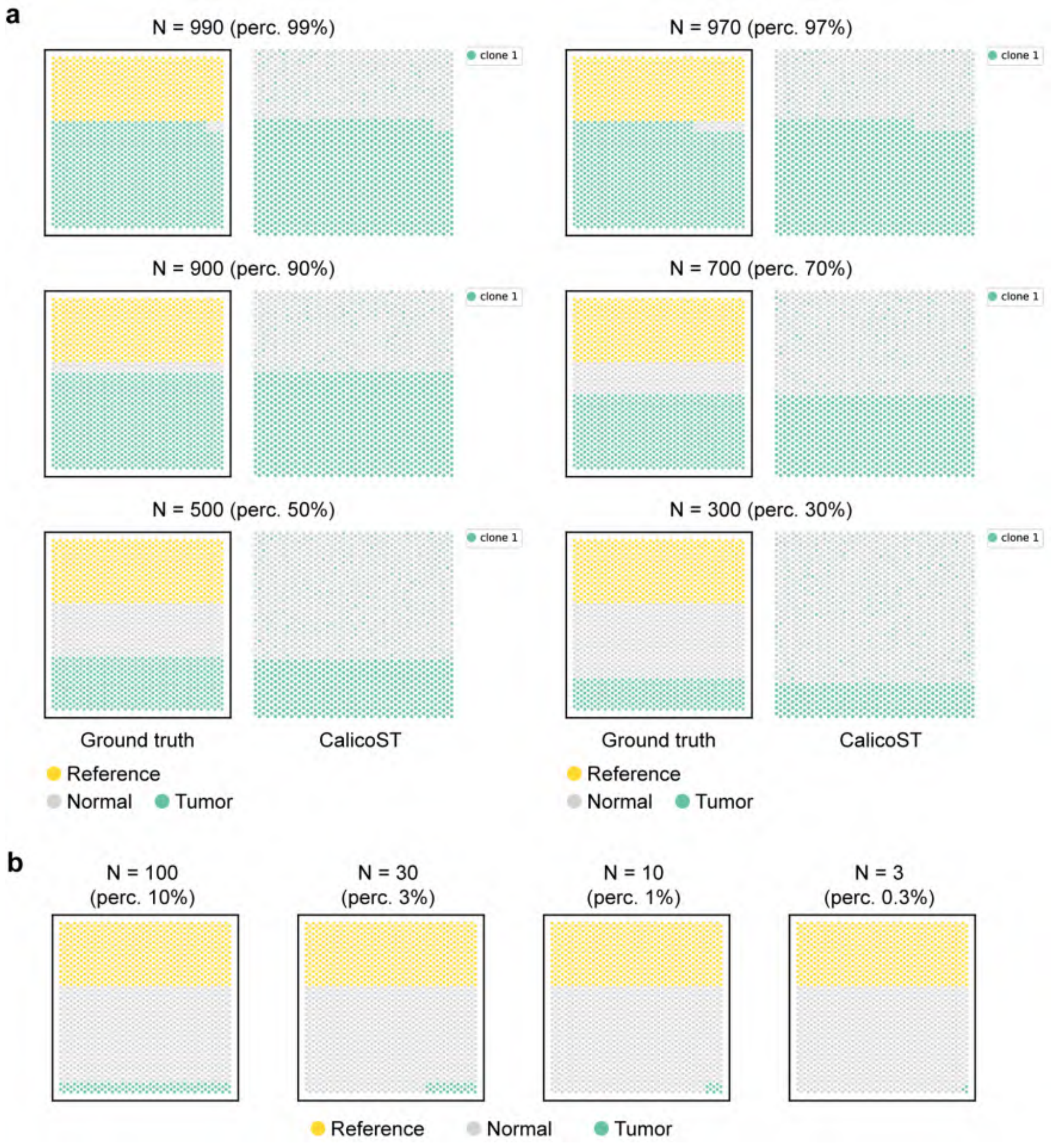


Figure S23: Spatial distribution of ground truth and CalicoST-identified tumor spots with a provided reference [ref(+)/purity(+)] (simulated data). (a) Comparison of ground truth and CalicoST-identified tumor spots across varying tumor spot densities. (b) Spatial distribution of ground truth only; CalicoST failed to generate outputs at these lower tumor spot counts. 'perc.' denotes percentage. Panel a (99% percentage) is related to and adapted from Fig. 2c.

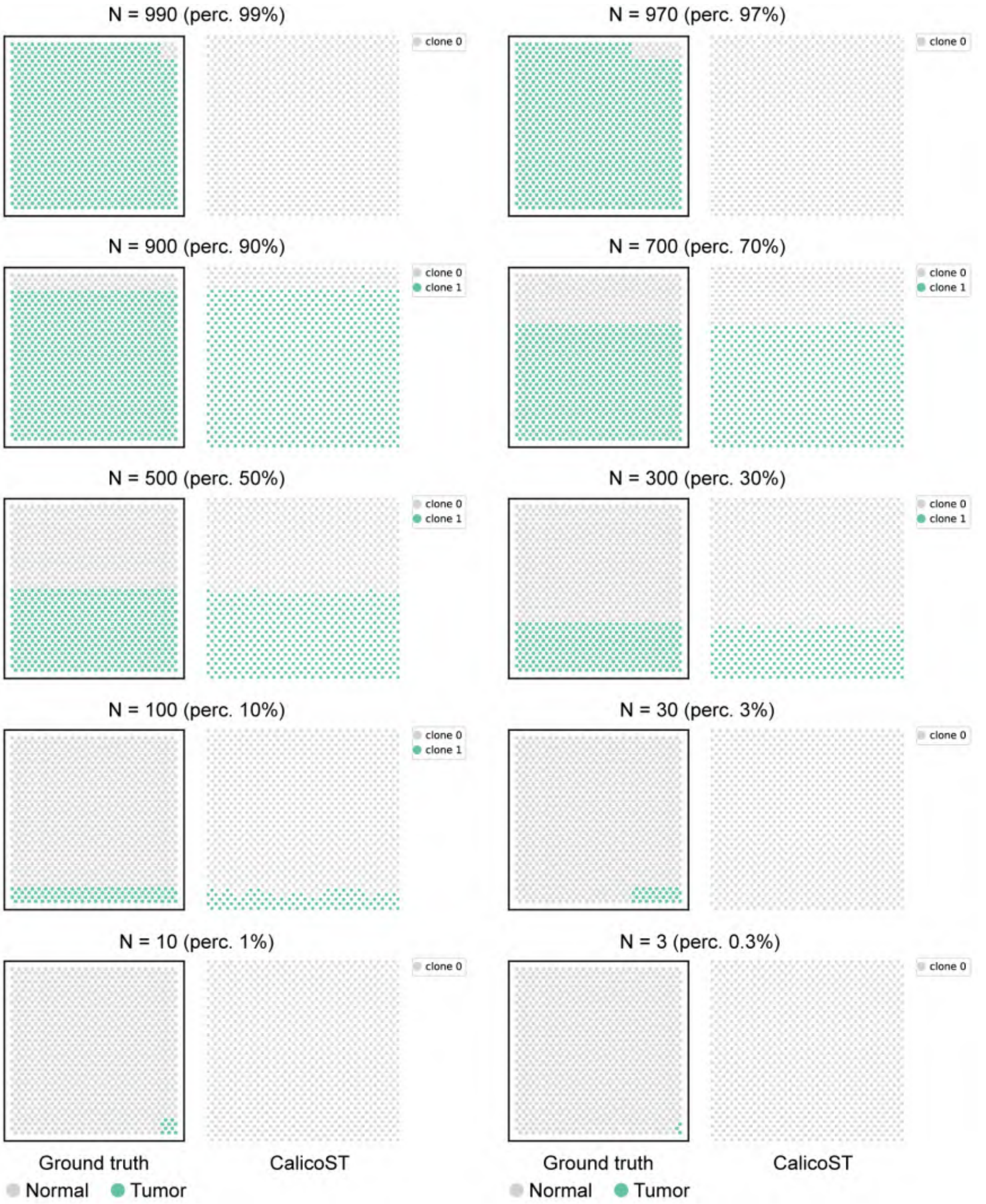


Figure S24: Spatial distribution of ground truth and CalicoST-identified tumor spots in the absence of a reference [ref(-)/purity(-)], across varying tumor spot densities (simulated data). 'perc.' denotes percentage. Panel a (99% percentage) is related to and adapted from Fig. 2c.

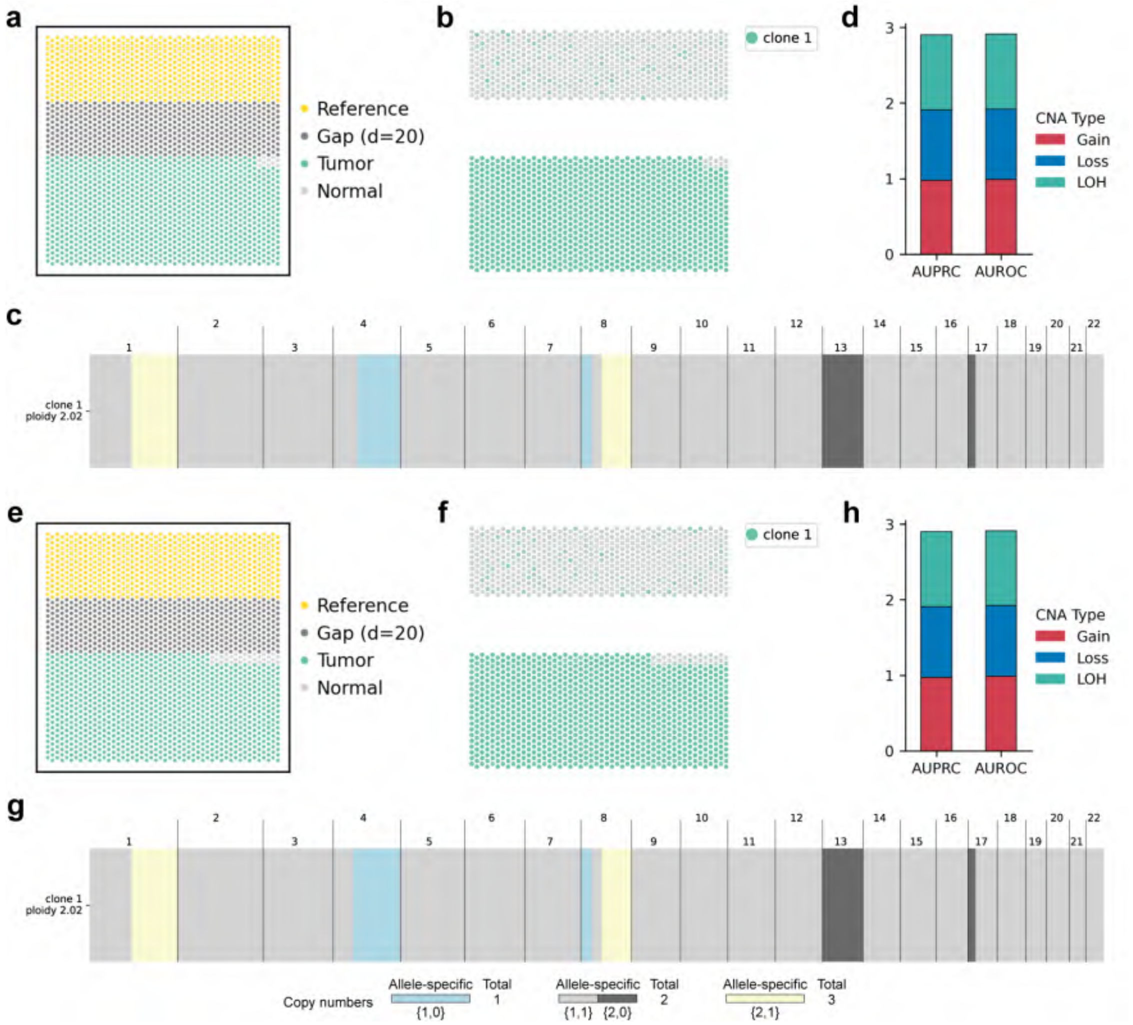


Figure S25: CalicoST CNA prediction performance with spatial gaps between reference and test (normal/tumor) spots under the [ref(+)/purity(+)] setting (simulated data). (a-d) 99% tumor percentage: ground truth (a) and inferred (b) spatial clone distributions; predicted CNA profiles (c); and corresponding AUPRC and AUROC metrics (d). (e-h) 97% tumor percentage: ground truth (e) and inferred (f) spatial clone distributions; predicted CNA profiles (g); and corresponding AUPRC and AUROC metrics (h).

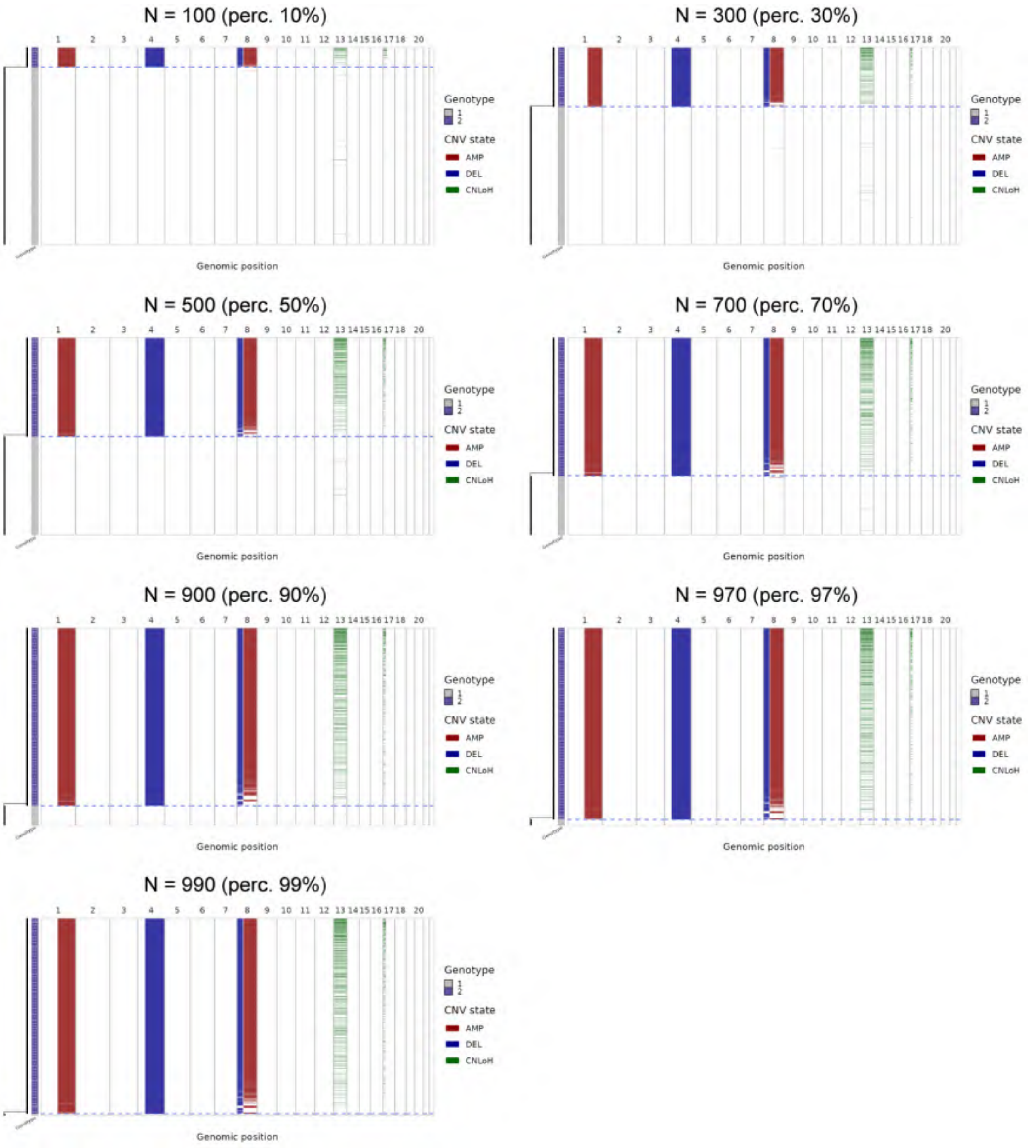


Figure S26: Numbat performance in CNA prediction across varying tumor percentages (simulated data). The method fails to produce outputs at 1% and 3% tumor content because no CNAs are detected at these levels. Note that although reference spots are provided to Numbat, they are not displayed as they were excluded prior to prediction using the `numbat::run_numbat()` function. 'perc.' denotes percentage.

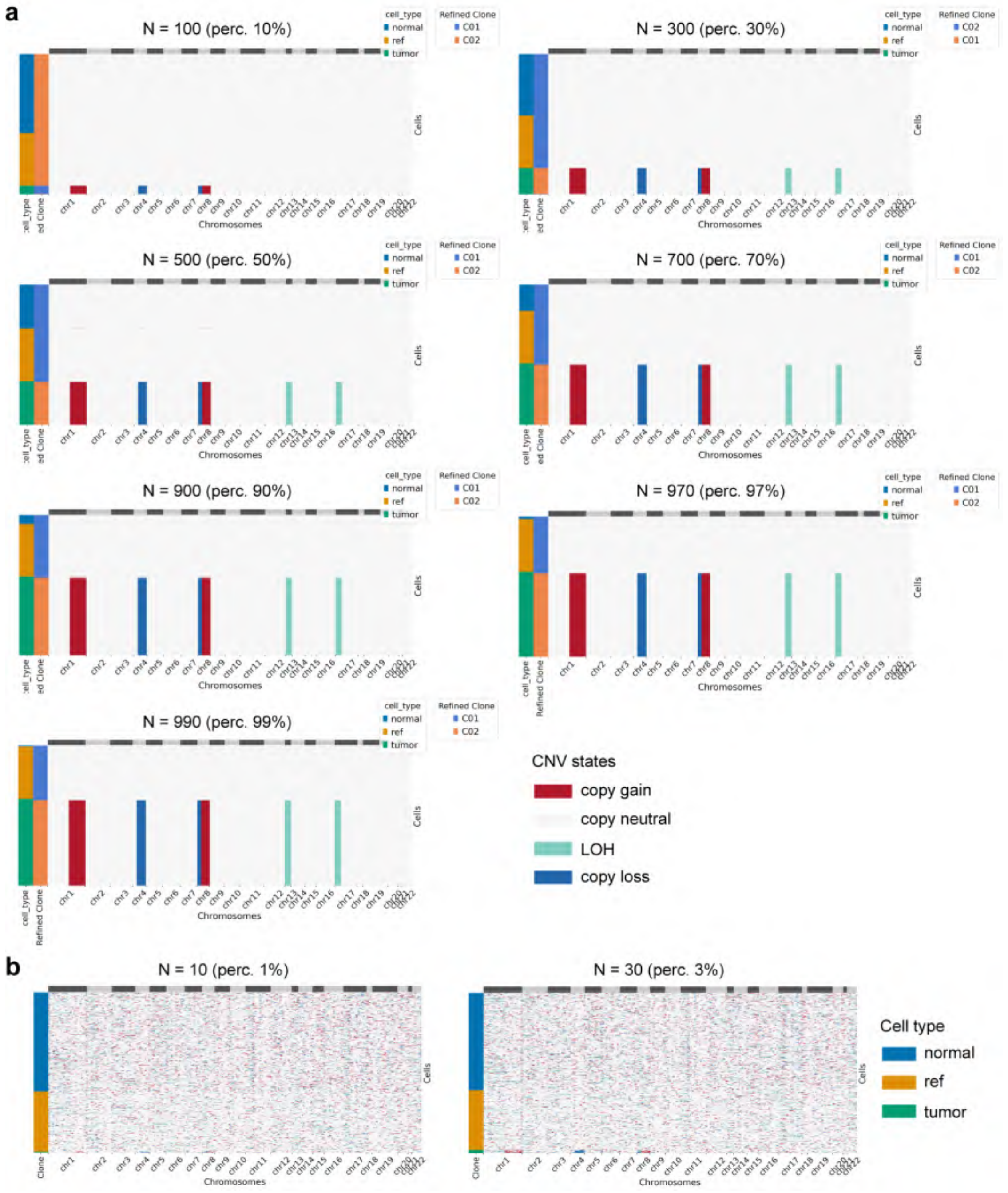


Figure S27: XClone performance in CNA prediction across varying tumor percentages (simulated data). (a) CNA profiles predicted by XClone in refined mode. (b) CNA profiles predicted by XClone base mode at 1% and 3% tumor content, where the refined mode failed to produce outputs. 'perc.' denotes percentage.

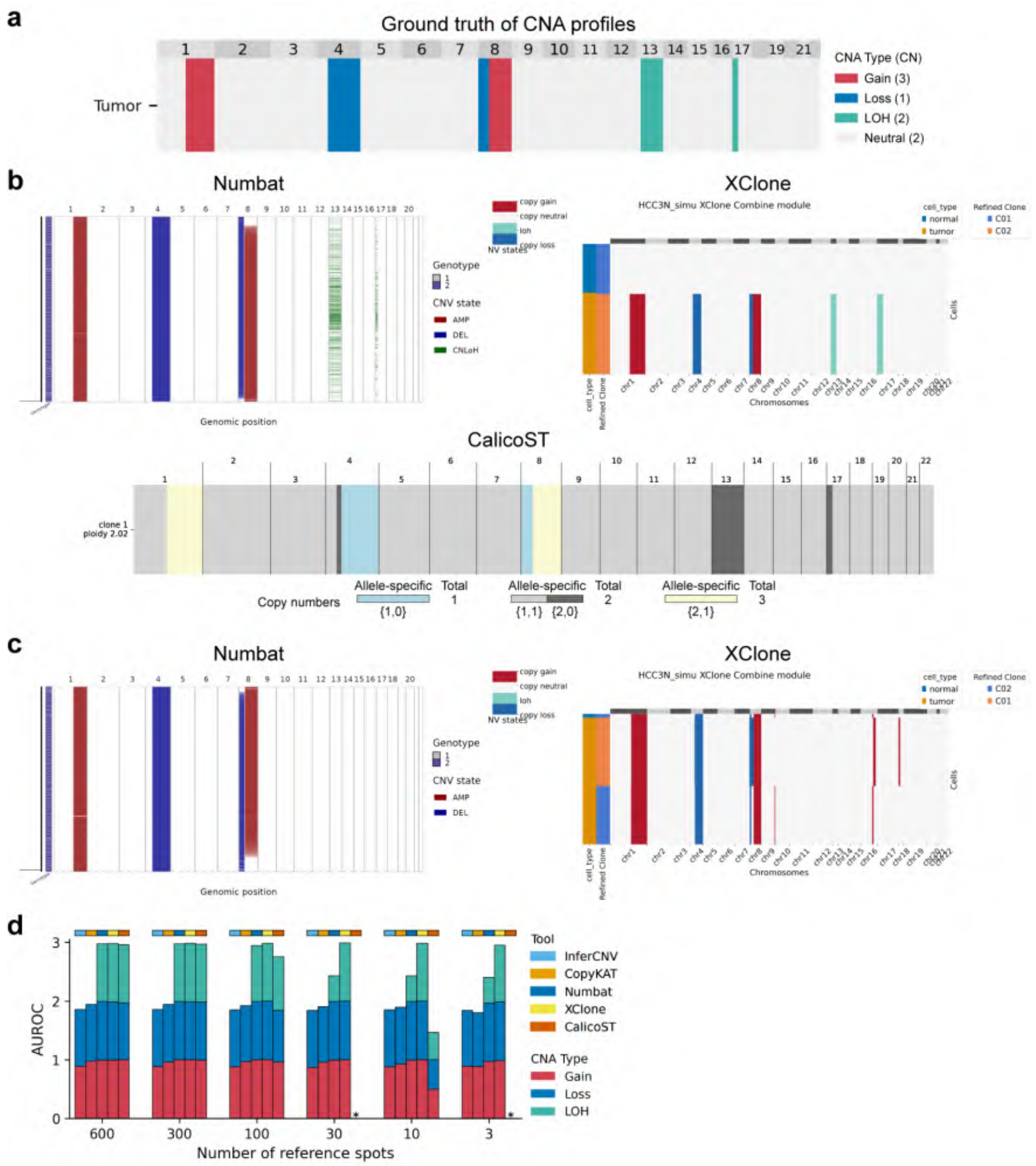


Figure S28: Reference spot downsampling experiment for CNA prediction (simulated data). (a) Ground truth simulated CNA profiles. CN: copy number. (b) CNA predictions by Numbat, XClone, and CalicoST with 600 reference spots. (c) Predictions with 30 reference spots; notably, CalicoST failed to produce a result under this condition. (d) AUROC analysis of predicted CNAs as a function of reference spot abundance. An asterisk (*) indicates no CNA was detected or a technical error occurred.

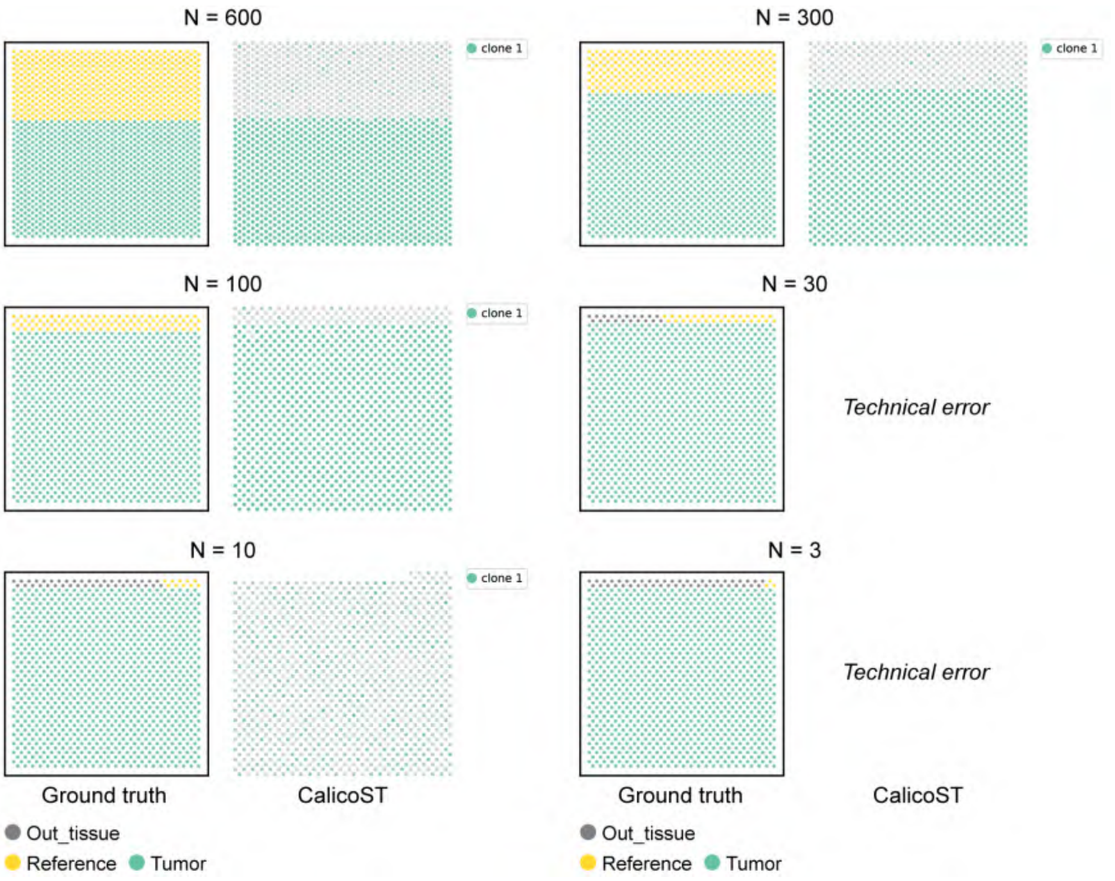


Figure S29: Spatial distribution of ground truth and CalicoST-inferred clones, across a range of reference spot densities (simulated data). While the number of reference spots varies in these simulations, the number of tumor spots remains constant (n = 1000).

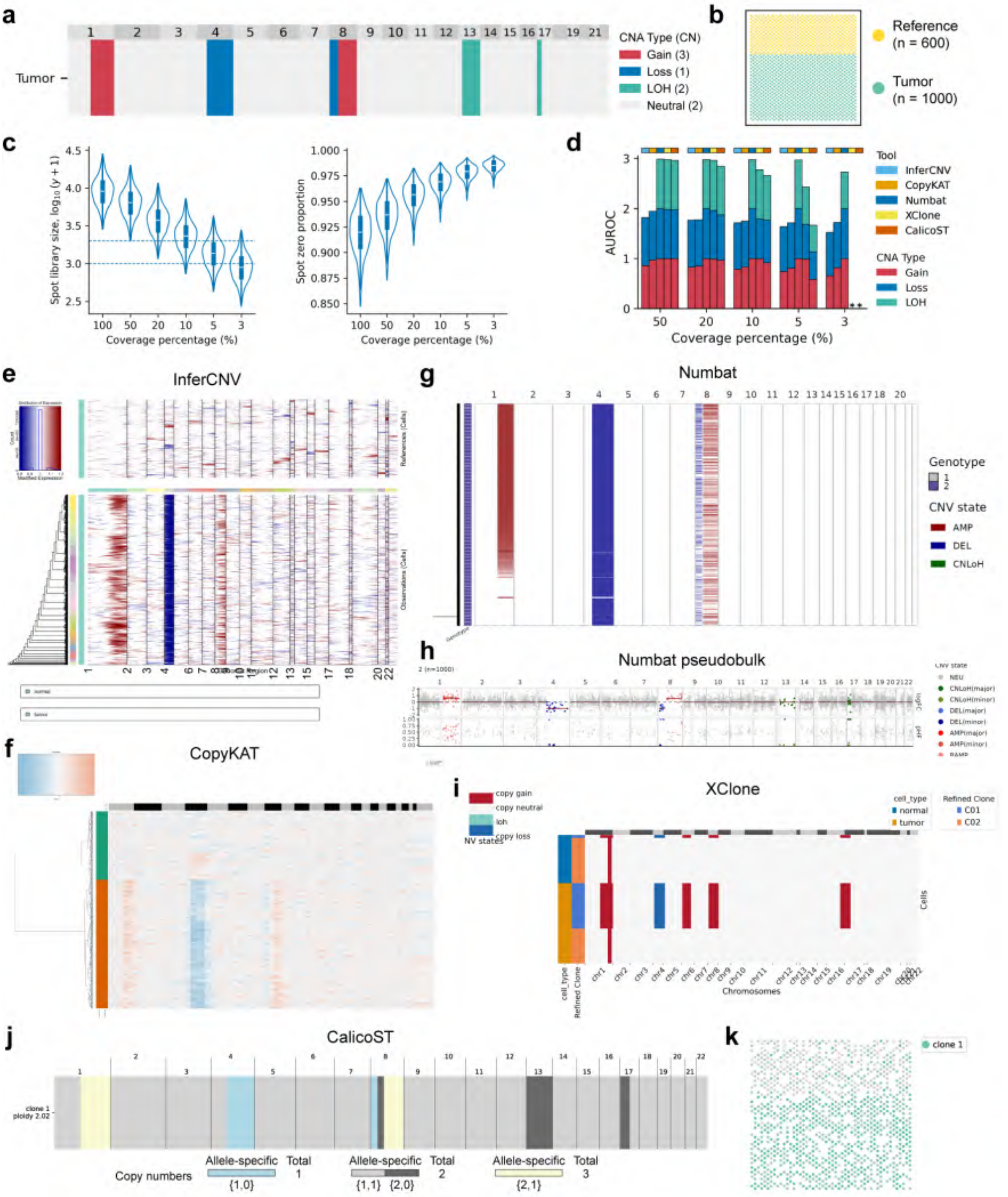


Figure S30: Coverage downsampling experiment for CNA prediction (simulated data). Note that spot library sizes are quantified at the UMI level, whereas coverage down-sampling operates at the read level. (a) Ground truth simulated CNA profiles. CN: copy number. (b) Spatial distribution of ground truth clones. (c) Distribution of spot-wise library size and zero proportion across varying coverage levels. The two blue dashed lines mark library sizes of 1000 and 2000, respectively. (d) AUROC analysis of predicted CNAs as a function of coverage percentage. An asterisk (*) indicates no CNA was detected or a technical error occurred. (e-j) Predicted CNA profiles at 5% coverage: InferCNV (e), CopyKAT (f), Numbat single-cell CNA landscape (g), Numbat final clone pseudobulk CNA profiles (h), XClone (i), and CalicoST (j). (k) Spatial distribution of clones identified by CalicoST at 5% coverage. Panel c (left plot) is related to and adapted from Fig. 2f.

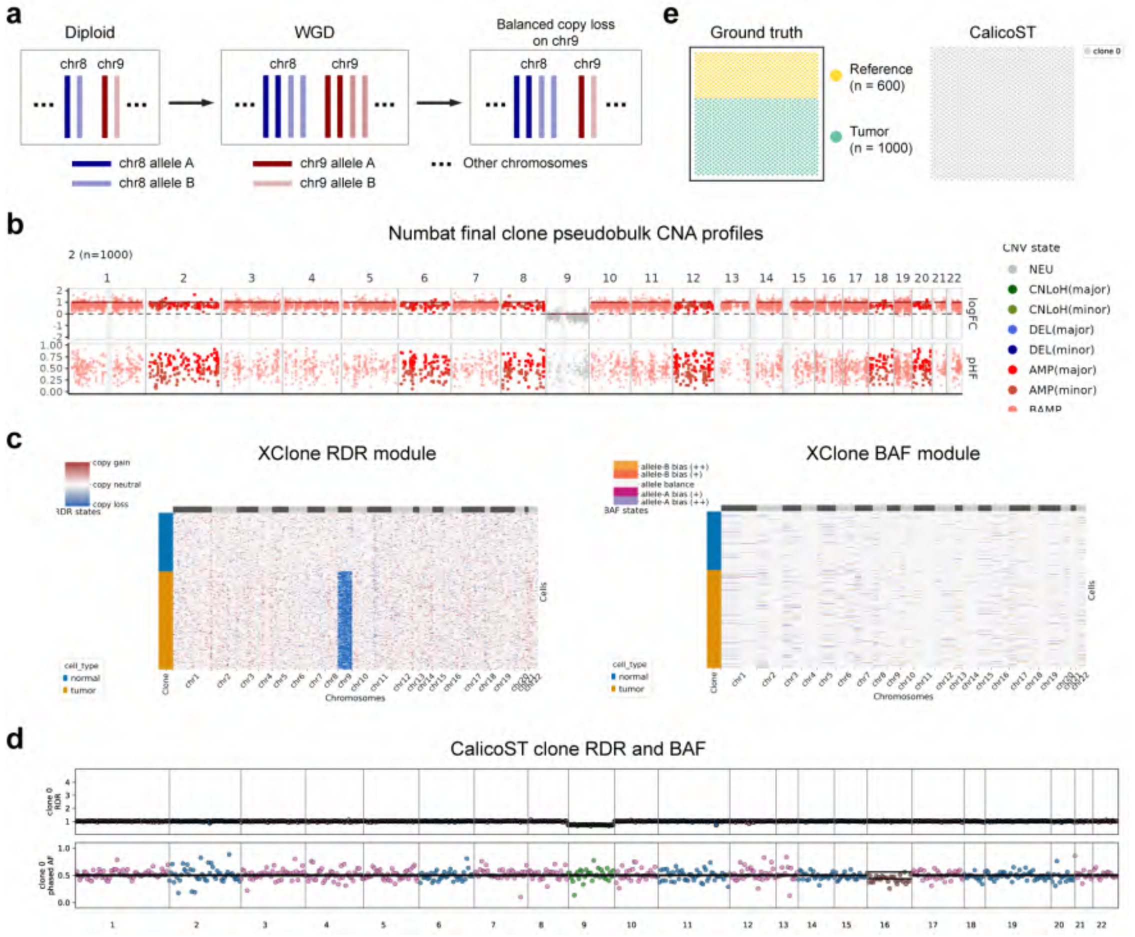


Figure S31: Performance evaluation in predicting whole genome duplication (WGD) followed by balanced copy loss on chr9 (simulated data). Note: for CalicoST, only results under the [ref(-)/purity(-)] setting are shown, as the [ref(+)/purity(+)] setting yielded no output due to technical errors. (a) Schematic illustration of the simulated data. (b) Numbat final clone pseudobulk CNA profiles. (c) XClone RDR and BAF outputs. (d) CalicoST clone-specific RDR and BAF along the genome. (e) Spatial distribution of ground-truth and CalicoST-identified clones.

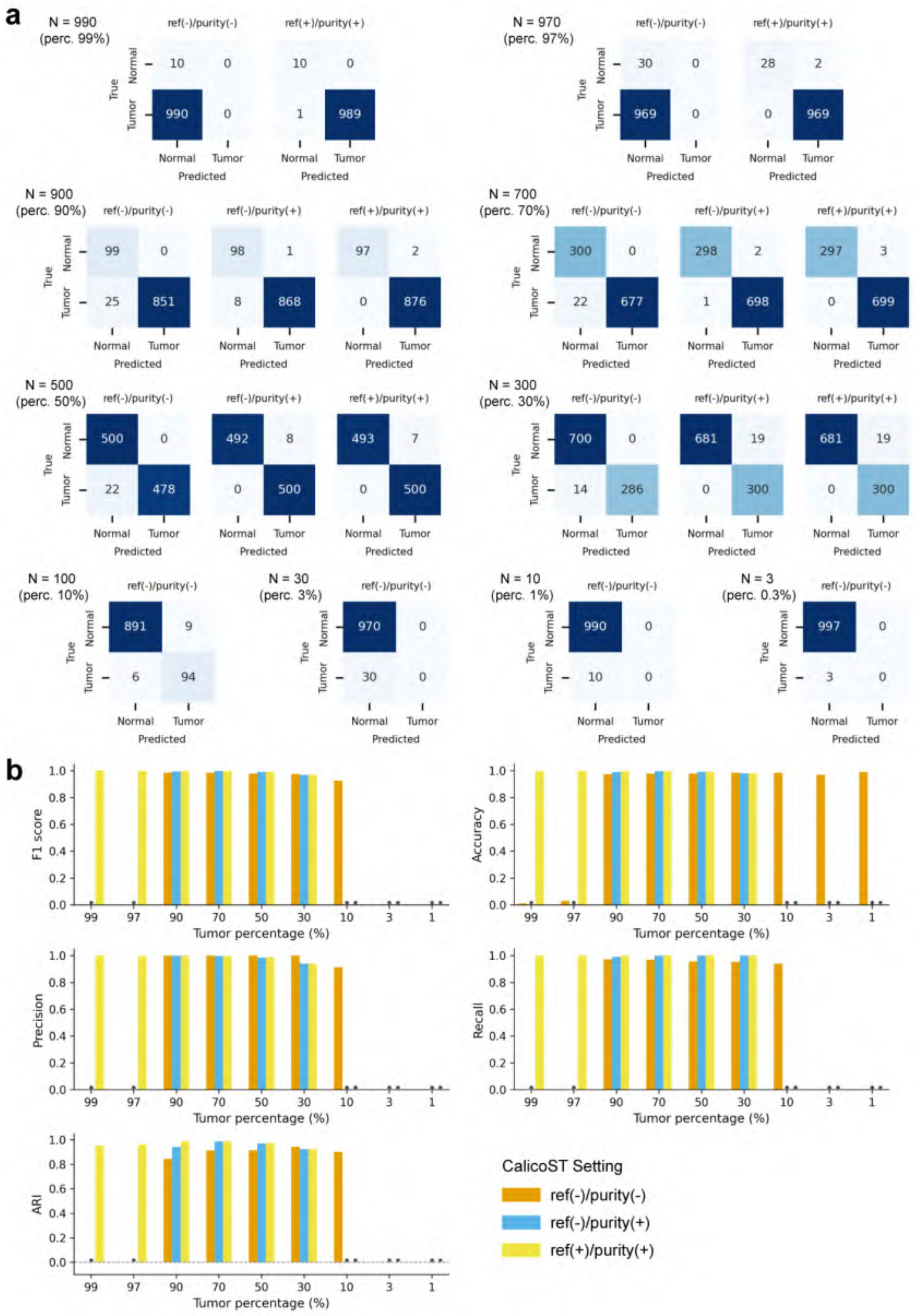


Figure S32: CalicoST performance in tumor identification across various tumor percentages (simulated data). (a) Confusion matrices comparing ground truth and CalicoST-predicted normal/tumor labels for all valid settings. Some results for the [ref(-)/purity(+)] and [ref(+)/purity(+)] configurations are omitted due to technical errors. The number of test spots could be lower than the input count (e.g., at 90% tumor percentage) due to CalicoST internal spot filtering. 'perc.' denotes percentage. (b) Performance metrics for each setting, including F1 score, accuracy, precision, recall, and ARI. 'Tumor' was treated as positive label in metric calculation. An asterisk (*) indicates a technical error occurred.

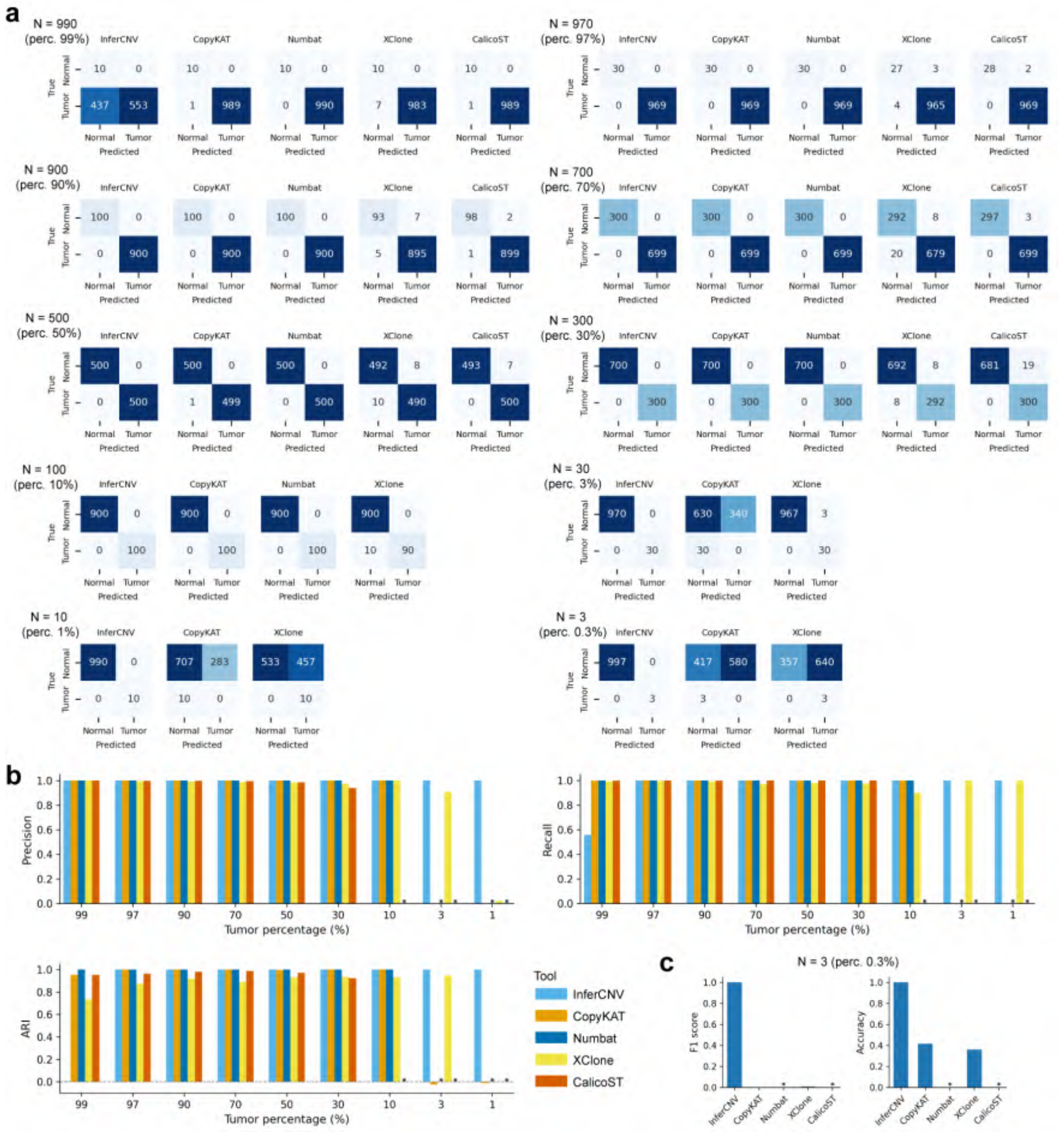

Figure S33: Benchmarking of tumor identification across various tumor percentages (simulated data). All tools use a reference. (a) Confusion matrices comparing ground truth and tool-predicted normal/tumor labels. Results with technical errors are omitted. The number of test spots could be lower than the input count (e.g., at 97% tumor percentage) due to tool's internal spot filtering. (b) Performance metrics, including F1 score, accuracy, precision, recall, and ARI. 'Tumor' was treated as positive label in metric calculation. An asterisk (*) indicates a technical error occurred. (c) Performance at extremely low tumor spot counts (n = 3). 'perc.' denotes percentage.

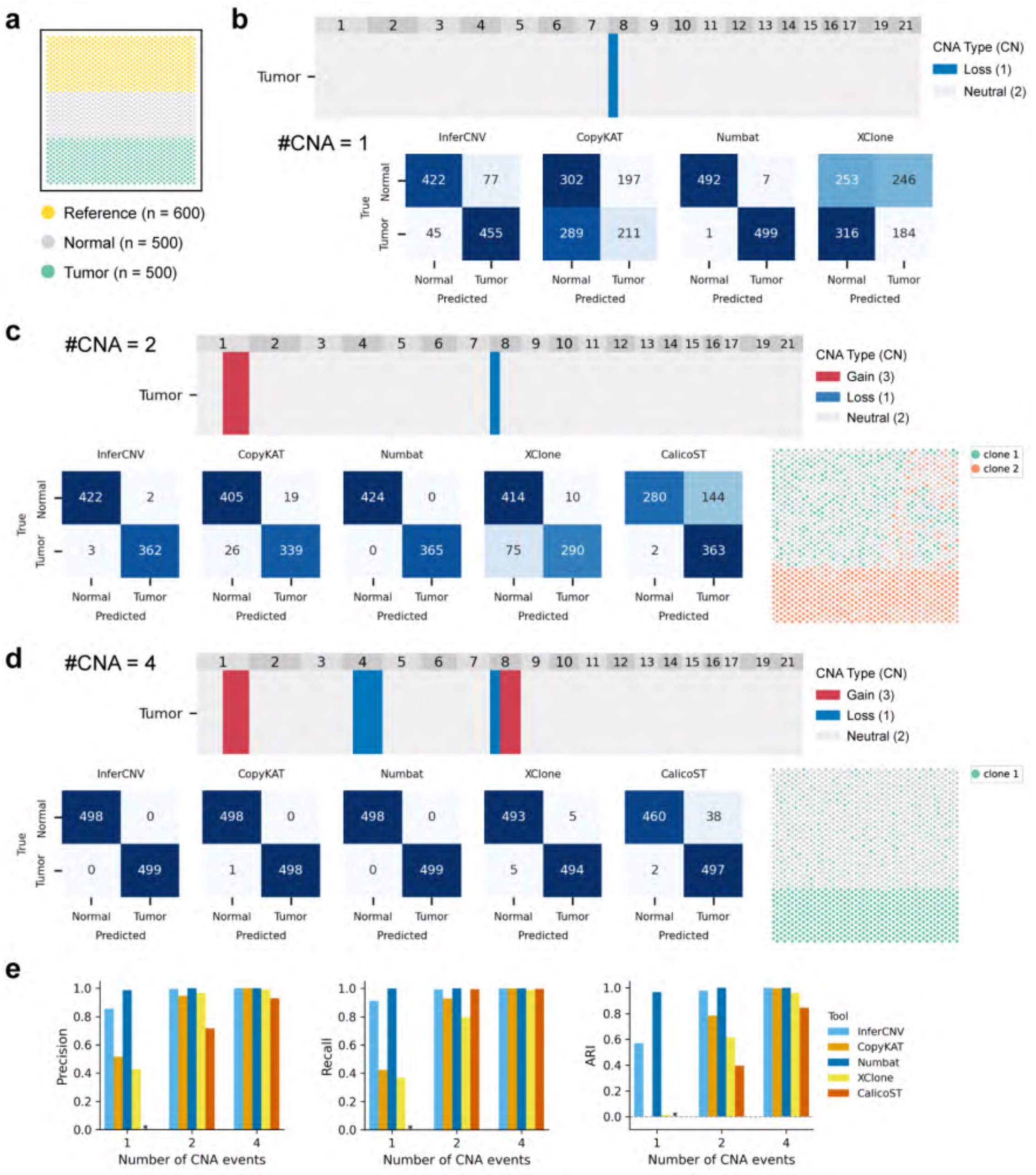


Figure S34: Benchmarking of tumor identification across various CNA profiles (simulated data). (a) Spatial distribution of ground truth clones, used by all simulations in this experiment. (b-d) Results for varying number of CNA events: 1 (b), 2 (c), and 4 (d). Each panel displays the ground-truth CNA profile of the tumor clone, the confusion matrix, and the spatial distribution of clones identified by CalicoST. Note that CalicoST results are not shown in panel (b) due to a technical failure. CN: copy number. (e) Performance metrics for each CNA profile, including precision, recall, and ARI. 'Tumor' was treated as positive label in metric calculation. An asterisk (*) indicates a technical error occurred.

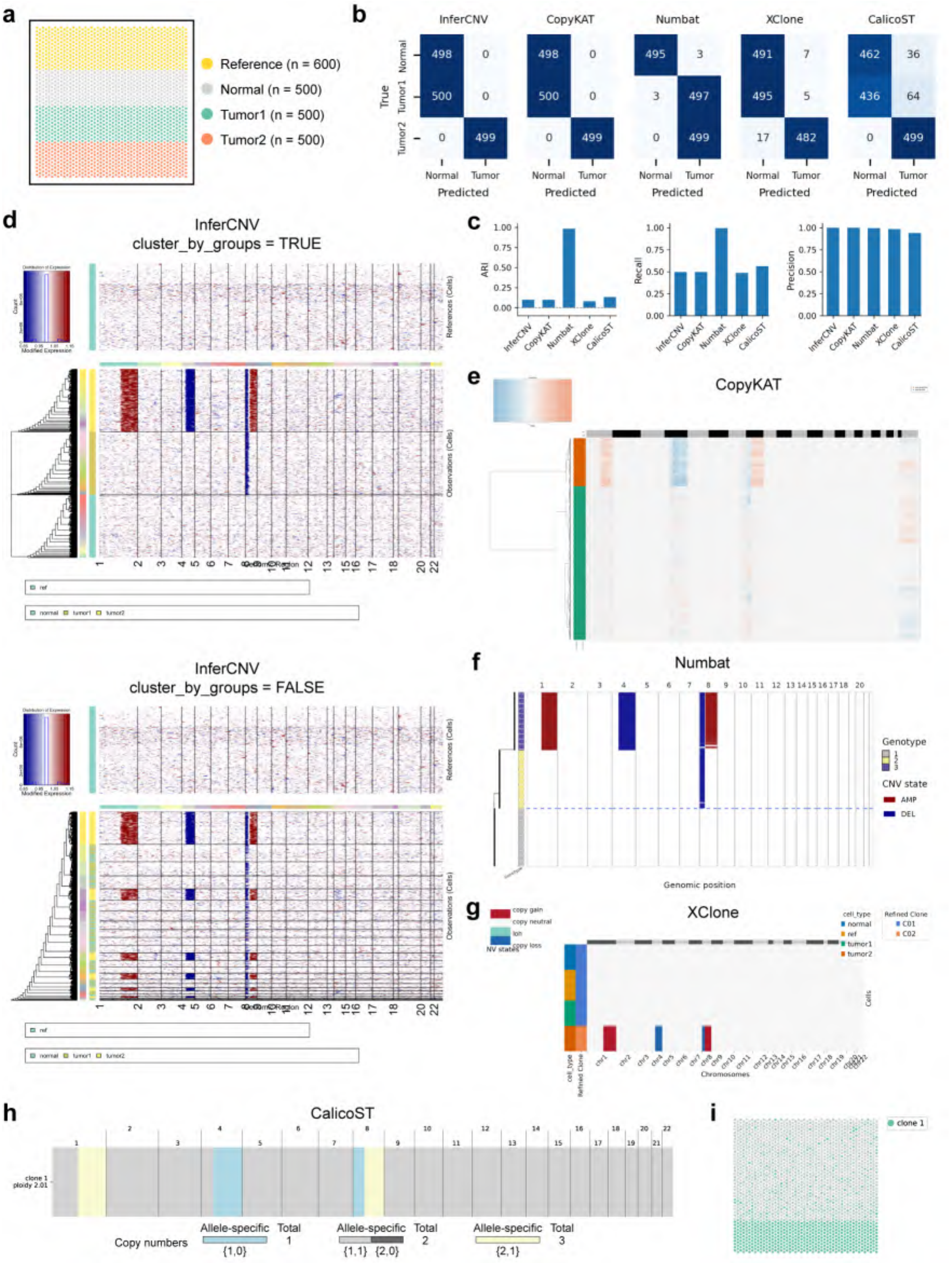


Figure S35: Benchmarking of tumor identification using a dataset with subclonal structure (simulated data). (a) Spatial distribution of ground-truth clones. (b) Confusion matrices comparing ground-truth (fine-grained 3-class clone labels) and predicted normal/tumor labels. (c) Performance metrics (for 2-class labels). 'Tumor' was treated as positive label in metric calculation. An asterisk (*) indicates no CNA was detected or a technical error occurred. (d-h) Predicted CNA profiles by: InferCNV (d), CopyKAT (e), Numbat (f), XClone (g), and CalicoST (h). (d) Here InferCNV `cluster_by_groups` indicates spots are clustered separately within each pre-defined clone (`TRUE`) or clustered together globally as a single unified group (`FALSE`). (i) Spatial distribution of clones identified by CalicoST.

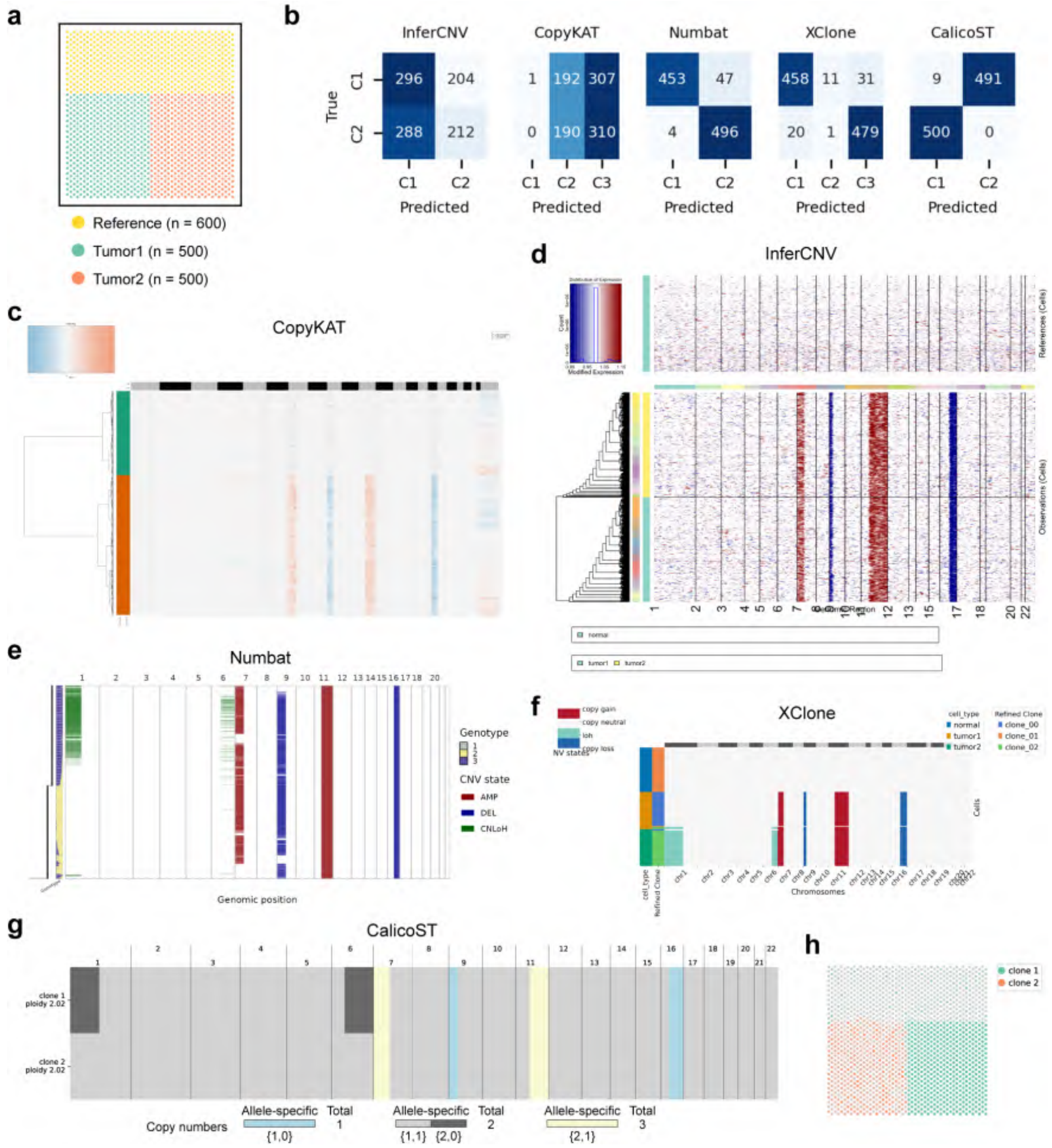


Figure S36: Benchmarking of subclone inference using a dataset with subclonal LOH events (simulated data). (a) Spatial distribution of ground-truth clones. (b) Confusion matrices comparing ground-truth and predicted clone labels. (c-g) Predicted CNA profiles by: CopyKAT (c), InferCNV (d), Numbat (e), XClone (f), and CalicoST (g). (h) Spatial distribution of clones identified by CalicoST.

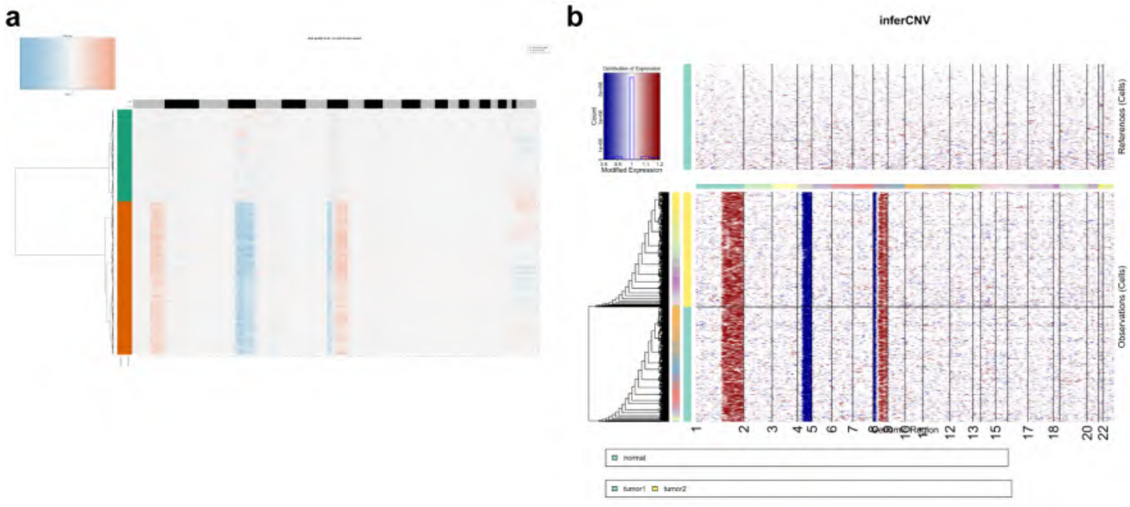


Figure S37: Benchmarking of subclone inference using a dataset with copy loss on distinct alleles (simulated data). Predicted CNA profiles by CopyKAT (a) and InferCNV (b).

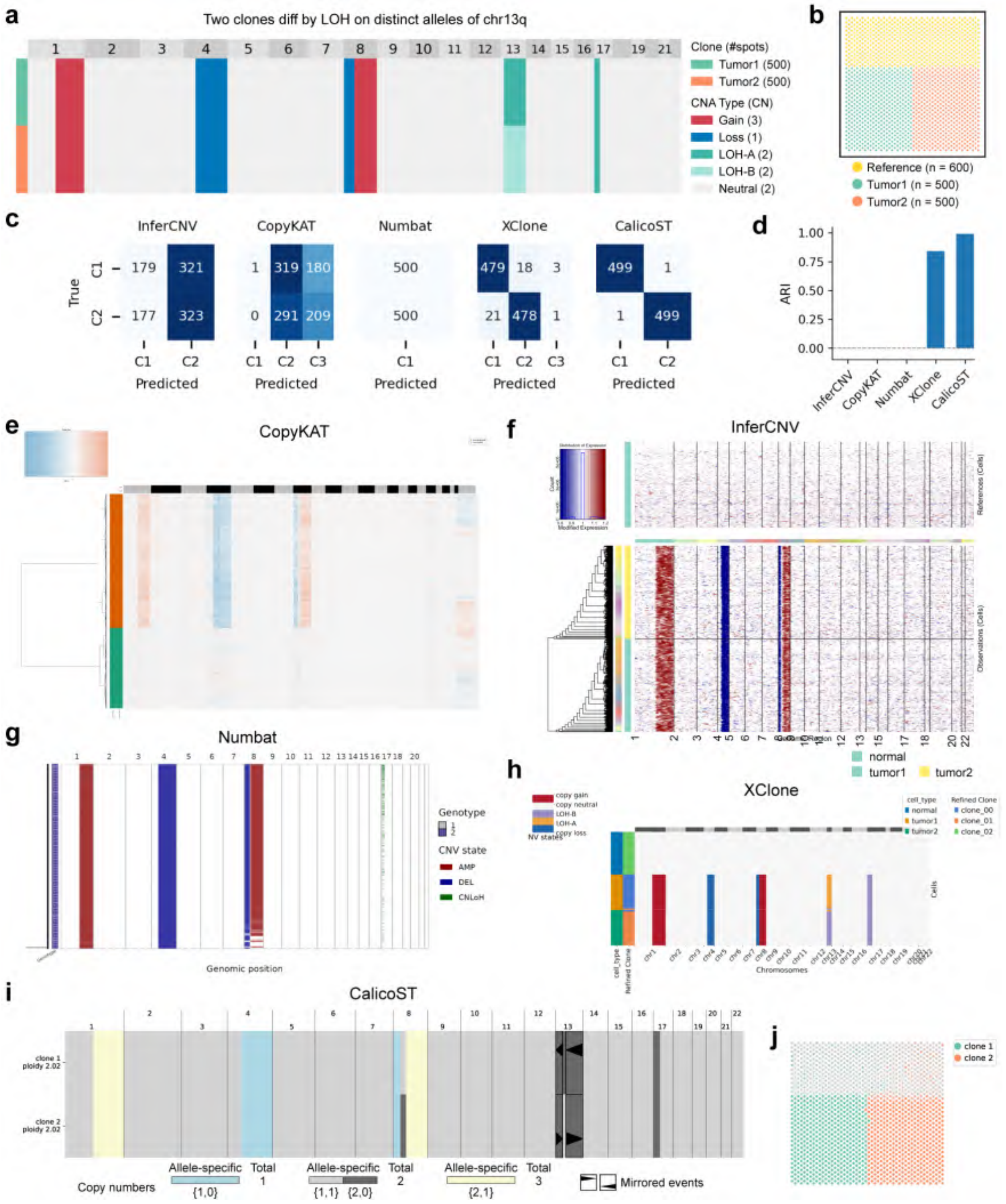


Figure S38: Benchmarking of subclone inference using a dataset with LOH on distinct alleles (simulated data). (a) Ground-truth CNA profile. Two clones diff by LOH events on distinct alleles of chr13q. (b) Spatial distribution of ground-truth clones. (c) Confusion matrices comparing ground-truth and predicted clone labels. (d) ARI metrics for ground-truth versus predicted clone labels. (e-i) Predicted CNA profiles by: CopyKAT (e), InferCNV (f), Numbat (g), XClone (h), and CalicoST (i). (j) Spatial distribution of clones identified by CalicoST.

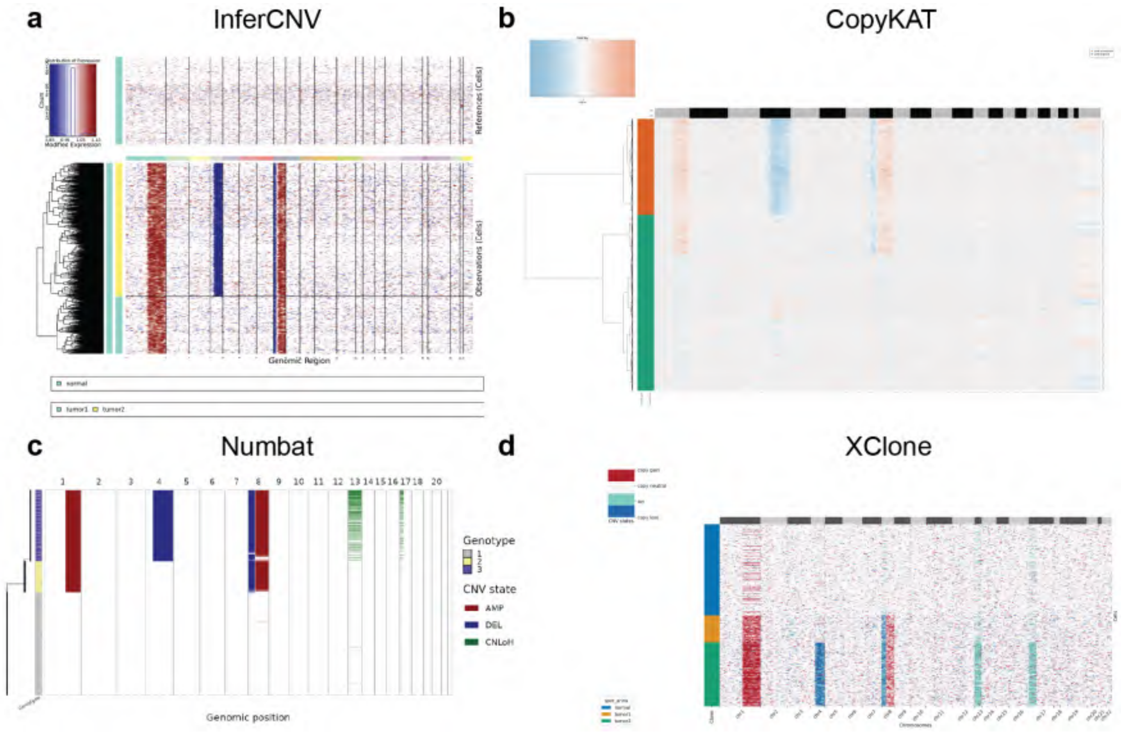


Figure S39: Benchmarking of spatial patterning using simulated data: CNA profiles generated by InferCNV (a), CopyKAT (b), Numbat (c), and XClone (d) as baseline.

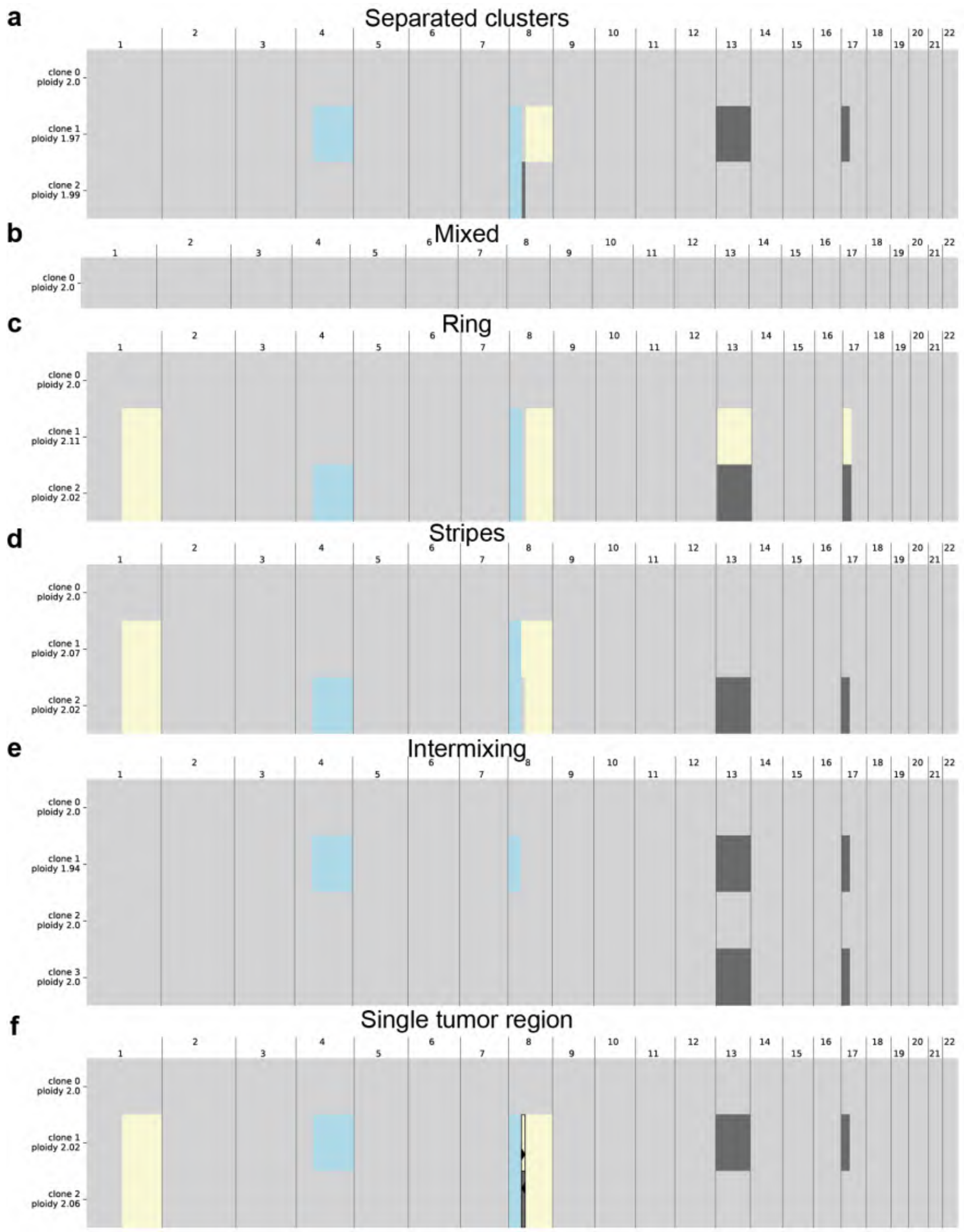


Figure S40: Benchmarking of spatial patterning using simulated data: copy number profiles inferred by CalicoST across diverse tumor clonal geometries. (a–f) Estimated copy number profiles generated by CalicoST under distinct spatial configurations: separated clusters (a), randomly mixed (b), ring (c), stripes (d), intermixing (e), and a contiguous single tumor region (f).

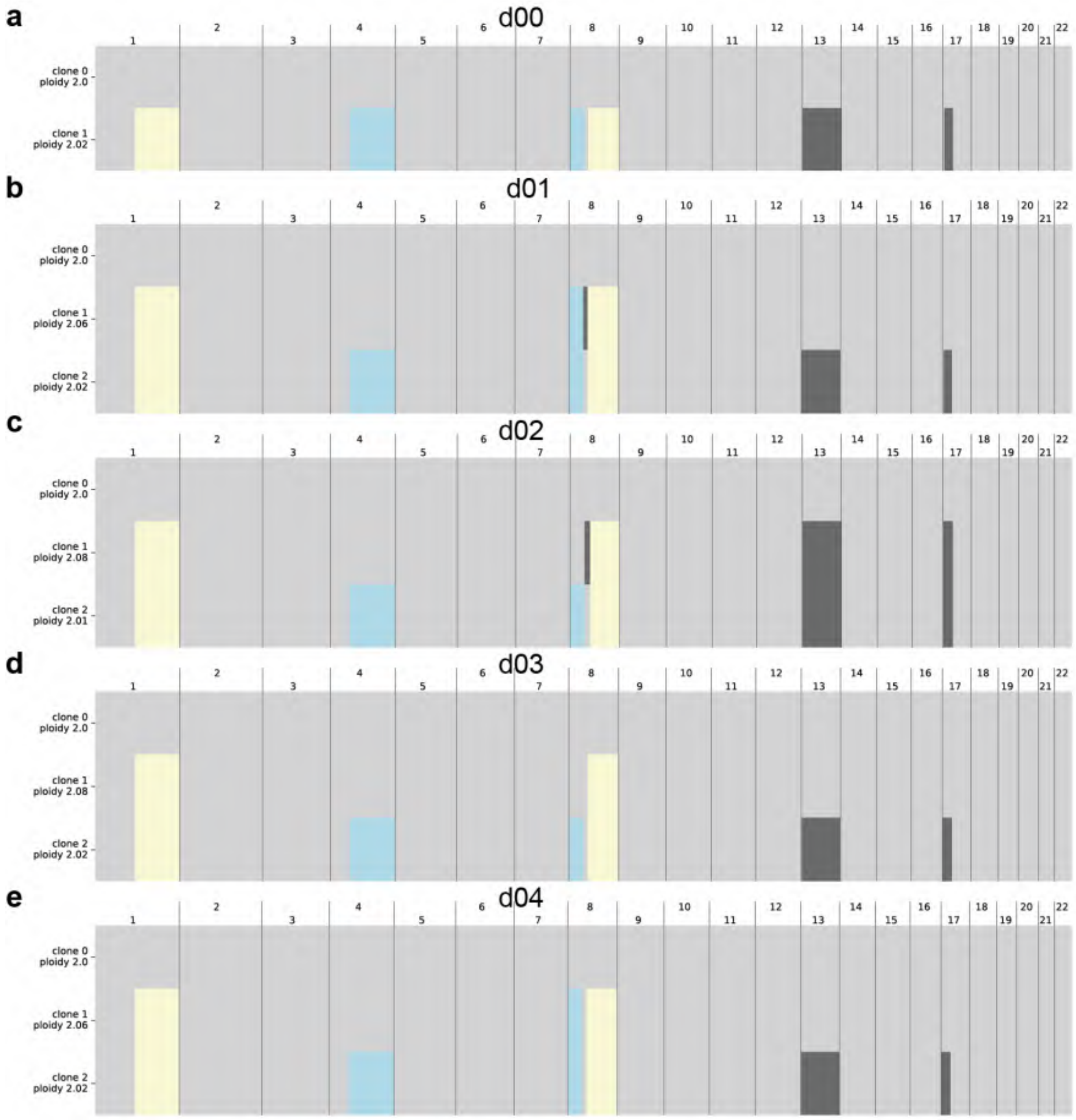


Figure S41: Benchmarking of spatial patterning using simulated data: copy number profiles inferred by CalicoST across varying interclonal spatial distances. (a–e) Estimated copy number profiles generated by CalicoST as the spatial distance between the two distinct tumor sub-clones incrementally increases.

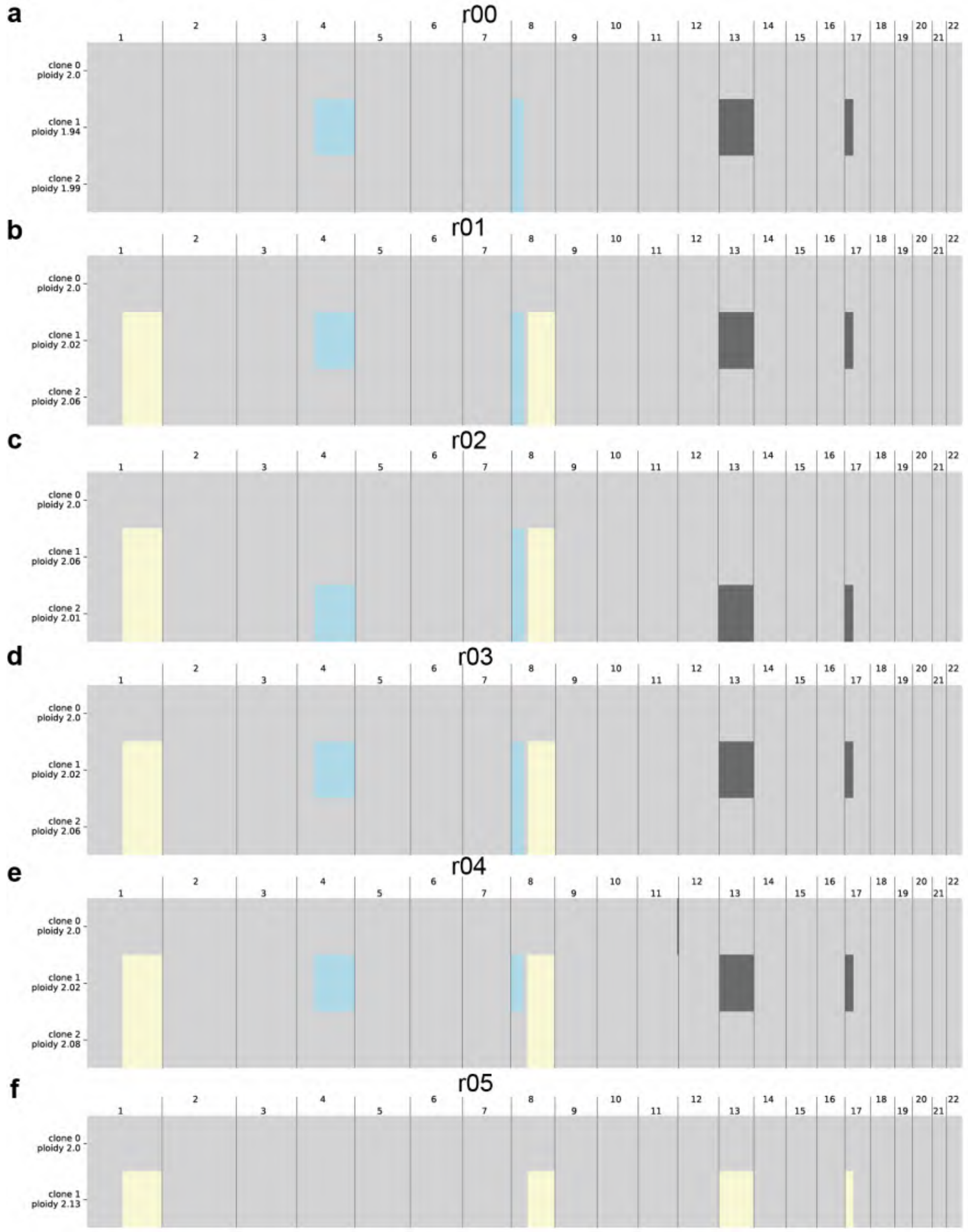


Figure S42: Benchmarking of spatial patterning using simulated data: copy number profiles inferred by CalicoST across varying interclonal cell intermixing rates. (a–f) Estimated copy number profiles generated by CalicoST under progressively increasing rates of interclonal cell mixing.

### 2 Supplementary Notes: stCNASim evaluation

#### 2.1 stCNASim Overview

StCNASim is an allele-aware spatial RNA-seq simulator designed for comprehensive benchmarking of CNA analysis across diverse CNA profiles, clonal structures, and spatial tissue patterns. Its seed data consists of real-world 10x Visium spatial transcriptomic dataset, specifically a subset BAM file [2] derived from annotated normal spots (all tumor spots excluded) paired with sample-specific haplotype data formatted as a list of phased single-nucleotide polymorphisms (SNPs). When supplied with clonal annotations and clonal CNA profiles as simulation parameters, stCNASim ultimately generates synthetic sequencing alignments embedded with user-specified CNA signatures and clonal architectures. These input CNA profiles and clonal labels would serve as *in silico* ground truth for benchmarking CNA detection tools applied to the resulting synthetic BAM files or associated count matrices. The simulator enables configurable spatial patterning by randomly distributing synthetic spots within each user-defined spatial clone compartment, supporting controlled modeling of tumor, normal, and subclonal tissue architectures. Final outputs include a synthetic BAM file, associated gene expression matrix, per-spot spatial coordinates, and spot-level clonal assignment annotations, all formatted to match standard 10x Visium data specifications. More details of the simulation framework can be found in the main text.

#### 2.2 Benchmark internal modules using the stCNASim-validation dataset

As far as we know, no established computational methods support allele-specific feature quantification in spatial transcriptomics data. Therefore, we validated the accuracy of the *afc* module (stCNASim-afc) against two standard total-expression quantification pipelines: SpaceRanger v1.1.0 and STARsolo [3]. Benchmarking was performed on two distinct BAM files: the seed BAM file (Supplementary Fig. S3) and the synthetic BAM outputs from the stCNASim-validation dataset (Supplementary Fig. S4). Since stCNASim-afc generates allele-resolved count matrices while the comparator tools output only total gene expression matrices, we harmonized the data by summing counts across stCNASim-afc's three haplotype groups. This ensured that all three tools yielded spot-by-gene total expression count matrices with matching dimensions and comparable magnitudes, facilitating direct quantitative comparison. Across all benchmarks, stCNASim-afc achieved near-perfect concordance with the two reference tools: linear regression of gene-wise mean expression yielded $R^2 = 1.00$ (p-value $< 0.001$) for all pairwise comparisons. Distributions of spot-wise library size, zero proportion, and gene-wise statistics (mean, variance, coefficient of variation, zero proportion) were virtually identical across tools. Log2 fold-change values were tightly centered around zero for highly expressed genes, with mild deviations from zero for lowly expressed genes when comparing stCNASim-afc and STARsolo. This bias was also observed between SpaceRanger and STARsolo, indicating it arises from inherent differences between STARsolo and 10x official pipelines rather than defects in stCNASim-afc. These results demonstrate that count matrices derived from the *afc* module outputs closely match those generated by well-established tools.

Additionally, we compare allele-specific count statistics for normal spots from the seed and simulated datasets. These metrics were generated by executing the afc module on the corresponding BAM files using 8866 phased SNPs. The two BAMs have identical dimensions, comprising 600 normal spots and 32,295 genes. The fraction of UMIs carrying valid haplotype information (i.e., UMIs assigned to haplotypes A and B) relative to all UMIs (A+B+U) is 4.38% and 4.41% for the seed and simulated data, respectively. When examining all UMIs with resolved haplotype information (i.e., UMIs assigned to A, B, and D), the proportion of D-state UMIs (i.e., UMIs with dual-haplotypes) is 0.56% and 0%, respectively. Complete statistics are provided in Supplementary Data 7.

To validate internal consistency between the *cs* and *rs* modules, we performed benchmarking across both aggregated and allele-resolved count matrices generated by the two modules within the synthetic stCNASim-validation dataset (Supplementary Fig. S5). By design, the *cs* module inherently outputs three allele-specific count matrices corresponding to haplotypes A, B, and U, respectively. In contrast, allele-specific count matrices for the *rs* module were obtained by processing *rs*-derived BAM file via the *afc* module. For both modules, we derived the total gene expression count matrix by summing values across the three allele-specific matrices. Aggregated total gene expression and allele-resolved count matrices generated by the two modules exhibited nearly identical statistical characteristics across both normal and tumor spots. Pairwise linear regression of gene expression mean values for aggregated and allele-stratified (Hap-A, B, U) count matrices yielded $R^2 = 1.00$ (p-value $< 0.001$), with regression slopes equal to 1.00 and negligible intercept terms. All spot-wise and gene-wise metrics fully overlapped, and smoothed trends linking gene-wise mean expression to other summary statistics were highly consistent between the two modules. Fold-change analyses further revealed minimal mean expression differences across the vast majority of genes when comparing outputs from the two modules, for both normal and tumor profiles. Collectively, these results demonstrate that the read-simulation module accurately recapitulates count-level signals produced by the count-simulation module, encompassing both total gene expression and allele-specific expression patterns.

Taken together, these results verify the reliability of the internal *afc* module and high concordance between outputs generated by the *cs* and *rs* modules. We next benchmark the count and read simulation performance of *cs* and *rs* modules against external pipelines in subsequent analyses.

#### 2.3 Benchmarking the *cs* module's count simulation performance against scDesign2-cna

We evaluated the count-simulation performance of stCNASim's *cs* module (stCNASim-cs) against a custom CNA-aware pipeline (scDesign2-cna) by comparing their simulated profiles to the seed data across multiple evaluation metrics. The original standard scDesign2 pipeline [4] lacks native CNA simulation functionality. To enable a fair comparison against the *cs* module, we modified scDesign2's independent mode to integrate CNAs via a straightforward procedure outlined in Supplementary Fig. S6, and named this revised workflow scDesign2-cna. Briefly, scDesign2-cna first simulates baseline gene expression counts using the core scDesign2 independent mode. It then rescales each gene's baseline UMI counts by gene-wise copy number (CN) ratios matched to the assigned CN state - specifically, ratios $<1$ for copy loss, $=1$ for copy neutral, and $>1$ for copy gain - thereby introducing transcriptional shifts driven by copy number changes. Notably, the resulting synthetic count matrix reflects total gene expression rather than allele-specific data.

For evaluation, the total gene expression count matrix generated by stCNASim-cs (aggregated from its three synthetic allele-specific matrices derived from the stCNASim-validation dataset) was benchmarked against two profiles: (1) the seed total-expression matrix, obtained by aggregating the three allele-specific outputs from the stCNASim-afc module on the seed BAM; and (2) the synthetic total-expression matrix output by scDesign2-cna. This tool adopted the previously described seed matrix as its template and used the CNA profiles extracted from the stCNASim-validation dataset as simulation parameters, yielding a synthetic dataset containing 600 normal and 600 tumor spots - exactly matching the spot count of the stCNASim-validation dataset. This setup maximizes consistency by ensuring that both simulators utilize input (allele-specific or aggregated matrix) templates quantified by the stCNASim-afc module.

Both stCNASim-cs and scDesign2-cna faithfully recapitulated count-derived expression features observed in the seed data (Supplementary Fig. S7). Pair-wise linear-regression analysis on gene-wise mean expression yielded near-perfect correlation ($R^2 = 1.00$, p-value $< 0.001$) across all genes for both normal and tumor spots. A high concordance was also observed for relationships between gene-wise mean expression and other summary statistics. Regarding spot-level metrics (library size and zero proportion), both simulators accurately captured reference median values, though scDesign2-cna exhibited a significantly narrower and more tightly centered distribution than the seed dataset.

For CNA-associated evaluations, we stratified genes by their distinct copy-number states for separate downstream analyses (Supplementary Fig. S7). Across all copy-neutral genes, gene-wise distributions of mean expression, variance, and coefficient of variation derived from stCNASim-cs and scDesign2-cna synthetic profiles closely matched those of the seed data. When examining read-depth-ratio (RDR) signatures under different CNA conditions, log2 fold-change analyses - an established RDR proxy routinely used in CNA detection workflows - confirmed that both tools reproduced the anticipated transcriptional patterns. Specifically, values clustered tightly around 0 for copy-neutral and LOH genes (CN=2), centered near 0.58 for copy-gain events (CN=3), and approximated -1 for copy-loss events (CN=1). Nevertheless, the two pipelines diverged markedly in gene-wise zero-proportion distributions: only stCNASim-cs generated realistic zero-fraction profiles within CNA-altered genomic regions, demonstrating its superior performance in modeling expression dropout and zero inflation under copy-number shifts.

Pseudobulk log2 fold-change (log2FC) profiles contrasting tumor and normal spots generated from stCNASim-cs and scDesign2-cna simulations were plotted along chromosomes 1-22 in Supplementary Fig. S8. These log2FC metrics were initially computed directly from raw UMI counts. We additionally assessed two alternative log2FC quantification strategies: (1) log2FC values derived from counts normalized to the median spot library size that were estimated separately across all simulated spots for each pipeline; (2) re-centering of the aforementioned normalized log2FC values, performed by subtracting the median log2FC calculated exclusively from copy-neutral genes. Raw log2FC, median-library-normalized log2FC, and neutral-gene-recentered log2FC all displayed highly concordant genome-wide trends between the two simulation pipelines. Both tools faithfully recapitulated the characteristic transcriptional shifts matching the ground-truth copy number states.

We further evaluated stCNASim-cs under a challenging simulation scenario engineered to introduce strong compositional biases within the synthetic dataset (Supplementary Fig. S9). In this test dataset, tumor spots contained 8 copy-gain events, 2 copy-loss events, and 2 LOH events, creating a pronounced global transcriptional skew toward elevated expression from copy amplification. Log2FC values calculated from raw counts exhibited canonical CNA-associated expression patterns centered at their theoretical values. In contrast, log2FC estimates computed from library-size-normalized counts suffered from global signal shifts, downward in this case, driven by compositional distortion. However, this systematic bias could be fully mitigated by the neutral-gene re-centering correction outlined above. Collectively, these findings demonstrate stCNASim's robustness in preserving distinct, interpretable CNA expression signatures even within datasets carrying severe compositional effects.

In summary, while both stCNASim-cs and scDesign2-cna recapitulate near-identical gene-level and spot-level statistics relative to the seed dataset, alongside faithful genome-wide CNA-associated RDR signals, stCNASim-cs uniquely generates realistic zero-proportion distributions within copy-number-altered genomic regions. This advantage renders it more appropriate for benchmarking spatial CNA detection pipelines.

#### 2.4 Benchmarking the *rs* module's read simulation performance against scReadSim-cna

We benchmarked the read-simulation performance of stCNASim's *rs* module (stCNASim-rs) against scReadSim-cna, a custom CNA-aware read simulator, by contrasting their simulated outputs against the seed dataset across multiple evaluation metrics, with special focus on RDR and BAF signatures. Standard scReadSim [1] is a single-cell RNA-seq read simulator that generates synthetic sequencing data from empirical read templates and gene-level count models. To avoid ambiguous read quantification, it partitions the reference genome into two non-overlapping feature classes: merged super-genes for overlapping coding loci, and inter-genes for all remaining intergenic intervals. Its workflow first extracts super-gene and inter-gene UMI count matrices from input seed BAM file, calls external count simulators (e.g., scDesign2) to generate synthetic UMI counts, and subsamples raw experimental reads to produce synthetic FASTQ files. An optional Bowtie2 [5] alignment step converts FASTQs to aligned BAMs. Critically, native scReadSim lacks native support for CNA signal injection and haplotype-resolved BAM generation.

**Benchmark CNA variants of the scReadSim workflow**

To build a CNA-aware baseline simulator, we engineered scReadSim variants by adjusting its core workflow configurations and then compared their performance (Supplementary Fig. S10). First, we replaced scReadSim's default scDesign2 count simulation module with either scDesign2-cna or stCNASim-cs to enable CNA injection. For a fair comparison between the two configurations, we provided both modified simulation pipelines with consistent input materials: a shared total count matrix as a template (quantified from the seed BAM via scReadSim), identical CNA profiles derived from the stCNASim-validation dataset, and matched numbers of tumor and normal spots. We subsequently compared the resulting synthetic count matrices for super-gene and inter-gene features output by each modified workflow. Both modified setups successfully recapitulated the anticipated CNA-associated RDR signatures, which appeared as tumor-vs-normal log2 fold-change (log2FC) shifts consistent with ground-truth copy gain, loss, and LOH events spanning super-gene and inter-gene regions. However, scDesign2-cna produced floating-point count matrices with incompatible data types that crashed downstream read sampling, whereas stCNASim-cs output valid integer counts fully compatible with scReadSim. We therefore selected stCNASim-cs exclusively for CNA-aware count generation within the modified scReadSim pipeline.

We next swapped scReadSim's default Bowtie2 aligner with CellRanger for FASTQ-to-BAM mapping. Unlike Bowtie2, CellRanger generates BAMs with mandatory `xf` tags required by stCNASim-afc for read filtering. We quantified three distinct BAM files to compare the two alignment backends: (1) BAM file generated by aligning synthetic FASTQs (simulated via the modified scReadSim pipeline using stCNASim-cs-derived counts) with CellRanger; (2) BAM file generated by aligning the same synthetic FASTQs using Bowtie2; (3) Original seed BAM. All three BAMs were processed with STARsolo [3] feature counting to produce spot-by-gene total expression matrices. Spot-wise library size and zero proportion distributions, alongside gene-wise zero fractions, were nearly indistinguishable between the two aligners. Linear regression of gene-wise mean expression yielded pairwise correlation $R^2 \geq 0.98$ (p-value $< 0.001$), and log2 fold-change analysis confirmed minimal quantification divergence between Bowtie2- and CellRanger-processed data.

We named this dual-modified pipeline scReadSim-cna, with its two core adjustments summarized in Supplementary Fig. S11: (1) stCNASim-cs replaces the native scDesign2 count simulator, enabling injection of user-defined ground-truth CNA signals into synthetic feature count matrices; (2) CellRanger replaces Bowtie2 for FASTQ alignment to generate BAMs carrying `xf` tags required for stCNASim's allele-specific feature counting module.

**Benchmark stCNASim-rs RDR signals**

To benchmark RDR and BAF signatures from stCNASim against scReadSim-cna, we generated a synthetic dataset using the scReadSim-cna pipeline, hereafter referred to as the scReadSim-cna-validation dataset. Briefly, scReadSim-cna took the seed BAM as its input template and adopted CNA profiles from the stCNASim-validation dataset as simulation parameters, outputting a synthetic BAM file containint 600 normal and 600 tumor spots to precisely match the spot count of the stCNASim-validation dataset. Unsurprisingly, no haplotype information (including phased SNP panel) was supplied to scReadSim-cna as input for this simulation.

We then compared STARsolo-quantified total-expression matrices derived from stCNASim-rs and scReadSim-cna synthetic BAMs against the seed dataset (Supplementary Fig. S12). At the count matrix level, stCNASim-rs and scReadSim-cna produced nearly identical spot-wise library size and zero-proportion distributions, as well as matching gene-wise mean expression, variance, coefficient of variation, and zero fractions, all closely recapitulating seed data statistics. Pairwise linear regression demonstrated strong concordance ($R^2 \geq 0.97$, p-value $< 0.001$) in gene-wise mean expression for both normal and tumor spots. This strong alignment extended to the relationships between mean expression and all other evaluated gene-level metrics. After stratifying genes by four CNA states (neutral, gain, loss, LOH), both tools recovered consistent zero fraction distributions and log2FC values across every subgroup.

For pseudobulk read-depth ratio (RDR) signatures, we calculated tumor-vs-normal log2FC under three standard preprocessing schemes: raw counts (Fig. 1e), median-library-normalized counts, and normalized counts recentered to the median log2FC of all copy-neutral genes (Supplementary Fig. S13). Across all three schemes, stCNASim-rs and scReadSim-cna generated highly concordant genome-wide log2FC landscapes, with elevated expression in copy-gain regions, suppressed expression in copy-loss regions, and intermediate shifts for LOH segments, consistent with the input ground-truth CNA profiles.

**Benchmark stCNASim-rs gene-level BAF signals**

We evaluated BAF performance at both gene and SNP resolution, as both layers encode haplotype information within our simulation framework. For gene-level allelic depth and allele frequency (AF) analysis (Supplementary Fig. S14), we compared spot-by-gene depth (DP) matrices derived from the seed, stCNASim-validation, and scReadSim-cna-validation datasets. Each DP matrix was computed by aggregating haplotype A and haplotype B allele-resolved counts generated by stCNASim-afc from the corresponding BAM file of each dataset. Phased AF was computed as the ratio of haplotype B counts to total DP, and assigned to `NaN` for entries with zero DP.

Both simulators produced matching distributions of mean gene DP and DP zero fractions across all four CNA strata. Nevertheless, phased AF profiles from the three pipelines exhibited stark divergence: (1) seed reference data showed AF values centered at 0.5 for both normal and tumor spots; (2) stCNASim-rs generated biologically realistic phased AF distributions matching theoretical allelic imbalance signatures for each CNA category. In tumor spots, AF values clustered around 0.5 for neutral loci, 0 or 1 for copy loss and LOH segments, and 0.33 or 0.67 for CN=3 gains; all genes displayed AF centered at 0.5 in normal spots. (3) in sharp contrast, scReadSim-cna produced phased AF values heavily skewed toward 0 or 1 across all genes in both normal and tumor spots. This systematic bias constitutes a technical artifact caused by the simulator's exclusive dependence on one single reference haplotype for BAM construction. Linear regression quantified this gap: stCNASim-rs AF correlated strongly with seed profiles ($R^2 = 0.66$), while scReadSim-cna AF exhibited near-zero correlation ($R^2 = 0.03$).

Genome-wide pseudobulk phased AF tracks for normal and tumor spots (Supplementary Fig. S15) further illustrated this divide. stCNASim-rs faithfully reproduced tumor-specific allelic shifts at simulated gain, loss, and LOH loci; scReadSim-cna displayed uniform AF polarization across all chromosomal segments and failed to resolve CNA-specific allelic signatures.

**Benchmark stCNASim-rs SNP-level BAF signals**

We next examined SNP-level BAF signals. First, we characterized the expression statistics of the 8,866 input phased SNPs using pileup outputs from cellsnp-lite [6] mode 1a on both the seed and simulated BAM files (Supplementary Data 8). Specifically, after filtering reads to retain only entries with valid `xf` tags, we recovered 4,064 SNPs with aggregated DP $\geq$ 1 in the stCNASim-rs synthetic BAM; corresponding counts were 4,151 for the seed dataset and 4,068 for scReadSim-cna. However, scReadSim-cna yielded only 377 SNPs with bi-allelic expression (defined as at least one supporting UMI for each allele), compared with 3,171 for the seed and 2,943 for stCNASim-rs. This observation further demonstrates that reads output by scReadSim-cna originate exclusively from a single reference haplotype, limiting its capacity to generate authentic heterozygous SNP profiles.

We then quantified statistics for *de novo* SNPs identified via cellsnp-lite pileup outputs in *de novo* calling mode (mode 2a, parameters: `minCOUNT=11`, `minMAF=0.1`) from seed and simulated BAMs (Supplementary Data 9). We counted SNPs whose chromosome, position, reference, and alternate alleles matched the original 8,866 phased SNP panel: 1,477 SNPs in the seed, 2,045 in stCNASim-rs, and just 1 in scReadSim-cna. The single recovered SNP from scReadSim-cna is likely a technical artifact. This implies the 377 bi-allelic SNPs detected in scReadSim-cna under cellsnp-lite mode 1a most likely stem from the pipeline's artificial sequencing error introduction module rather than genuine heterozygous variation. Collectively, these results confirm that all *de novo* SNPs recovered from stCNASim derive from the input phased SNP panel, validating the accuracy of the SNP masking procedure implemented during stCNASim read simulation.

Next, we extracted raw unphased SNP AF (ratio of alternative allele to total DP) via cellsnp-lite for a filtered panel of 7,381 phased SNPs residing within copy-neutral genomic regions (Supplementary Fig. S16). Seed and stCNASim-rs datasets displayed continuous AF distributions centered at 0.5 for normal loci across all spots, whereas scReadSim-cna AF values collapsed near zero, introducing pervasive allelic bias. Pairwise regression again demonstrated strong seed concordance for stCNASim-rs ($R^2 = 0.66$) and negligible concordance for scReadSim-cna ($R^2 = 0.02$), confirming scReadSim-cna cannot recapitulate authentic unphased SNP allelic signals due to its single-reference-haplotype architecture.

We extended this comparison to the full set of 8,866 genome-wide phased SNPs stratified by CNA state (Supplementary Fig. S17). All BAMs were genotyped with cellsnp-lite mode 1a using the input phased SNP panel; SNP-wise phased AF was calculated by flipping raw AF values according to each variant's assigned haplotype phase. Then we compare the resulting spot-by-SNP count matrices of stCNASim-rs against seed and scReadSim-cna. Consistent with gene-level results, stCNASim-rs and scReadSim-cna produced matching SNP DP and zero-fraction statistics but sharply divergent phased AF quality. stCNASim-rs generated continuous, predictably shifted AF values across all CNA subgroups, while scReadSim-cna AF remained artificially polarized to 0 or 1. Linear regression fitted to phased SNP allelic fractions from normal spots quantified the discrepancy between simulated and ground-truth profiles: the stCNASim-rs versus seed comparison yielded $R^2 = 0.67$, while scReadSim-cna versus seed returned $R^2 = 0.04$.

Pseudobulk genome-wide phased AF tracks across all SNP datasets (Supplementary Fig. S18) reinforced these trends. StCNASim-rs tumor profiles exhibited distinct allelic distortion at simulated loss, gain, and LOH regions, matching the ground-truth CNA map (Fig. 1f), whereas scReadSim-cna pseudobulk AF tracks lacked any CNA-specific allelic stratification, rendering it unsuitable for benchmarking haplotype-aware CNA detection pipelines.
